# A combined morphometric and statistical approach to assess non-monotonicity in the developing mammary gland of rats in the CLARITY-BPA study

**DOI:** 10.1101/783019

**Authors:** Maël Montévil, Nicole Acevedo, Cheryl M. Schaeberle, Manushree Bharadwaj, Suzanne E. Fenton, Ana M. Soto

**Author notes:** Corresponding author: Ana M. Soto, Department of Immunology, Tufts University School of Medicine 136 Harrison Avenue, Boston, MA 02111. **Declaration of competing financial interests (CFI):** The authors declare they have no actual or potential competing financial interests.

## Abstract

**Background:** CLARITY-BPA is a rare collaboration of guideline-compliant (core) studies and academic hypothesis-based studies to assess the effects of bisphenol A (BPA).

**Objectives:** 1) determine BPA’s effects on the developing rat mammary gland using new quantitative and established semi-quantitative methods in two labs, 2) develop a software tool for semi-automatic evaluation of quantifiable aspects of the mammary ductal tree, and 3) compare those methods.

**Methods:** Sprague Dawley rats were exposed to BPA, vehicle, or positive control (ethinyl estradiol, EE2) by oral gavage beginning on gestational day 6 and continuing with direct dosing of the pups after birth. There were two studies; sub-chronic and chronic. The latter used two exposure regimes, one stopping at PND21 the other continuing until tissue harvest. Glands were harvested at multiple time points; whole mounts and histological specimens were analyzed blinded to treatment.

**Results:** The subchronic study’s semiquantitative analysis revealed no significant differences between control and BPA dose groups at PND21; whereas at PND90 there were significant differences between control and the lowest BPA dose and between control and the lowest EE2 dose in animals in estrus. Quantitative, automatized analysis of the chronic PND21 specimens displayed non-monotonic BPA effects with a breaking point between the 25 and 250μg/kg/day doses. This breaking point was confirmed by a global statistical analysis of chronic study animals at PND90 and 6 months analyzed by the quantitative method. The BPA response was different from the EE2 effect for many features.

**Conclusions:** Both the semiquantitative and the quantitative methods revealed non-monotonic effects of BPA. The quantitative unsupervised analysis used 91 measurements and produced the most striking non-monotonic dose-response curves. At all-time points, lower doses resulted in larger effects, consistent with the core study which revealed a significant increase of mammary adenocarcinoma incidence in the stop-dose animals at the lowest BPA dose tested.

## Introduction

The Consortium Linking Academic and Regulatory Insights on bisphenol-A (CLARITY-BPA) Toxicity is a collaboration between academic and federal government scientists, organized by the National Toxicology Program (NTP), the National Institute of Environmental Health Sciences (NIEHS), and the U.S. Food and Drug Administration (FDA) National Center for Toxicological Research (NCTR). This research consortium was “expected to significantly improve the interpretation of the wealth of data that is being generated by all consortium partners, including the characterization of the dose response of the effects observed and their interpretation in an integrated biological context” (Schug et al. 2013;Heindel et al. 2015).

The endocrine disruptor bisphenol-A (BPA) is widely employed in the manufacture of polycarbonate plastics and epoxy resins. It is present in various consumer products used on a daily basis (Vandenberg et al. 2013), such as thermal paper (Thayer et al. 2016;Hehn 2016). BPA had a > 90% detection rate in urine from samples representative of the US population suggesting that human exposure to the chemical is widespread (Calafat et al. 2005). BPA has also been detected in the blood of adults, and in the placenta, umbilical cord and fetal plasma indicating that the human fetus is exposed to BPA in the womb (Vandenberg et al. 2010;Gerona et al. 2013;Vandevoort et al. 2016). A large number of animal studies have revealed that exposure to environmentally relevant levels of BPA results in various deleterious effects including decreased fertility and fecundity, neuroanatomical, behavioral and metabolic alterations, obesity, and an increased propensity of developing prostate and mammary cancer (Soto et al. 2013;Acevedo et al. 2018;Cabaton et al. 2013;Diamanti-Kandarakis et al. 2009;Zoeller et al. 2012).

Regarding the etiology of breast cancer, exposure to estrogens during a woman’s lifetime has long been considered a main risk factor (Interagency Breast Cancer and the Environment Research Coordinating Committee.Department of Health and Human Services (IBCERCC) 2013;Kostopoulos et al. 2010). Developmental exposure (fetal and neonatal) to natural estrogens and estrogen mimics have long been proposed to increase the risk of developing breast cancer (Trichopoulos 1990). This hypothesis is backed by more recent data showing that iatrogenic exposure to DES as well as environmental exposure to the estrogenic pesticide DDT during fetal life increases the risk of developing breast cancer (Hoover et al. 2011;Palmer et al. 2002;Cohn et al. 2015). Likewise, the ubiquitous xenoestrogen BPA increased the propensity of developing mammary lesions in rodents (Murray et al. 2007;Acevedo et al. 2013;Durando et al. 2007;Lamartiniere et al. 2011;Jenkins et al. 2011). These data were gathered using different rat and mouse strains, different routes and timing of exposure, and different diets. Despite all these differences, the increased risk of effect attributed to BPA was consistent.

Our previous work on BPA-induced mammary gland carcinogenesis used the mouse model to address the effect of fetal and neonatal exposure on mammary gland morphogenesis (Vandenberg et al. 2007;Markey et al. 2001;Munoz de Toro et al. 2005;Wadia et al. 2013;Sonnenschein et al. 2011). The Tissue Organization Field Theory of carcinogenesis (TOFT) posits that carcinogenesis is akin to development gone awry (Soto and Sonnenschein 2011;Sonnenschein and Soto 2016). With this theoretical framework and previous data in mind, the main objective of the present study was to explore the effects of gestational and post-natal exposure to BPA on the morphogenesis of the rat mammary gland, as part of the CLARITY-BPA program.

While the mouse mammary gland is easily amenable to morphometric measurements from its earliest developmental stage to full maturity due to the flat, planar structure of the ductal tree, the rat mammary gland poses challenges due to the florid structure of the ductal tree that grows more conspicuously into the third dimension. This feature of the rat mammary gland hinders the application of conventional morphometric tools to the analysis of the rat mammary ductal system (Stanko et al. 2015). Hence, the second, subordinate objective of this work was to develop a proper software tool to perform computer driven, unsupervised analysis of the structure of the rat mammary ductal tree. The use of five BPA doses over a wide dose range, allowed us to explore whether the dose response curve to BPA is monotonic for the mammary gland end points examined in this study. A third objective of this study was to assess the non-monotonicity of the dose response using rigorous statistical tools. Our last objective was to provide a comparison between the semi-quantitative methods used to analyze the rat mammary gland (Davis and Fenton 2013) and the novel quantitative methods we are describing herein.

## Materials and Methods

### Experimental Design

This study was conducted as part of the CLARITY-BPA Consortium and we were provided with uniquely identified samples. The methods for this consortium have been published in detail (Delclos et al. 2014;Heindel et al. 2015;Churchwell et al. 2014) but are briefly described below.

#### Animals

The CLARITY-BPA studies used the NCTR-specific Sprague Dawley rat model and included 5 BPA doses, as well as a vehicle control and two doses of a positive reference estrogen control [ethinyl estradiol (EE2)]. Exposure by oral gavage to pregnant dams began on gestational day 6 and continued by direct dosing of the pups after birth. There were two exposure regimes, one stopping at PND21 and another whereby daily exposure continued until the time of sacrifice. All animal procedures for the subchronic and chronic exposure studies were approved by the NCTR Laboratory Animal Care and Use Committee and conducted in an Association for Assessment and Accreditation of Laboratory Animal Care (AALAC)-accredited facility, and in compliance with FDA Good Laboratory Practice (GLP) regulations. Sprague-Dawley rats (Strain Code 23), only available from the NCTR rodent breeding colony, were used in all experiments. Throughout the duration of the study, all animal rooms were kept at 23±3°C with the relative humidity of 50±20% under 12-hour light/dark cycles. Breeders were housed in polysulfone cages with hard chip bedding and glass water bottles (silicone stoppers) known to be free of contaminating BPA and provided food (soy- and alfalfa-free verified casein diet 10IF, 5K96, Purina Mills, Richmond, IN) and water for *ad libitum* consumption until weaning (approximately PND21). Resulting offspring were housed under the same study conditions from birth.

#### Reagents

BPA (CAS# 80-05-7, TCI America, Portland, OR, >99% pure) and EE2 (CAS # 57-63-6, Sigma-Aldrich Chemical Co., St. Louis, MO, 99% pure) were used in these studies (Delclos et al. 2014). The purity of BPA and EE2 were verified at 6-month intervals during the study and again at the end of the study to test article stability. The vehicle used to deliver BPA and EE2 was 0.3% aqueous carboxymethylcellulose (CMC; Sigma-Aldrich).

#### Dose Groups

Timed-pregnant rats that generated offspring used in these studies were dosed by gavage at a rate of 5 ml/kg body weight (bw) with vehicle control (0.3% CMC), BPA or EE2. <u>90-day subchronic study design (Pilot study):</u> four doses of BPA (2.5, 25, 260, 2700 µg/kg bw/day) and two doses of EE2 (0.5 and 5.0 µg/kg bw/day) were delivered from gestation day 6 until the initiation of parturition (PND 0). Starting on PND1, pups were directly gavaged with the same dose level of vehicle, BPA, or EE2 as their dams until the termination of the study. These treatment groups are referred to as Control, BPA2.5, BPA25, BPA260, BPA2700, EE2 0.5 and EE2 5.0. <u>Chronic study design:</u> five doses of BPA (2.5, 25, 250, 2500, and 25000 µg/kg bw/day) and two doses of EE2 (0.05 and 0.5 µg/kg bw/day) were delivered from gestational day 6 until the initiation of parturition. Starting on PND1, pups were directly gavaged for the period described below with the same dose level of vehicle, BPA, or EE2 as their dams. These treatment groups are referred to as Control, 2.5BPA, 25BPA, 250BPA, 2500BPA, 25000BPA, 0.05EE2 and 0.5EE2 from here on. Determination of BPA doses in the chronic study were based on 1) the results from the 90-day subchronic study (pilot), conducted by NCTR prior to the CLARITY-BPA chronic study (Delclos et al. 2014), 2) reported estimates of human exposure levels (Heindel et al. 2015), and 3) agreement among all CLARITY-BPA program stakeholders to focus the dose range for regulatory concern. All doses were administered at NCTR by daily gavage with a modified Hamilton Microlab 500 series programmable pump (Hamilton Co., Reno, NV; (Lewis et al. 2010)). Dosing was always conducted from the lowest to highest dose within a dosing pump, and cleaning and maintenance of the equipment were performed as described in (Delclos et al. 2014). The accuracy of dose delivery from the pumps was assessed every three months and established to be within 10% of the target volume accuracy. Litters were randomly culled to 3-5F:3-5M pups per litter on PND1. Direct gavage dosing of pups at the same dose level of vehicle, BPA, or EE2 as their dams started on PND1 (day of birth is PND0). Therefore, negligible lactational transfer of treatments was anticipated in this study (Doerge et al. 2010). Each dose group in the chronic study was split into two dosing arms, a continuous-dose (CD) group and a stop-dose (SD) group, with the latter having treatment terminated at weaning on PND21. Terminal body weight was assessed for all animals at time of sacrifice.

#### Tissue Collection

Offspring were euthanized on PND21 and 90 (both subchronic and chronic study), as well as at 6 months of age (chronic study only). One female per litter was necropsied (n=9-12 [subchronic] and n=8-10 [chronic] per treatment group per time point) for both Fenton and Soto lab evaluations. Samples from the chronic study were received from both the SD and CD arms of the study at 90 day and 6-month end points. In the chronic study, cycling females were euthanized when predicted to be in estrus based on a vaginal smear from the previous day, but that was not the case in the subchronic study. These latter females were necropsied at PND21 and PND90 (regardless of estrous stage). Estrous stage in PND90 animals was determined post-mortem based on vaginal histopathology, as determined by a NCTR pathologist.

The fourth inguinal mammary glands were collected per animal. One mammary gland was whole mounted to a charged glass slide and fixed in 70% ETOH, while the contralateral was placed in a cassette and fixed in 70% ETOH in a sealed plastic bag. The fixed mammary glands were shipped from NCTR to Tufts University School of Medicine. The whole mounted glands were stained with carmine and processed as previously described (Murray et al. 2007) and the contralateral glands were processed for paraffin embedding and histological sectioning. Figure 1 recapitulates the different animal sets used and their analyses. The following abbreviations are used to reference the study and exposure-dose groups in relation to the age of animal at time of tissue collection: PND21P and PND90P refer to the subchronic (pilot) study animal sets. SD and CD animals in the chronic study are referred to as PND21C, PND90SD, PND90CD, 6MSD and 6MCD.

**Figure 1:**
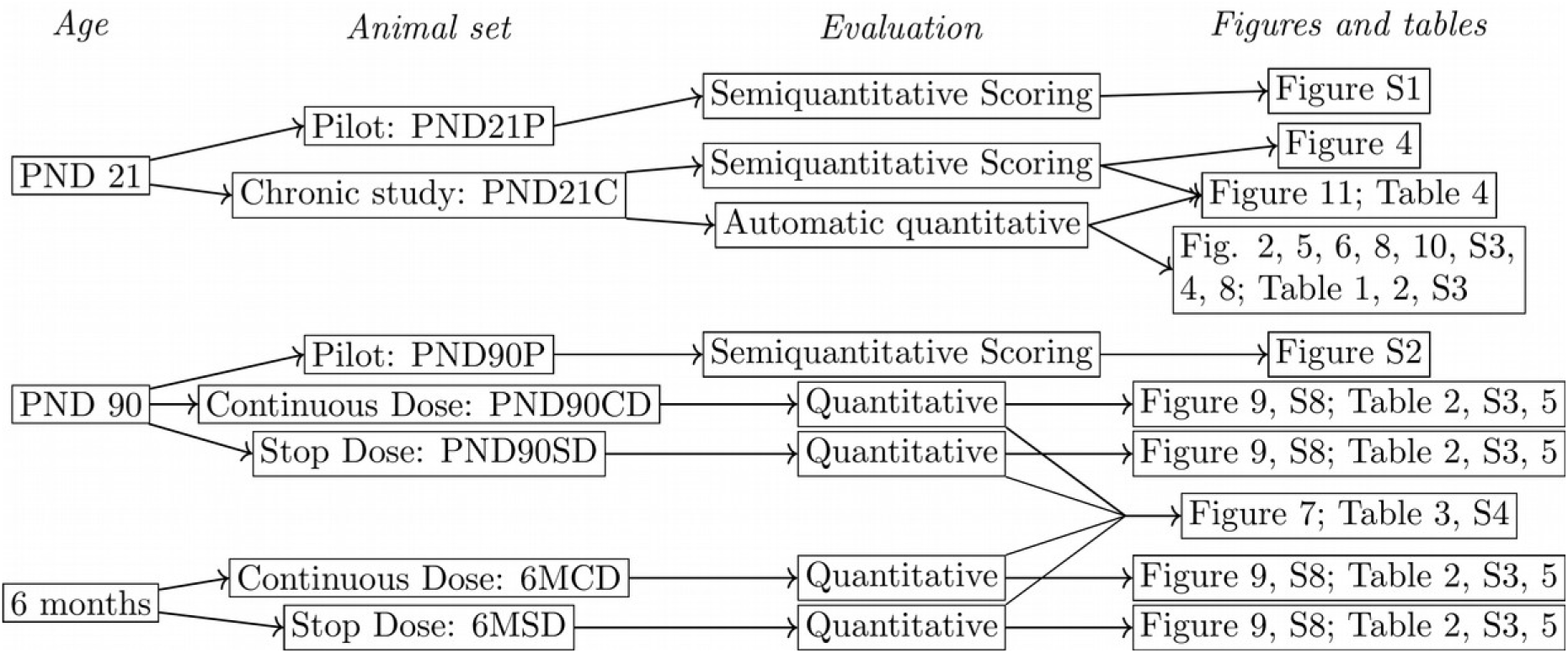
*Study design.* Each animal set contains a distinct group of animals and is therefore independent of the others. In the sub-chronic (Pilot; P) study, n=9-12 animals per group per end point and the groups are Control, BPA2.5, BPA25, BPA260, BPA2700, EE2 0.5 and EE2 5.0. In the chronic study, (C) n=8-10 animals per group per end point and the groups are Control, 2.5BPA, 25BPA, 250BPA, 2500BPA, 25000BPA, 0.05EE2 and 0.5EE2. Offspring were divided into continuous dose (CD) and stop-dose (at PND 21; SD) groups.

The whole mounts were evaluated by morphometric analysis using semi-quantitative methods (PND21P, PND90P, and PND21C), a semi-automatic morphometric method (PND21C), and a standard morphometric assay (PND90SD, PND90CD, 6MSD and 6MCD). Sections of the PND90 fixed mammary glands were used to assess the time-course of histoarchitectural changes and the emergence of preneoplastic and neoplastic lesions. All samples were received without knowledge of treatment group, and data were not decoded until data collection of histological and morphometric analyses were complete and the raw data were recorded in the NTP Chemical Effects in Biological Systems (CEBS) database.

### Mammary gland development scoring

#### Mammary gland scoring

In the subchronic BPA study (Delclos et al. 2014), the negative and positive control samples were identified *a priori* to the investigators and were evaluated to determine the range of response. A stereomicroscope (Nikon SMZ800, Nikon Instruments, Inc., Melville, NY) was used to develop the range of scores reported in Figure S1, which was modified for the range of responses in this study from the criteria reported by Davis and Fenton (Davis and Fenton 2013). A score of 7 represents a gland that is most well-developed, while a score of 1 suggests that few of the necessary developmental criteria are present. The scoring was adjusted to a 7-point scale, because there was a dramatic difference between control and high dose EE2 groups (5µg/kg bw/day; (Delclos et al. 2014)) in the subchronic BPA study. All PND21 whole mounted glands were given a morphological developmental score from 1 to 7 which considered 1) the number of terminal end buds (TEBs) relative to the number of duct ends, 2) the degree of ductal branching and/or ductal budding, 3) the number of primary ducts growing from the point of attachment, 4) the degree of lobule formation, and 5) the lateral and longitudinal growth of the gland (extension).

In a blinded manner, slides from PND21P and PND21C treated and control animals were evaluated using the scoring criteria summarized in Table S1. Stacks of slides were created for each score and all mammary glands within each score were reviewed a second and third time to ensure that the scores were assigned consistently over the course of the evaluation. Two individuals with knowledge of rat mammary morphology independently evaluated all slide sets. Disagreement in score of more than a full point for any sample required reconciliation between the two scorers.

The PND90P whole mounts were assessed using criteria described in Table S1, adjusted for age and stage of development. For instance, number of branch points, size and density were important contributors to assigned scores.

### Semi-quantitative mammary gland morphometric analysis

#### PND90 and 6-month mammary gland quantitative analysis

Mammary glands from PND90P, PND90CD, PND90SD, 6MCD and 6MSD animals were assessed for overall glandular development and density. Wet mammary gland weight was recorded at NCTR at time of collection. The chronic study glands were imaged with a Stemi 2000 stereomicroscope (Carl Zeiss) and Axiovision software (Carl Zeiss). Image J (NIH) software was used to process and analyze captured images to assess epithelial density of the gland (a measure of total fat pad area and epithelial area). Three standardized separate areas of each gland were measured to determine average density of the gland. Area 1 (rostral) was closer to the third mammary gland, area 2 was in the middle of the gland and area 3 (caudal) was closer to the fifth mammary gland. The size, and especially the thickness, of these whole mounts precluded a complete scan using a confocal microscope. Therefore, PND90 and 6-month glands were visually scored for the following countable morphological parameters: number of leading edge/internal terminal ends, as well as incidence of lateral branching, lateral budding, alveolar budding, and lobuloalveolar development. Putative lesions identified in whole mounts were excised for histopathological assessment.

### Semi-automated morphometric analysis of PND 21 mammary glands in chronic study

#### Imaging

In order to reduce ambiguity in the analysis due to overlapping branches, we obtained optical sections to generate a 3D image instead of a bright field image of the gland. This method was only applicable to PND21 mammary glands due to their smaller size and thickness compared to the later time points. Samples were imaged with a Zeiss LSM 510 confocal microscope using the auto-fluorescence of carmine as the signal. Due to the large size of the whole glands, the imaging was done on a grid, leading to 150 to 600 partially overlapping stacks. The resolution used was 5µm for the optical plane (x-y) and 3.5µm for the depth (z). The resulting stacks were stitched in Fiji using the method described in (Preibisch et al. 2009).

#### Identification of epithelium

Segmentation separates a region of interest from the background. In the mammary gland, the region of interest is the epithelium whereas the background includes the stroma, blood and lymph vessels. Currently available algorithms for reconstructing branching structures in the vascular system (Luboz et al. 2005) cannot be used for the mammary gland due to the presence of ductal buds (shown in Figure 2A). Therefore, we designed a custom semi-automatic method. Additionally, due to optical limitations, the presence of lumen did not provide a consistent pattern that could be used for segmentation. Because of this limitation, we found it easier to segment the stroma first instead of focusing directly on the epithelium. The segmentation algorithm used the following steps:

**Figure 2:**
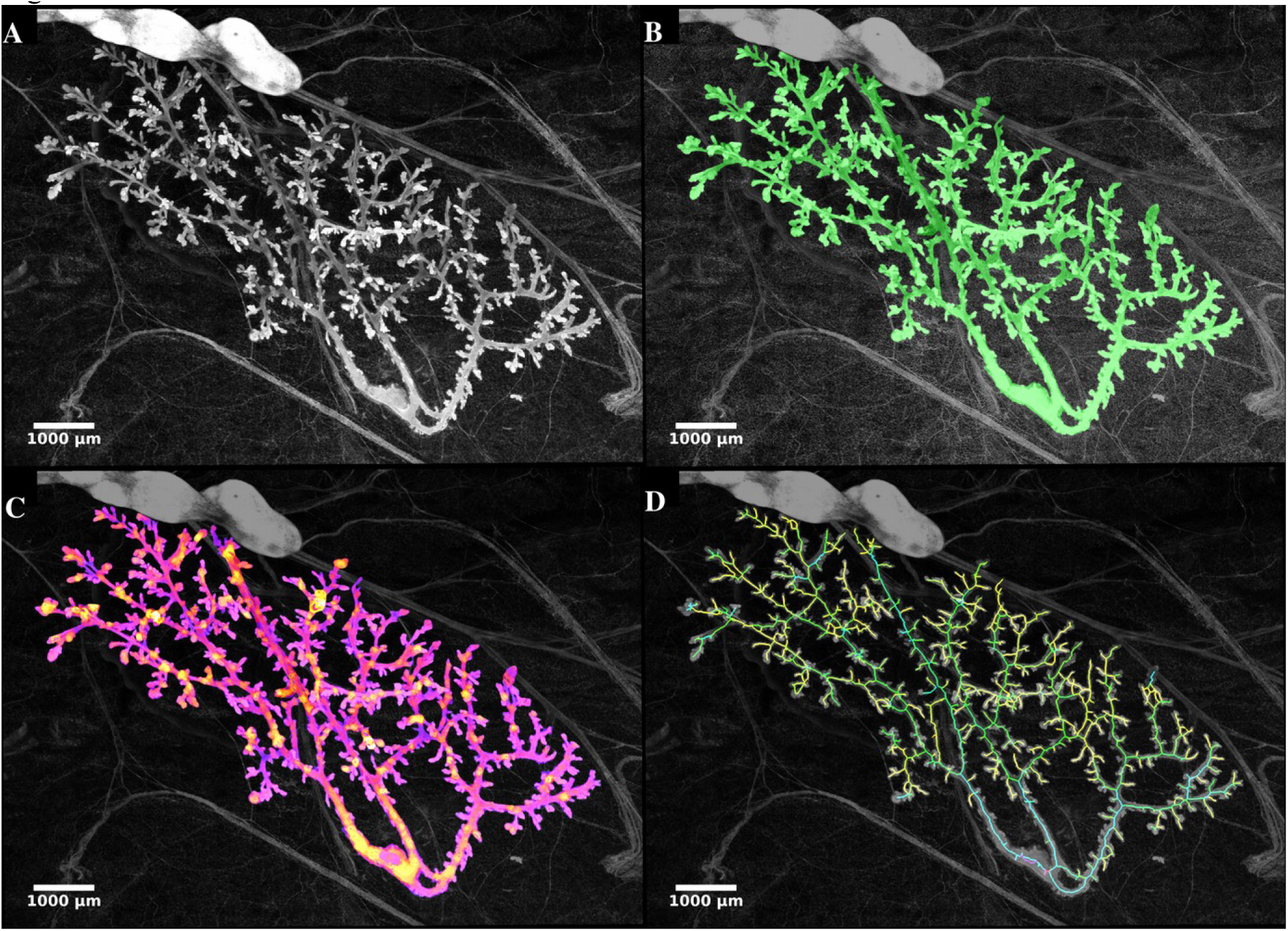
*PND 21 mammary gland from chronic study at different steps of analysis.* All images are projections and all data are processed as 3D stacks. (A) Original image after stitching. (B) Green overlay of the epithelium after segmentation (identification of the epithelium). (C) Analysis of the local thickness of the epithelium, warmer colors correspond to thicker parts of the epithelium in 3D. (D) Estimated skeleton of the epithelial tree, the color of a branch corresponds to the depth of the tree that starts at this branch. Scale bars are indicated.

Step 1: To remove nuclei of stroma cells and noise from image acquisition, we used bilateral filtering (with spatial radius 4 and range 150) followed by the subtraction of local background. The resulting image was then used for the segmentation.

Step 2: The image was inverted, and the stroma segmented as a bright connected region, with a uniform threshold. Then, 3D Gaussian blur (radius 2) was applied to the resulting binary image to remove small structures such as blood vessels, adipocytes, etc. Next, the image was inverted and the epithelium was obtained as the connected region above a given brightness which included a point in the epithelial tree that had been manually selected. Holes in the epithelium which were due to lumen were filled in and another Gaussian blur was performed. Finally, we performed a second selection of the connected region corresponding to the epithelium and above a given brightness. This second segmentation reduced possible artifacts which mostly stemmed from small blood vessels and adipocytes.

Step 3: Human intervention was required for comparing the segmented epithelium with the original image. The purpose of this comparison was, first, to assess whether all the epithelium was accurately segmented. Missing epithelium typically corresponds to a loss of brightness in deeper parts of the sample or particularly thin epithelial structures. Second, the user ensured that structures other than mammary epithelium were not segmented (such as blood, lymph vessels or lymph nodes). If the output was deemed acceptable, segmentation was complete; otherwise, human intervention was required to correct the segmentation issues. Intervention corrected the stacks that were used at the beginning of Step 2. Epithelial structures lost during segmentation were recovered by increasing the brightness. Non-epithelial structures were removed either completely or by decreasing the brightness around these structures. After this operation was performed, the program went back to Step 2 performing the segmentation and subsequent verification again.

#### Extraction of quantitative morphological features

The result of segmentation was a 3D reconstruction of the epithelium which is illustrated by the green layer of Figure 2B. This representation of the epithelium was then used to extract several quantitative morphological features of mammary glands. Analyses were performed using ImageJ. Before performing these analyses, we standardized automatically the orientation of the glands on the basis of the axes of inertia. We first analyzed the properties of the 3D epithelial reconstruction on the xy plane, which was comparable with assessments performed on bright field microscopy such as the developmental scoring. The analysis included quantities such as the aspect ratio (length/width), the epithelial area, and the fractal dimension of the epithelium in 2D (the projection of its 3D image). The same kind of global analysis was also performed in 3D and included an evaluation of the surface of the epithelium, of its volume, and of its 3D fractal dimension (based on the box counting method) (Longo and Montévil 2014).

Last, the analysis used several plugins from ImageJ: the 3D object counter (Bolte and Cordelières 2006), the plugin 3D shape (Sheets et al. 2013), and the bonej plugins (Doube et al. 2010). The latter included an evaluation of the local thickness (see Figure 2C), which was performed after the normalization of the scales of the three spatial dimensions. The epithelium was skeletonized, see Figure 2D, and this skeleton was analyzed by generic methods (counting the number of branches, average branch length, etc.). The analysis was performed both with and without terminal branches since some of the terminal branches may not have corresponded to actual epithelial structures but may have been artifacts from the process of skeletonization.

Finally, a more specialized approach to reconstructing the epithelial tree was performed by a custom plugin. This plugin started from the skeleton generated as discussed above and a manual selection of the starting point of the gland (the point of attachment). The plugin then reconstructed the mammary tree with the main duct as the root. To assess secondary branching, we performed the following operation recursively: for each branching, the distance (depth) of the two daughter trees was assessed; if the ratio between these depths was smaller than 0.3, then the branch associated with the smaller tree, A, was identified as a secondary branch and the other, B, was identified as a part of the parent duct. In this case, the parent branch and B were merged. This reconstruction was then used as the basis for evaluating various properties. For example, the distance from a branch to the point of attachment counted both in terms of the number of branching points and as the sum of the lengths of the branches that linked the two. We also considered branching angles and the tortuosity of the branches (i.e., for a branch, the ratio between its length by the length of a straight line between its extremities: the less straight the branch, the higher the tortuosity). Other quantities such as the local thickness were also determined by considering the average and standard deviation of their values on the skeleton points of every branch.

Overall our method assessed 91 structural features of mammary glands (shown in Table S2) complemented by three features: animal weight, mammary gland weight and manual assessment of the number of TEB.

### Statistical analysis

#### Rationale of the statistical analysis

In the semi-automated analysis, 6 dose groups (vehicle and 5 BPA doses) containing 10 animals each were used. More than 90 end points for each animal were measured to assess dose responses. This is quite different than the customary situation when one animal provides a much smaller number of end points, with the exception being certain types of -omics studies (transcriptomics-metabolomics). These situations (our quantitative measurements and the -omics) are more conducive to different types of analysis, such as principal component analysis, the permutation tests, etc. (Goh and Wong 2018). In addition, classical statistical tests such as Dunnett’s t and Student’s t lose power rapidly when the amounts of end points and doses increase. With the number of tests necessary for our quantitative experimental designs, these tests are not generally useful.

Another consideration is that there are neither theoretical nor empirical bases that predict a specific type of dose-response curve for BPA. Empirically, BPA dose-response curves could be monotonic for some end points and non-monotonic for other endpoints (Villar-Pazos et al. 2017); this is also the case with natural estrogens when comparing the effects on the uterus (monotonic) and the mammary gland (non-monotonic), using the same animal set (Vandenberg et al. 2006). Here, we consider morphological features which are the result of non-linear processes of morphogenesis at the tissue level *in vivo*, where many levels of organization are entangled. Thus, we decided that phenomenological curve-fitting was the best approach for this exploratory analysis.

Non-monotonic responses can take numerous forms. We focus on showing that a specific dose is the locus of a breaking point which is a specific kind of non-linear behavior and use the permutation test to assess this hypothesis. In our analysis, we distinguish exploratory and confirmatory statistics.

#### Principal component analysis

We performed principal component analysis (PCA) with R (R Development Core Team 2008) and the FactoMineR package (Lê et al. 2008). We use the dimdesc function of this package to assess the meaning of dimensions resulting from PCA and the effect of treatments; for further details see Supplemental Information.

#### Global analysis to identify a breaking point

To avoid errors stemming from multiple comparisons of non-independent variables, we designed an analysis at the level of the whole dataset. The responses detected were not U-shaped; instead they seem to be characterized by a sudden drop or a breaking point – these two patterns cannot be set apart on purely empirical bases. To explore the presence of a breaking point, we used the PND21C data set to formulate a precise statistical hypothesis and we used the four other animal sets of the chronic study (PND90CD, PND90SD, 6MCD, and 6MSD) together for a confirmatory analysis.

Each data set stemmed from a unique set of animals and were thus independent. However, the different features observed for an animal are not independent *a priori*. To accommodate this complex structure, we use the permutation test. Unlike traditional tests that use a standard distribution, the permutation test builds the statistic of the intended random variable on the basis of the data and the statistical hypothesis.

##### Defining the random variable X

We used the PND21C dataset to propose a variable *X* describing the presence of an overall breaking point between the consecutive treatments *C_b_*. *X* is large when *C_b_*is the locus of the largest change for most variables over the 4 remaining datasets.

More precisely, for each variable measured (see list in Table S2 for PND21C), we identified the consecutive concentrations where the difference was the largest. To avoid differences that stemmed from noise, we added an algorithmic criterion to check whether a difference was large enough to be included. We proposed two types of criteria:

The first series of criteria, *A(r_thr_)*, was met when the ratio between the mean values at consecutive conditions was larger than a threshold *r_thr_*. The larger *r_thr_*, the stricter this criterion became. For each variable, we looked for the consecutive concentrations with the largest *r_thr_* ratio.
The second series of criteria, *B(p_thr_)*, was met when a t-test between the consecutive conditions had a p-value that was smaller than a threshold *p_thr_*. Therefore, the smaller *p_thr_*, the stricter this criterion was. With this criterion, we compared the difference between the means of the consecutive conditions and looked for the largest difference meeting this criterion. Note, that we did not take *p_thr_*=0.05 because this condition was too strict. The aim of using the threshold was to disregard very small differences between consecutive conditions taking into account standard deviation, not to assess significance. The latter is done using the statistical test below.

We considered the variables:

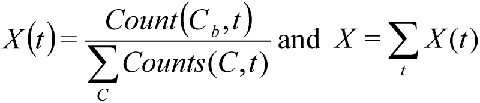

where *t* corresponds to one data set (PND90CD, PND90SD, 6MCD, 6MSD) and *C* corresponds to consecutive concentrations (Control-2.5BPA, 2.5-25BPA, etc.). The division, here, aimed to normalize the impact of the different data sets, so that they all contributed equally to *X*. As an illustration of the variables, assuming that there is no specific breaking point, since there were five groups of two consecutive concentrations, the mean of *X(t)* would be 1/5=0.2 for all data set *t* and the expectancy of *X* should be 0.2×4=0.8. This approximation is not used in our statistical analysis. Note that *X* summarizes the four datasets and does not enable us to draw conclusions for each dataset taken individually.

##### Statistical hypotheses

The null hypothesis H_0_ is that the treatment did not impact the rat mammary gland morphology, or in other words that all treatment conditions are equivalent. On the basis of the PND21C dataset, we formulate the alternative hypothesis H_1_: *X_observed_*is higher than in H_0_, meaning that there is a remarkable change at *C_b_*.

##### Statistical test

To assess whether our results were significant, we used the Monte Carlo permutation test (Nichols and Holmes 2001). Like most statistical methods, the permutation test assesses whether *X_observed_*is likely under the null hypothesis. The statistics of *X* is complex because the different variables describing a mammary gland are not independent. The permutation test provides an accurate solution to this problem. The permutation test provides an estimation *X_sim_*, of the distribution of *X* under the null hypothesis that treatment has no effect.

No effect of the treatment means that randomly shuffling the “exposure group” label in the data set yields an outcome that is equally probable as that of the initial data set. Therefore, performing several such permutations and computing the resulting value of *X* every time generates an estimation of the statistic of *X*. The operation of permutation is equivalent to randomly assigning the condition (exposure) of each animal but holding the number of animals for every condition constant and preserving all the measured biological properties of each individual animal. Since all variables describing individual animals except their condition (BPA exposure dose) are left unchanged, all correlations between the morphological features in data sets are preserved in the permutations. The permutation test requires that the order of the observations can be exchanged. It does not require independence of the different features observed for each animal.

This operation was iterated 10,000 times to obtain the distributions of *X_sim_(t)* for each of our 4 datasets. Then we added the values of *X_sim_(t)* to obtain the approximate distribution of *X*, *X_sim_*, under the null hypothesis. Since the datasets are independent, we computed *X_sim_* by an approximation of the convolution of the distributions *X_sim_(t)* to obtain a more precise approximation of *X.* Figure 3 shows the resulting distributions for criterion *A(1.2)* (Figure 3A) and criterion *B(0.5)* (Figure 3B) with *C_bc_=25BPA-250BPA*. 10,000 iterations and convolutions were sufficient to obtain smooth distribution in both cases.

**Figure 3:**
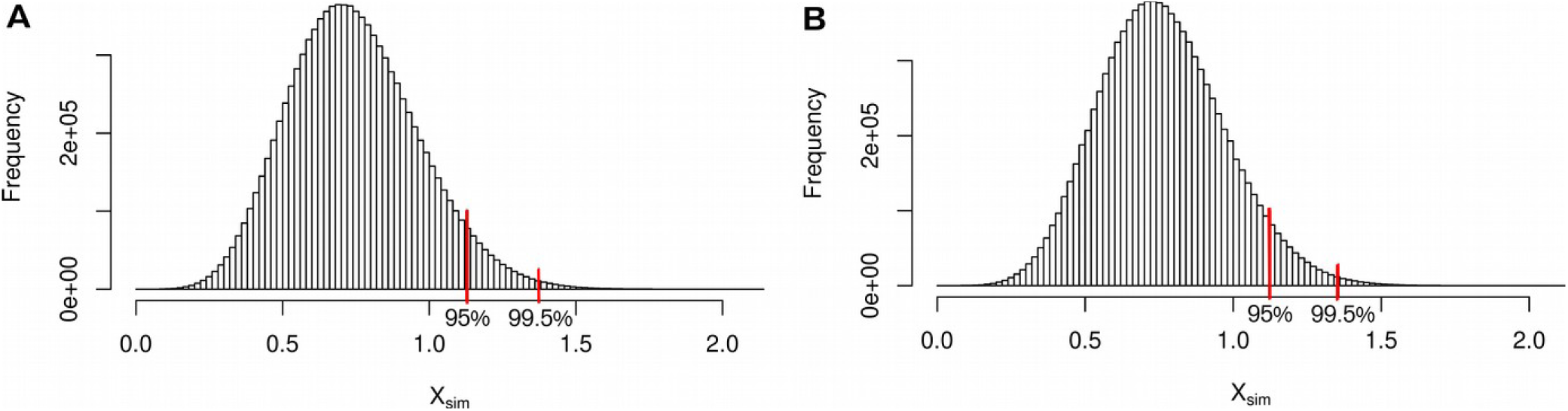
*Distribution of X_sim_ as a result of permutations of animal exposures. X* evaluates whether the change from 25 BPA to 250 BPA is different from changes between other consecutive concentrations in datasets from the chronic study. To evaluate the significance of results concerning *X,* we used the permutation test. The measurements performed in each animal were not rearranged; only the exposure labels were permutated. We computed *X* for 10,000 permutations and performed a convolution for the contribution of each dataset. We generated the distribution *X_sim_* which approximates the one of *X.* We marked the thresholds for values higher than 95% and 99.5% of the distribution of *X_sim_*. *X_observed_* above this threshold leads us to decide against the null hypothesis with p<0.05 and 0.005 respectively. (A) *X_sim_* for criterion *A(1.2)* with 10000 iterations: the ratio between consecutive values has to be at least 1.2 to be taken into account. (B) *X_sim_* for criterion *B(0.5)* with 10000 iterations: the p-value between consecutive values has to be at least 0.5 (t-test) to be taken into account. In both cases, we see that the simulation converges towards a smooth distribution. Number of animals per group n=8-10, number of groups: 6.

Next, we looked at the threshold (*X_thrs_*) for significance (p<0.05 and p<0.005) in the simulated distribution *X_sim_*. *X_thrs_* is such that 95% (0.995%) of *X_sim_* is smaller than *X_thrs_*. Since we did not perform multiple tests and the permutation number is high, our estimation of the p-value can be identified with the actual p-value (Phipson and Smyth 2010). Last, we compared *X_observed_*with *X_thrs_* to choose between the null or alternative hypothesis.

#### Mean comparisons and correlations

Developmental score is a synthetic quantity were the problems mentioned above do not apply, and its use corresponds to the assumption that BPA effects are similar to EE2 effects. ANOVA was used to assess the effect of BPA treatment on body weight and mammary gland weight, as well as the interaction of body weight or mammary gland weight with mammary developmental scores. The effect of BPA or EE2 on mammary developmental scores were analyzed by Kruskal Wallis non-parametric tests, and a Dunn’s post-hoc comparison of vehicle vs. treated glands (all at PND21, and at PND 90 from females in the estrus stage only, and in all stages not including diestrus and metestrus; GraphPad Prism, Version 7.05).

For other quantitative and unsupervised measures, we used the Student’s t-test to perform an exploratory comparison of means between control, 0.5EE2 and a dose singled out by the other analyses on PND21. The normality of the distributions was assessed by the Shapiro test. When this criterion was not met, we used a simple permutation test for the absolute difference in mean. We assess the false discovery rate due to multiple comparisons by the method described in and implemented in the LBE package for R (Dalmasso et al. 2005).

We also performed multiple comparisons between control and the other doses using Dunnett’s t test, controlling normality with the Shapiro test. To analyze correlations, we used Pearson’s product-moment correlation implemented in R (R Development Core Team 2008).

#### Regression

To perform regression, we used the linear model (lm function of R) on variables of interest. We compared the performance of the chosen model with simpler models (having fewer parameters) with a likelihood ratio test (using the lrtest function of R). We produced graphs for the qualitative assessment of normality and the distribution of residuals in supplementary materials. Since we are performing multiple regressions we use the LBE package to assess the false discovery rate.

## Results

We considered three hypotheses: (i) BPA effects were qualitatively similar to the effects of 0.5EE2, (ii) BPA impacted different features and/or had opposite effects of 0.5EE2, and (iii) BPA had no effect on mammary gland development.

### Developmental scoring of glands

#### PND 21 mammary gland development

Assessment of PND21P mammary gland development parameters showed that EE2 5.0 produced extensive ductal growth, twice the average developmental score of vehicle controls (Figure S1A). These weanling mammary ductal trees displayed developmental characteristics akin to adult mammary glands (Figure S1D). Therefore, this dose was deemed inappropriate as a positive control and doses were reduced to 0.5 and 0.05 for the chronic study. Although the effects of EE2 (at both 0.5 and 5.0) were significant, there were no statistically significant effects of BPA on the PND21P mammary glands in the subchronic study (Figure S1A).

In the chronic study, BPA and EE2 exposures had no significant effect on the body weight of female weanlings, nor on the weight of the excised mammary fat pad. Scoring of PND21C mammary gland morphology revealed that only treatment with 0.5EE2 resulted in significantly different glandular development compared to vehicle control (p=0.001) (Figure 4). There was a significant correlation between body weight and developmental score (p=0.009), as well as mammary tissue weight and developmental score (p=0.04). However, there was not a single dose group driving those effects (as evidenced in Figure 4).

**Figure 4:**
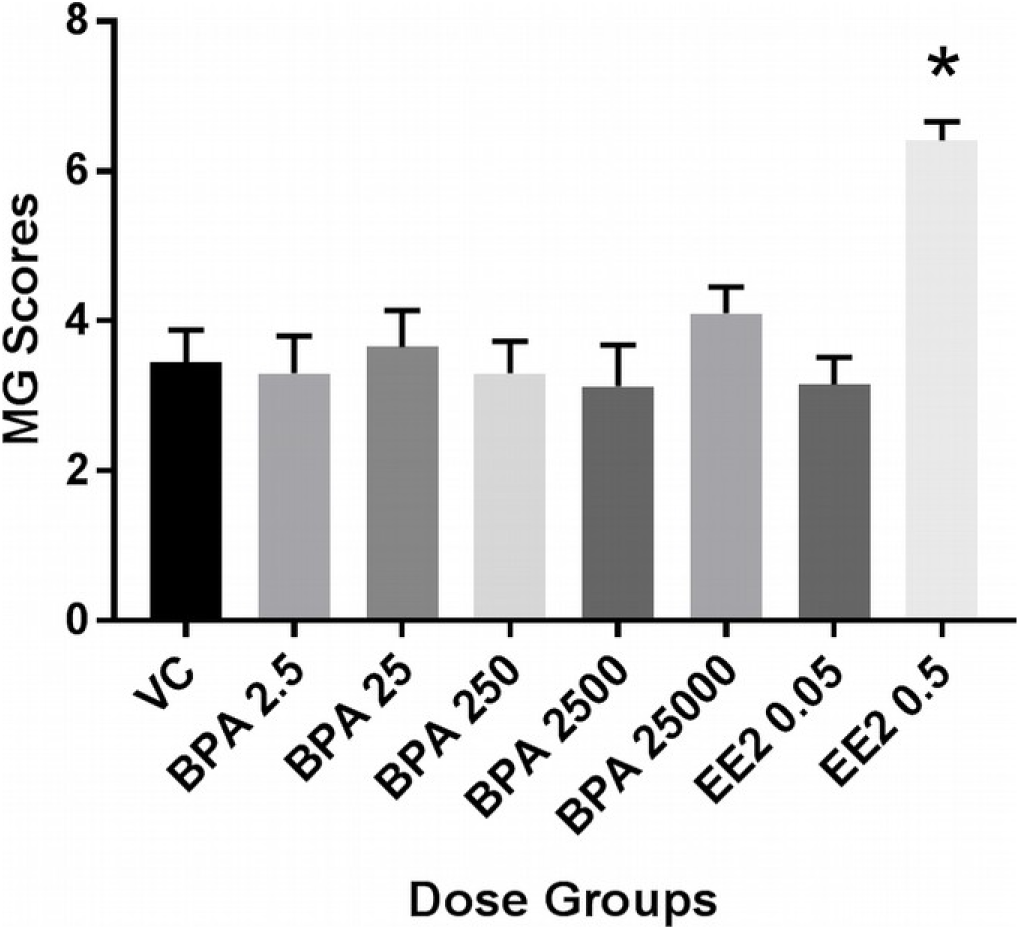
PND21 Mammary gland development across all exposure groups in the chronic study (Fenton group). Based on data from the pilot 90-day subchronic study, glands were scored on a 7-point scale, where a score of one (1) relates to poor development and score of seven (7) relates to accelerated growth and development for this age group. Values shown are mean +/-SEM for n=8 (2500BPA only) or 10 (all others) females per dose group. *significant at p=0.001 (Kruskal Wallis, Dunn’s post-hoc).

#### PND 90 and 6-month mammary gland development

Mammary whole mounts of PND90P animals were assessed for developmental scoring. Females in the subchronic study were not necropsied at a predetermined stage of the estrous cycle. Analyses of developmental scores accounted for dose group and vaginal pathology-based cycle stage, and those data demonstrated significant estrous stage × score interaction (p=0.05), and significantly accelerated gland development in the BPA2.5 and EE2 5.0 animals, compared to vehicle controls, when evaluated in estrus (Figure S2A). The EE2 0.5 and BPA25 group was also advanced in development but did not reach significance. No developmental effect of treatment (BPA or EE2) was seen when all cycle stages were considered within a group or across groups of glands. The stage of the estrous cycle by itself appeared to affect the morphological outcomes, thereby validating the importance of assessing all the tissues at the same stage of the cycle for the determination of treatment effect.

#### Global analysis: 25BPA-250BPA as a breaking point

**Exploratory Principal Components Analysis (PCA) on PND21C computer assisted morphological measurements:**

PCA provided an overview of the 91 structural features (shown in Table S2) assessed in PN21SD mammary glands through our computational analyses, plus three additional features assessed separately (body and mammary weights and TEB number). Dimension (Dim) 1 represented the size of the gland and its highest correlation was with the number of branches and the surface area of the epithelium in 3D [correlation coefficient (CC):0.97; p<e-49]. Dim 1 separates 0.5EE2 glands from the other exposure groups (t-test: p=6.4e-09). Dim 2 is related to the thickness of ducts in 3D and is more highly correlated with the average thickness of ducts (CC: 0.84; p=4.1e-22). Dim 2 also separates 0.5EE2-exposed glands (p=0.031, Figure 5A). Dim 3 corresponded to mean duct length (CC: 0.75; p=2.9e-14) and separates 250BPA-exposed group from the others (p=0.038) (Figure 5B). For additional details see Supplemental Material on PCA analysis and Figure S3 and S4. Dim 3 is the first dimension with significant differences for BPA exposure and seems to follow a non-monotonic pattern with a drop in the response between 25 and 250 BPA (Figure 5B). In the following section we assess whether this pattern is real, provided that it could be non-significant or an artifact from PCA.

**Figure 5:**
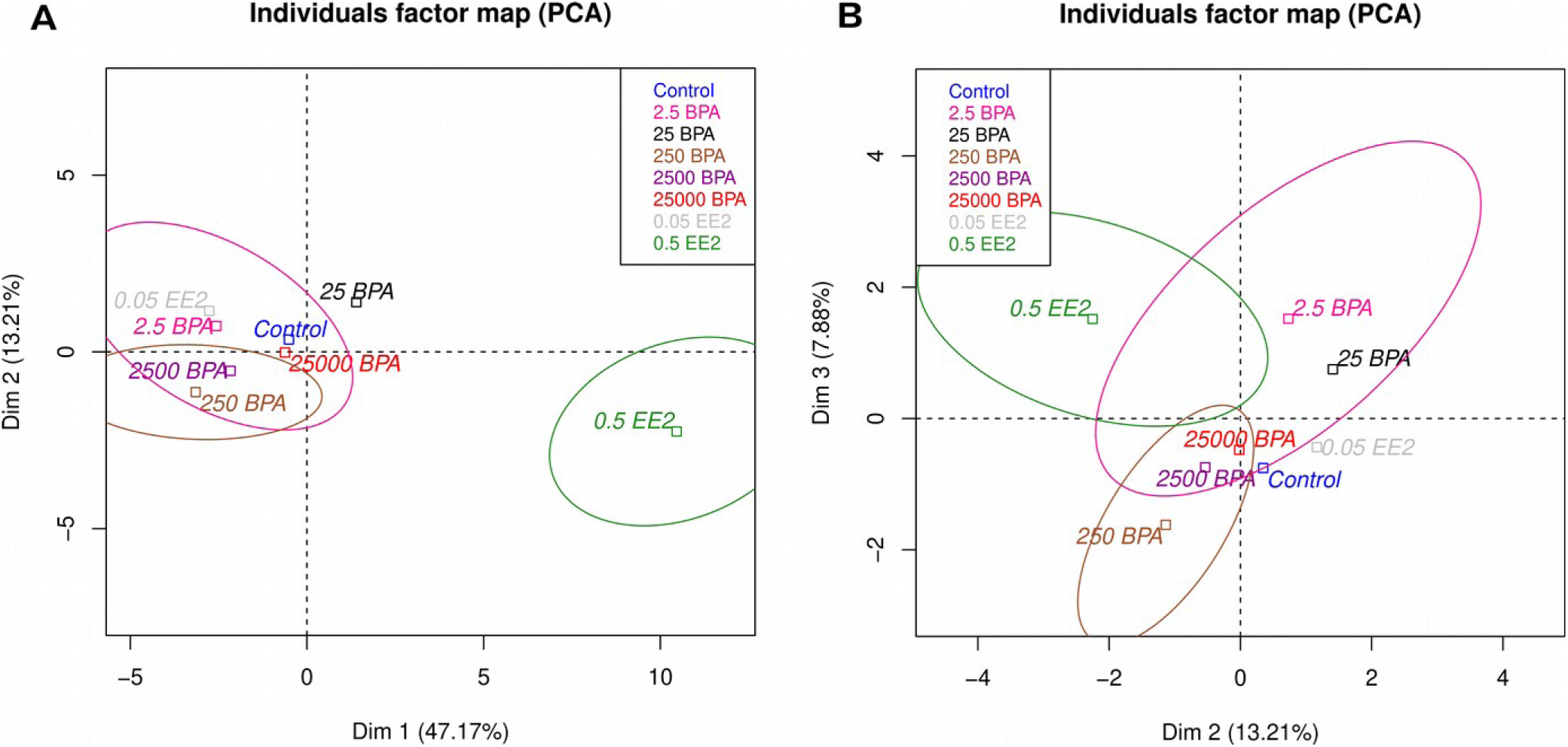
*PCA of data obtained by the quantitative analysis of PND21C animals.* In all cases, we represent only the mean of each exposure group. A: PCA revealed that the high dose of EE2 (0.5 µg/kg/day) produced a strong change in mammary gland morphology illustrated by Dim 1 (∼size) and 2 (∼local thickness). B: At the level of Dim 2 (∼local thickness) and Dim 3 (∼ductal length) (B), the response seems non-monotonic. 2.5BPA and 25BPA are close, while 250BPA is very different from 25BPA and control and high BPA doses are roughly between 25BPA and 250BPA. Ellipses show some confidence intervals, and the variability of the data remains high. Number of animals per group n=8-10.

PCA provides clues against hypothesis (i), BPA effects are similar to EE2, and for (ii), BPA treatment is associated with different morphological changes than that of 0.5EE2. Moreover, PCA shows significant differences between BPA treatments and vehicle which suggests that the hypothesis that BPA has no effect (iii) does not hold.

### Hypothesis formulation on the basis of PND21C datasets

We used our PND21C results as the basis to formulate our statistical hypothesis, and we used the four other independent datasets (PND90CD, PND90SD, 6MCD, 6MSD) to test this hypothesis. As detailed in methods, a simple way to formulate our question was to look at every feature we measured and for each of them to assess which consecutive concentrations have the largest difference. We used criteria to define differences and decide whether they were sufficient to be used. With the criterion *A(r_thr_),* we compared the ratio between means (met when the ratio between the consecutive conditions was larger than *r_thr_*), and, with criterion *B(p_thr_)*, we considered the arithmetic differences (met when a t-test between the consecutive conditions had a p-value that is smaller than the threshold).

In PND21C, with criterion *B(p_thr_)* the consecutive concentrations of 25 - 250BPA are associated with the largest number of changes in morphology (Table 1). The different thresholds *p_thr_* and criterion *A(r_thr_)* provide similar results. On the basis of this result and the discussion above, we hypothesize that the consecutive concentrations 25-250BPA is the locus of the largest change for most variables.

**Table 1.** Number of observed quantities where the largest difference is between each consecutive condition in the PND21C data set.

| $p_{thr}$ | Control – 2.5BPA | 2.5 – 25BPA | 25 - 250BPA | 250 - 2500BPA | 2500-25000BPA |
| --- | --- | --- | --- | --- | --- |
| 0.05 | 3 | 0 | 17 | 0 | 0 |
| 0.5 | 14 | 5 | 57 | 6 | 9 |
| 1 (no threshold) | 15 | 6 | 52 | 7 | 9 |

Note: Differences are counted when the significance of this difference has a $p$ value lower than $p_{thr}$ for a t-test (criterion $B(p_{thr})$ ). For all values of $p_{thr}$ , the interval 25-250BPA is the one that shows the largest value.

For a given data set *t* and a given criterion, *X(t)* is the proportion of quantities whose largest consecutive difference meeting the criterion is between 25BPA and 250BPA, for example for dataset PND21C and criterion *B (0.5)*, 57/(14+5+57+6+9)=0.62 (based on data in Table 1). Figure 6 illustrates the PND21C dataset and the result of permutations. If we assume that BPA has no effect, then the observed number of features having their strongest difference between 25 and 250 is unlikely to occur. *X_observed_*is the sum of *X(t)* over all data sets (PND21C only for the exploratory analysis and all other datasets for the confirmatory analysis below), see Table 2. Our statistical hypothesis *H_0_*is that the treatment does not impact *X*, and the alternative hypothesis *H_1_* is that *X_observed_* is higher than under *H_0_*. To assess whether we should reject *H_0_*in favor of *H_1_*, we used the permutation test on the PND21C data set. The test yields p=0.0094 for criterion *B (0.5).* This exploratory analysis suggests that we should reject *H_0_* for *H_1_*.

**Figure 6:**
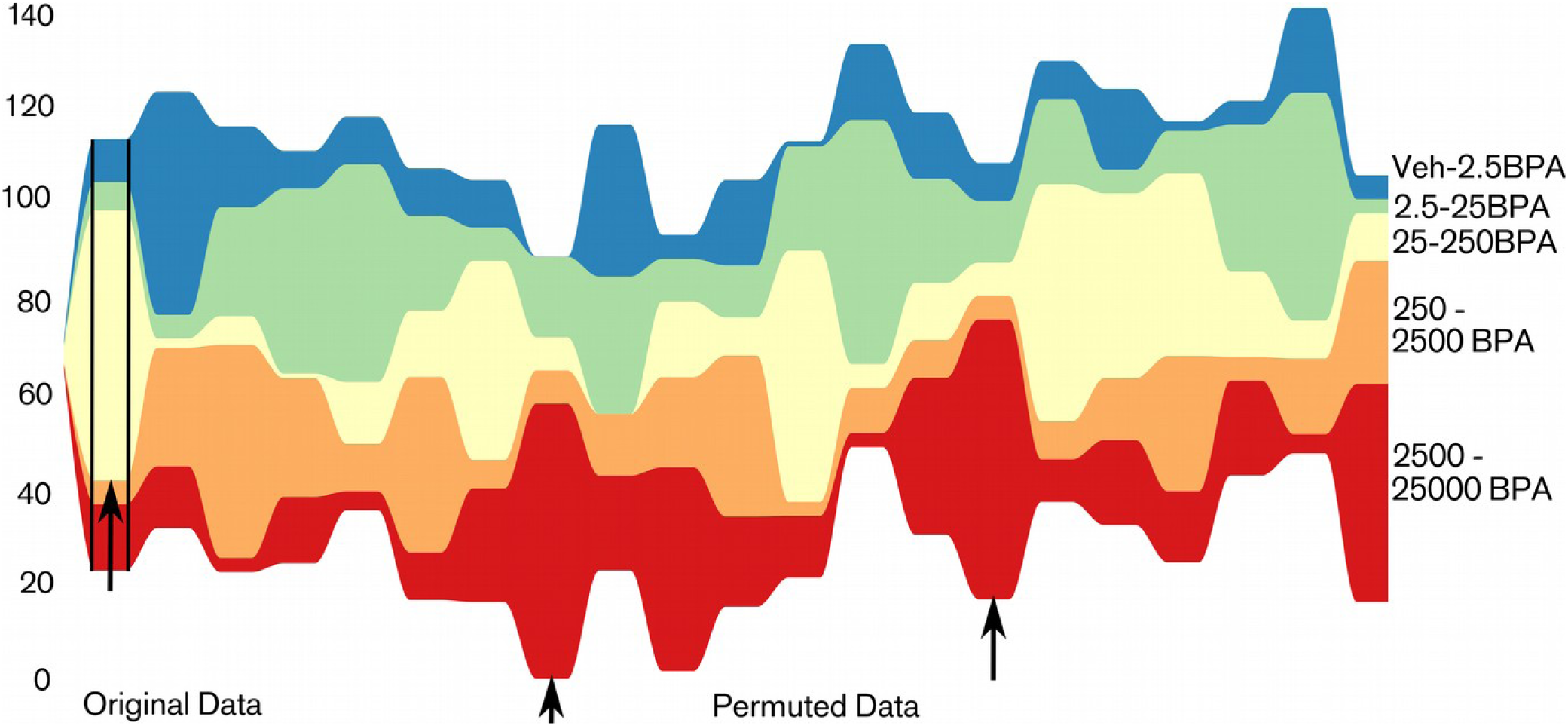
*Streamgraph of the number of quantities that have their largest difference between one of the 5 possible consecutive doses in PND21C.* Only differences meeting criterion B (0.5) are taken into account. We represent the PND21C original data (left) and 20 datasets obtained by random permutations of the “doses” in this original dataset (right of the second black vertical line). In the original data, the consecutive doses 25-250BPA is the location of the largest number of differences by far (left black arrow). In sets with randomly permuted doses, such an extreme situation is relatively rare but nevertheless happens occasionally. This is due to the correlations between the different variables that are preserved in permuted data (for example correlation between the area of a gland and its volume): except for the dose, the properties of individual gland described by many variables remain unchanged in permutations. These correlations tend to increase the probability that several variables will behave in the same manner, and therefore the frequency of large deviations (other black arrows). We use this result as a “pilot” to state our statistical hypotheses. H_0_: BPA exposure has no effect and H_1_: 25-250BPA is the location of the largest change for a larger number of variables than in H_0_.

**Table 2.** Number of features for which the largest change takes place between consecutive conditions in the chronic study.

| Dataset | Control-2.5BPA | 2.5BPA – 25BPA | 25BPA-250BPA | 250BPA - 2500BPA | 2500BPA - 25000BPA | X(t) | p |
| --- | --- | --- | --- | --- | --- | --- | --- |
| PND21C (training set) | 14 | 5 | 57 | 6 | 9 | 0.62 | 0.0094 |
| PND90CD continuous | 2 | 7 | 6 | 6 | 1 | 0.27 | 0.0038 |
| PND90SD | 3.5 | 3.5 | 7 | 1 | 3 | 0.39 |  |
| 6MCD | 5 | 3 | 9.5 | 0 | 6.5 | 0.40 |  |
| 6MSD | 3.5 | 8 | 7.5 | 2 | 3 | 0.31 |  |

Note: For criterion $B$ with $p_{thr}=0.5$ , we find $X_{observed}=0.27+0.39+0.40+0.31=1.37$ which is higher than the estimated expectancy of 0.8 and $X(t)$ is higher than the estimated expectancy of 0.2 for all datasets. Table 3 reports the significance of these results. Non integer values stem from ties.

#### Confirmatory analysis with PND90CD, PND90SD, 6MCD and 6MSD datasets

We used the remaining, independent datasets PND90CD, PND90SD, 6MCD and 6MSD for a confirmatory analysis. We found that using criteria *A(r_thr_)* with reasonable values of *r_thr_*(between 1 and 1.5), *X_observed_* was significantly higher than mean (*X_sim_*) for all significance criteria we have chosen (p<0.005), see Table 3 and S4. Similarly, using criteria *B(p_thr_)* with *p_thr_* between 0.4 and 1, *X_observed_* was significantly higher than the mean of *X_sim_* (p<0.05) with a loss of significance when the threshold was too strict or not strict enough. In particular, *B (0.5)* is the best compromise between type 1 and type 2 error rates in simulations (Figure S5, S6, S7) and it leads to p=0.0038<0.005. Figure 7 illustrates this result, and the fact that 25-250 is remarkable in comparison to the other consecutive doses. We could then safely reject the null hypothesis *H_0_* and adopt the alternative *H_1_*: the treatment led to a higher *X_observed_* than if BPA did not have an effect. 25-250BPA is the locus of a “jump” in the dose response. A remarkably high number of variables had their largest change between 25BPA and 250BPA, and this interval was the locus of a modified response to BPA.

**Figure 7:**
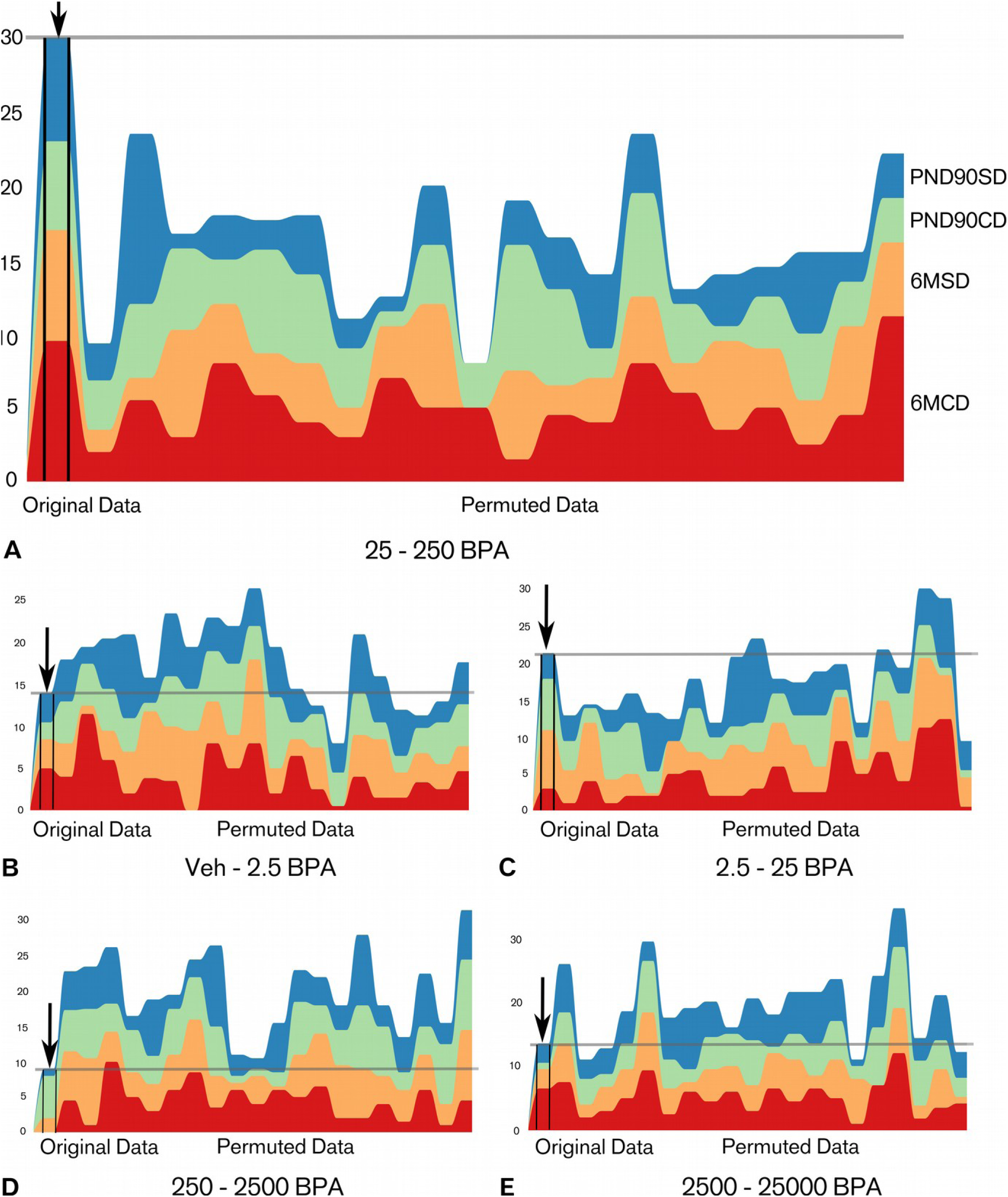
*Number of quantities that have their largest difference between the target consecutive concentration 25 and 250BPA (A) and the others (B, C, D, E) for the datasets PND90SD, PND90CD, 6MSD and 6MCD.* In each graph: left, original data, right 20 datasets obtained by random permutation of the “condition” of each animal. A: consecutive concentration 25-250BPA. The figure shows that the original data are remarkable: the sum of these quantities (grey horizontal line) is higher with the original dataset than with each one of the 20 permuted datasets. This representation suggests that the result is significant with p <=0.05, which is shown by more extensive simulations (Table 3 and S4). B, C, D, E: Consecutive concentrations Veh – 2.5BPA, 2.5 - 25BPA, 250 - 2500BPA, 2500 - 25000BPA respectively. In B, D, E the sum of the number of quantities in the original datasets (grey horizontal line) is below most of the permuted data which is expected since the consecutive concentration 25 - 250BPA is the locus of the largest change for many variables. In C, 2.5 - 25BPA, this effect is not as strong.

**Table 3.** Comparison of X_observed_ and the results of the permutation test show consistently that a significant, remarkable change occurs between 25 and 250BPA in PND90CD, PND90SD, 6MCD and 6MSD datasets analyzed together.

| Criterion | $X_{observed}$ | 95 % of $X_{sim}<$ | 99.5% of $X_{sim}<$ | $P_{estimated}$ |
| --- | --- | --- | --- | --- |
| A(1.2) | 1.67 | 1.13 | 1.38 | 0.00016 |
| B(0.5) | 1.37 | 1.12 | 1.35 | 0.0038 |

Note: $X_{observed}$ is significantly higher than the mean of the simulations. Results for other thresholds can be found in Table S4. Criterion A(1.2) [B(0.5)] means that the ratio [p value of a t test, resp.] between successive means has to be at least 1.2 [0.5] to be taken into account.

To conclude this global analysis, it is noteworthy that the semi-quantitative scoring did, in fact, display a non-monotonic response in morphological development with a slight breaking point between 25 and 250BPA-exposed glands in PND21C (Figure 4) and a more pronounced one in PND21P (Figure S1) and PND90P (Figure S2). PND21P and PND90P are animal sets that were not used in the global analysis, thus the fact that they reproduce the same pattern qualitatively is compelling.

### Exploratory analysis of 25BPA-250BPA as a breaking point and comparison with the effect of EE2

#### Assessing non-monotonicity

##### In PND21C

A non-monotonic response is characterized by a change in the trend of the response. Non-monotonicity was observed in various measurements obtained from PND21 mammary glands. One end point was the *mean variation of ductal thickness* (sd width 3D) which describes whether structures are more tubular or, the opposite, irregular. This measurement is associated with budding since small buds are not recognized as individual structures and lead instead to duct width variations in the automatic analysis. This quantity increased between control and 25BPA, dropped, and then increased again between 250BPA and 25000BPA (Figure 8A).

**Figure 8:**
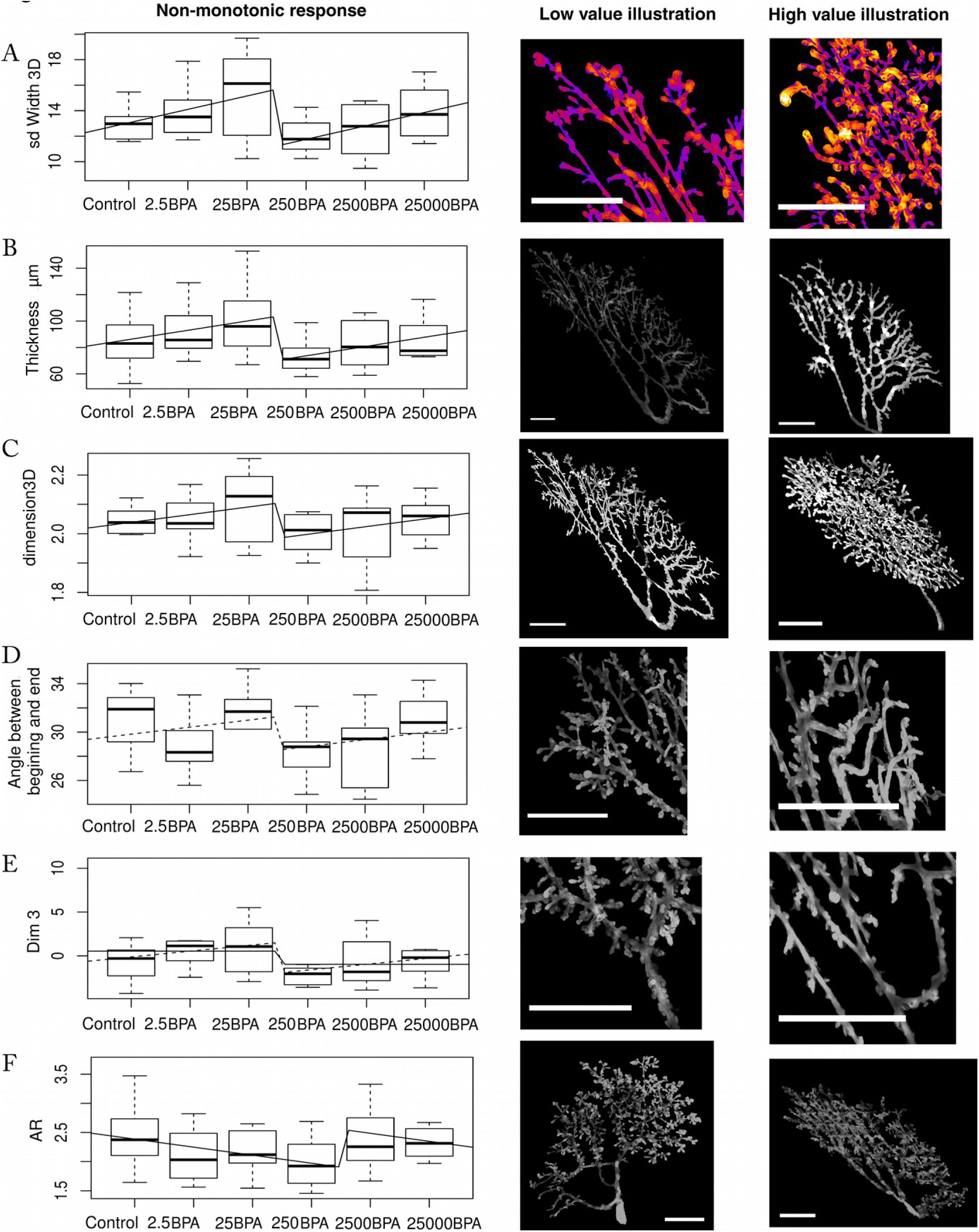
*Non-monotonic responses to BPA doses (x-axis, BPA μg/kg bw/day) shown by non-linear regression in PND21C animals and illustration of the corresponding morphological features.* Number of animals per group n=8-10. Graphs on the left represent mean and standard deviation for each dose and the fit with the combination of a linear and step function. Left panels are representative images of a low value, right panels illustrate high values. Scale bars = 2mm. All features but the aspect ratio (F) show a break between 25BPA and 250BPA. In (F) the break is between 250BPA and 2500BPA. (A) *Mean variation of ductal thickness:* the gland on the right has many structures that have both thin and thick parts while the gland on the left has more regular structures. (B)*Mean thickness of the epithelium:* the brightness in the pictures is proportional to the local thickness of the points of the gland. (C) *Fractal dimension in 3D*. The gland on the right grows more conspicuously in the third dimension than in the left figure. (D) *Angle between the beginning and the end of ducts:* ducts are straighter on the left and turn more on the right. (E) *Third dimension from PCA*. (F) *Aspect ratio (AR)*. A large AR leads to an elongated gland while a low AR means that the gland is round. Low doses of BPA increase the roundness of glands and high doses lead to an aspect ratio similar to control.

The responses detected seemed to be characterized by a sudden drop or even a breaking point, which implies two changes of trend. The model chosen for describing these data was the sum of a linear response and a step function which models a breaking point:

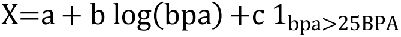

where *a*, *b* and *c* were found by linear regression. *b* represents a linear trend whereas *c* quantifies the sudden change between 25 and 250BPA. If *b* and *c* have opposite signs, the change between 25 and 250BPA breaks monotonicity. Note that we are not interested in the significance of *a* because *a* being different from 0 is not important in our analysis.

At low doses, our model describes a linear response for the considered variable (0-25BPA). Then, it leads to a drop in the response that is triggered at a critical concentration that we identified by maximum likelihood. In most cases, this negative effect started around 25BPA and reached a maximum around 250BPA. Beginning at 250BPA, increasing BPA levels resulted in a linear response. We emphasize the significance of the non-linearity of the response that took place between 25 and 250BPA (Figure 8); however, we do not make strong claims on the equational form of the model. Our model is what is usually called a phenomenological or a heuristic model; it reproduces the trends of the dose response of several endpoints. The choice of a specific model may only be decided by a theoretical discussion (Montévil 2018).

Since we are performing exploratory statistics on all features observed, we assess the false discovery rate for the comparison with a constant model noted *q*, with a threshold of 0.1. We do not report all features displaying these patterns, instead we report features with distinct biological meaning.

For sd width 3D, the non-monotonic model leads to significant fit (p=0.0039, 0.00038 for *b* and *c* respectively). This model is significantly better than a constant model (LR test, p=0.0011, q=0.020), a linear model (p=0.00025) or a step function (p=0.0029) (see Figure 8A). This model captures two changes of trend because *b*>0 and *c*<0 whereas a quadratic model can only fit one. Therefore, a quadratic model did not fit the data.

Figure 8B represents the *average of the local thickness*. At a point, the local thickness is the radius of the biggest sphere that contains this point and that is contained in the structure. The model’s fit of the data is significant (p=0.023, 0.0017 for *b* and *c* respectively), and the model is better than a constant model (LR test, p=0.0029 and q=0.031), a linear model and a step function (LR test, p=0.0012, 0.019 respectively).

Figure 8C represents the *fractal dimension in 3D* by the box counting method. The higher this quantity is, the more the epithelium is filling the stroma of the gland in 3D. This quantity is an assessment of the complexity of the gland. The fit is significant (p=0.062, 0.011 for *b* and *c* respectively), and the model is significantly better than a constant (p=0.024, q=0.073) and a linear model (p=0.0086), and almost better than the step model (p=0.054). The step model alone is not a better fit (p=0.057).

Figure 8D represents the *average angle between the beginning and the end of ducts*. This quantity assesses how much the ducts change direction during their growth. The fit is not entirely significant with p=0.17 for *b* and p=0.047 for *c*, and it is not significantly better than a constant model and a step model, only than a linear model (p=0.090, 0.16 and 0. 041 respectively). Nevertheless this feature is biologically interesting, and we will discuss it again below.

Figure 8E represents the third dimension constructed by PCA, which is associated with duct length. The linear part of the model is not significant (p=0.11, 0.018 for *b* and *c*). Nevertheless it is better than a constant model and a linear model, but not a step model (p= 0.031, 0.015, 0.097 respectively, q=0.078). The latter is a good fit (p=0.045), and is better than a constant model (p=0.041).

Figure 8F represents the *aspect ratio* (AR). This quantity is the ratio between the largest axis of the gland and its smaller axis and is one aspect of how the epithelium invades the fat pad. For 250- 2500BPA instead of 25-250BPA, our model is a good fit (p=0.032, 0.0092 for *b* and *c* respectively), and is better than a constant, linear and step model (p=0.027, 0.0072, 0.027 respectively, q=0.073).

The existence of various end points exhibiting a non-monotonic response is against hypothesis (iii) which postulates that BPA is devoid of effect. The conclusion of this exploratory analysis is that non-monotonicity can be different than quadratic (U shaped) responses, and that features related to thickness, duct width, fractal dimension in 3D are of interest. The aspect ratio has also an interesting pattern, but not with respect to the 25-250BPA breaking point.

#### In PND 90 and 6-month

Non-monotonic responses in PND90CD, PND90SD, 6MCD and 6MSD were found which are similar to the ones in PND21C (Figure 9). More specifically, the gland weight (determined at necropsy) in PND90SD (Figure 9A) is a significant fit to our model (p=0.039 and 0.039 for *b* and *c* respectively). The model is not significantly better than a constant model; nevertheless, the p value is below 0.1 (p=0.088). It is significantly better than a linear or a step model (p=0.033 and 0.033 respectively).

**Figure 9:**
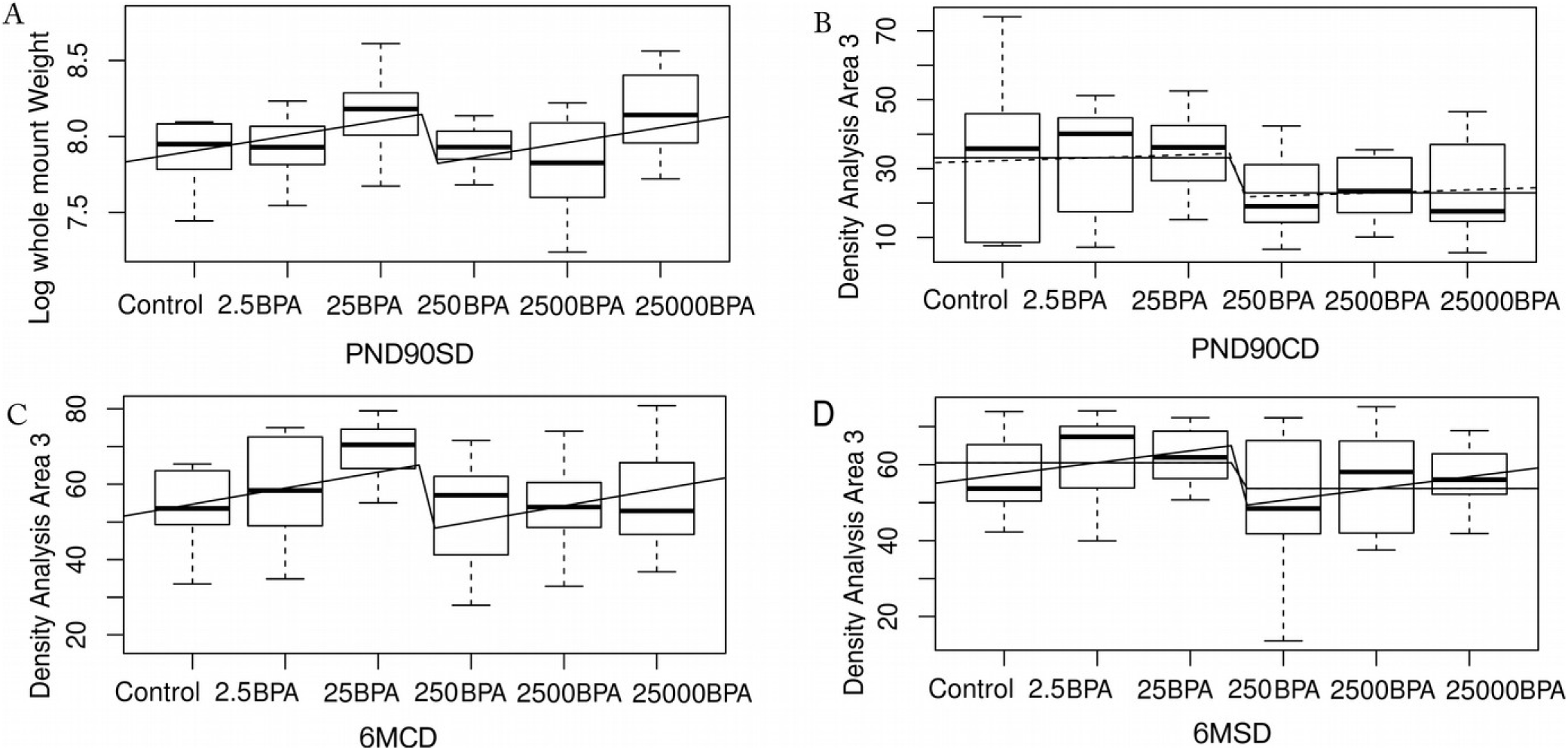
*Non-monotonic mammary epithelial responses in glands from BPA-exposed PND90 and 6 month-old animals.* The curves represent the fit with the sum of a linear and step function and also a step model when relevant. (A) Log of mammary gland whole mount weight in PND90SD. (B, C, D) Density analysis in the anterior area (area 3) in PND90CD, 6MCD and 6MSD respectively. In all cases, the complete model is not significant for all our statistical criteria; however, the step model is significant in B and D. Number of animals per group n=8-10.

Interestingly, the same quantity can be described by our model in PND90CD, 6MCD and 6MSD (Figure 9B, C, D). This quantity is the branching density of the posterior region of the mammary gland, closest to the 5^th^ mammary gland (area 3). In PN90CD, the complete model is not a good fit (p=0.7 for b) but it is still better than a constant model (p=0.033, q=0.038). The step model alone is a good fit (p=0.011) and is better than a constant model (p=0.0094).

In 6MCD, the model is a good fit (p=0.058, 0.024 for *b* and *c* respectively), is almost significantly better than a constant model, and is better than a linear or step model (p=0.064, 0.020, 0.049 respectively, q=0.13).

In 6MSD, the situation is similar (0.11, 0.017 for *b* and *c*), is better than a constant and a linear model, and almost significantly better than a step model (p=0.026, 0.014, 0.098 respectively, q=0.046). A step model is a good fit, with (p=0.036) and better than a constant model (p=0.032).

#### Comparison between negative control, BPA inflection point and positive control (0.5EE2): In PND21C

Because an inflection point was detected between 25BPA and 250BPA for several features, such as ‘sd width 3D’ or the ‘fractal dimension in 3D’ (Figure 8), we systematically investigated differences between 250BPA and control using the t-test for all features observed. We assessed the false discovery rate due to multiple testing with a threshold of 0.25. In the cases where 250BPA is significantly different from control, we investigated whether the effect of the 0.5EE2 dose was comparable to the effect of 250BPA (hypothesis (i)) or was qualitatively different (hypothesis (ii)). Since we investigated individual quantities, we decided against (i) and for (ii) when the effect of 0.5EE2 is lower than the one of 250BPA, which we tested by a t-test. (ii) can correspond to two different situations, (iia) there is no effect of 0.5EE2 by comparison with control or, alternatively, (iib) the effect of 0.5EE2 is opposite to the effect of 250BPA. We used Bonferroni corrections to control multiple comparisons among treatments. Results from these comparisons are summarized and illustrated in Figure 10.

**Figure 10:**
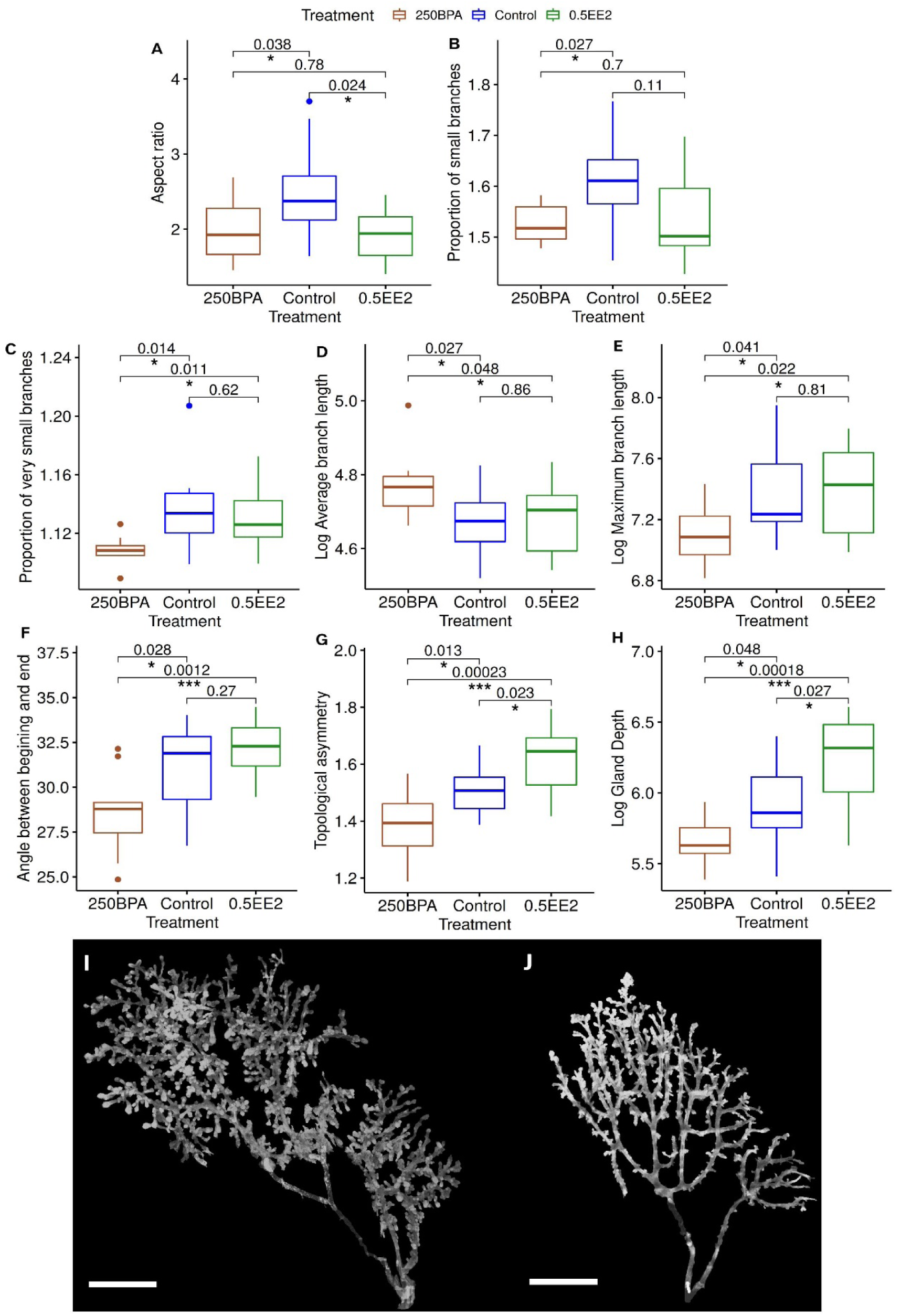
*Comparisons between Control and 250BPA, control and 0.5EE2, and 250BPA and 0.5EE2 in PND21C. A-H Box plot of several quantities for which the mean is significantly different between Control and 250BPA.* p-values correspond to t-test for Control versus 250BPA and t-test to assess whether the effect of EE2 was similar to the one of BPA, corrected for multiple comparisons. In all cases, the distributions do not differ significantly from a Gaussian by the Shapiro test. Number of animals per group n=8-10. Graphs A-B show quantities where the effects of 250BPA and 0.5EE2 are similar, matching. hypothesis (i). Graphs C-E show quantities where 250BPA is different from Control but 0.5EE2 is not, which fits hypothesis (iia). Graphs F-H show features where the effects of 250BPA and 0.5EE2 are opposite, matching hypothesis (iib). Panels I-J illustrate the morphological differences between Control (I) and 250BPA (J) in PND21C animals. These samples are chosen because they exhibit the differences outlined in A-H. Scale bars = 2mm.

##### Aspect Ratio (AR)

Glands of rats exposed to 250BPA were rounder (had a smaller aspect ratio) (p=0.038, q=0.21) compared to controls. 0.5EE2 was similar to 250BPA (p=0.78) and rounder than control (p=0.024). The response to BPA and EE2 were similar, consistent with hypothesis (i) (see Figure 8F, 10A).

##### Branching /budding

The proportion of small branches (buds and small ducts <15 pixels=75µm) was smaller in 250BPA than in control (p=0.027, q=0.21). For 0.5EE2, this quantity was between 250BPA and control, and closer to 250BPA (Figure 10B). This feature suggests a similar effect of BPA and EE2; compatible with hypothesis (i). The proportion of very small branches (<4pixels=20µm) can be interpreted as epithelial budding. There was fewer very small branches in 250BPA than in the control (p=0.014, q=0.21). Against hypothesis (i), there was more very small branches in 0.5EE2 than in 250BPA group (p=0.011). 0.5EE2 was comparable to the control (Figure 10C). This feature matches hypothesis (iia): BPA impacts a feature that EE2 does not.

##### Branch length measurements

Branches were defined computationally as a path from a bifurcation to the next bifurcation. The branches were longer on average in 250BPA than in control (p=0.027, q=0.21), but their maximum length was smaller (p=0.041, q=021) (Figure 10D and 10E). Similar results were obtained when the terminal branches were removed (“pruning”) (p=0.022 and 0.041, q=0.21 and 0.21 respectively). Against hypothesis (i), 0.5EE2 was different from 250BPA (p=0.048, 0.023, 0.039 and 0.032 respectively) and was similar to control for all these endpoints (p>0.8). These results matches hypothesis (iia).

##### Angle of branches

when removing small structures (<75µm), the remaining ducts tended to be straighter (turn less) in 250BPA than control (p=0.028, q=0.21). Against hypothesis (i), 250BPA ducts were also straighter than in 0.5EE2 (p=0.0012). 0.5EE2 seems to have an opposite effect than 250BPA but it is not significant; therefore, hypothesis (ii) holds but we cannot decide between (iia) and (iib) (Figures 8D, 10F).

##### Topological asymmetry

The average of the number of branching points from a branch to a terminal end (average depth of subtrees) is high when the epithelial tree is more asymmetric and low when, on the opposite, it is more balanced or compact topologically. Epithelial trees were more symmetric (p=0.013, q=0.21) in 250BPA than in control. Against hypothesis (i), trees were also more symmetric in 250BPA than in 0.5EE2, (p=0.00023). Against hypothesis (iia), Trees were also more symmetric in control than in 0.5EE2 (p=0.023), Figure 10G. Here, BPA and EE2 had opposite effects, matching hypothesis (iib).

##### Depth of the gland

similarly, the overall size of the epithelium along the z axis was lower in 250BPA than in control (p=0.048, q=0.22). The depth was also lower in 250BPA than in 0.5EE2 (p=0.00018) which invalidates hypothesis (i). Against hypothesis (iia), it was higher in 0.5EE2 than in control (p=0.027). This feature also matches hypothesis (iib) (Figure 10H).

We also performed multiple comparisons between all treatments and control using Dunnett’s test. Despite the important loss of statistical power when performing all comparisons, we found that the average branch length was significantly longer for 2.5BPA than for control, both without (p=0.031) and with (p=0.019) pruning.

As illustrated in Figure 10, some of the features analyzed are consistent with hypothesis (i), similar response to BPA and EE2. Others are consistent with hypothesis (ii) where BPA impact features that EE2 does not impact (iia) and in some cases, BPA has opposite effects than EE2 (iib).

#### In PND 90 and 6-month (data not shown)

We used the same methodology as above. In PND90CD, the gland density was lower on average in 250BPA-exposed glands than in vehicle (p=0.020, q=0.11). 0.5EE2 was between these two conditions so that we cannot decide between (i) and (ii). Interestingly, when the three distinct gland regions (rostral, middle and caudal) used to determine gland density were examined independently, there were treatment-dependent differences. Exposure to 0.5EE2 led to a significantly decreased gland density in the middle of the gland (area 2) compared to vehicle (p=0.011). Density in the rostral area (area 1) was lower in the 250BPA group than for females dosed with either vehicle or 0.5EE2 (p=0.031, q=0.099 and p=0.016 respectively) which is against hypothesis (i). 0.5EE2 is similar to control which is consistent with hypothesis (iia). Lobuloalveolar budding was higher in 250BPA and 0.5EE2 than in vehicle mammary glands (p=0.0049, q=0.08 and p=0.049 respectively, permutation test), which is consistent with hypothesis (i).

Performing comparisons between control and other continuously-dosed treatment groups showed that gland density (area 2) in 25BPA and 25000BPA was lower than in vehicle controls (p=0.047 and 0.0098 respectively, corrected by Dunnett’s test).

In PND90SD, lateral budding was higher in 250BPA than in vehicle, albeit not significantly (p=0.095 by permutation test). In agreement with hypothesis (i), the response was similar in 0.5EE2 treated females (p=0.0063 by permutation test).

In 6MCD, the fat pad area was larger in 250BPA than in vehicle controls (p=0.030, q=0.17) when 0.5EE2 was not distinct from either conditions. The percent coverage was lower in 250BPA than in control and 0.5EE2 (p= 0.022 q=0.17 and p=0.012 respectively) which is against hypothesis (i). The percent coverage was higher in EE2 than control albeit not significantly (p=0.083), which does not decide between (iia) and (iib).

In 6MSD, the standard deviation of gland density was higher in 250BPA than in control (p=0.012, q=0.067). Comparisons with 0.5EE2 were not significant albeit the situation was closer to hypothesis (iia). Lateral branching was lower in 250BPA than in control (p=0.011, permutation test, q=0.067), 0.5EE2 almost significantly lower than 250BPA and was similar to control, matching hypothesis (iia) (p=0.087 permutation test). Lateral budding and alveolar budding were almost significantly lower in 250BPA than control (p=0.082, 0.054; q=0.12, 0.12 respectively, permutation test) and were significantly lower in 250BPA than in 0.5EE2 (p=0.0046, 0.019 respectively, permutation test). 0.5EE2 is similar to control, therefore this result is consistent with hypothesis (iia).

### Other results

#### Comparison between the semi-automated measurements and the semi-quantitative scoring of PND21C glands

We compared the automated quantitative measurements of the glands with the developmental scores reported above for the chronic study and found correlations between this score and numerous morphological features. Quantities representative of the highest correlations with the score are the 2D fractal dimension of the gland (CC: 0.88, p=7.7e-27) and the number of branches (CC: 0.86, p=4.5 e-24).

The developmental score was also compared with the dimensions resulting from PCA. Table 4 shows that the scoring captured aspects of the two first dimensions of PCA (∼ size and thickness of glands, respectively) and was not correlated to Dim 3 (∼length of ducts) or to any additional dimensions. This relationship between the developmental score and the dimensions of PCA is meaningful, because it corresponds to the directionality of developmental characteristics observed between control and 0.5EE2 treated glands (Figure 11). In this sense, the developmental scoring criterion alone was optimized to detect effects resulting from exposure to EE2 but was not sufficient to detect significant non-linear responses in ductal length and several other morphological features that were shown by other analyses. Nevertheless, it is important to note that semi-quantitative scoring did show a non-significant non-monotonic response in morphological development between 25 and 250BPPA-exposed glands (Figure 4, S1 and S2).

**Figure 11:**
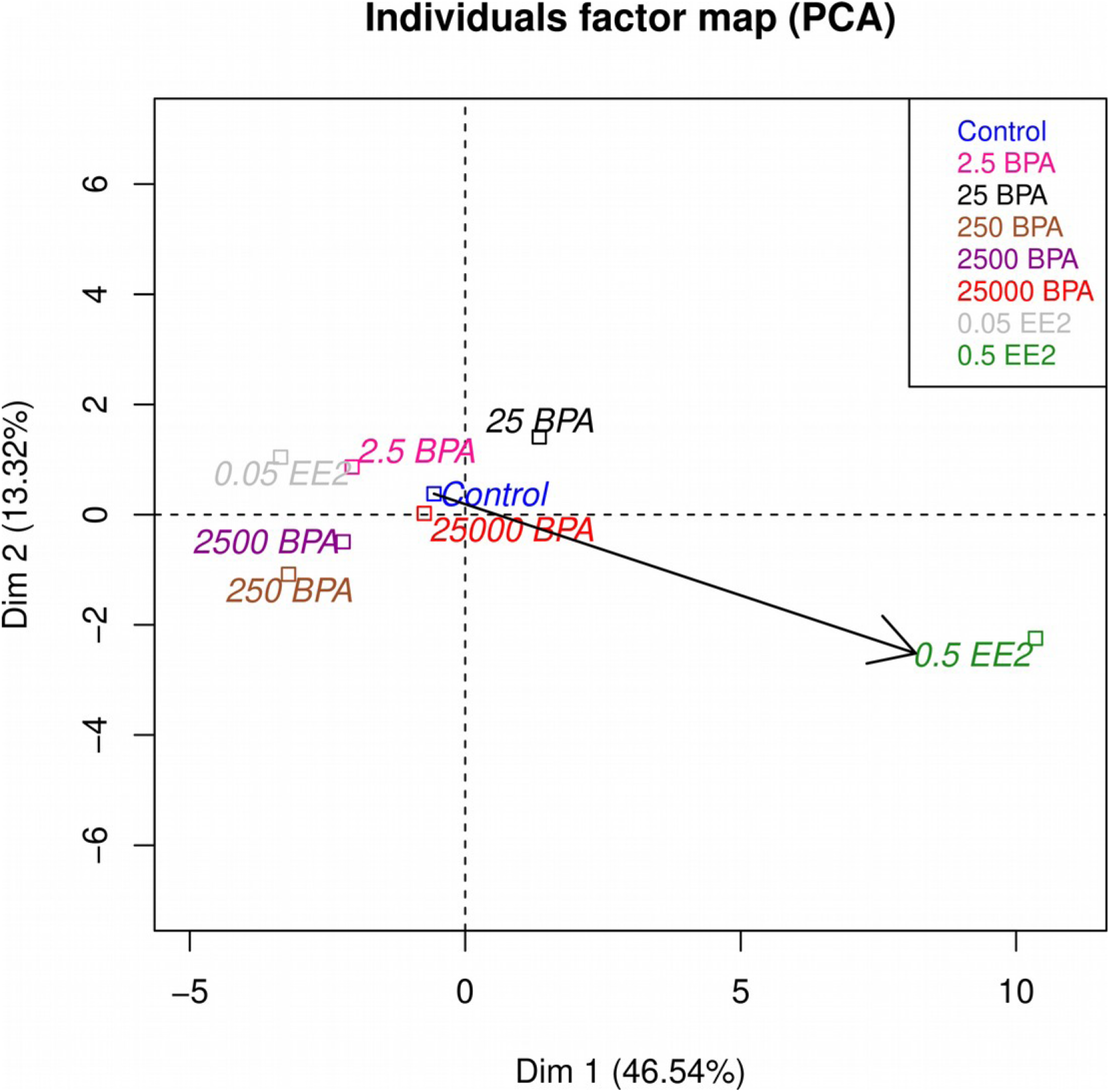
*Comparison between semi-quantitative scoring and the principal components from morphological analysis in PND21C animals*. The arrow represents the semi-quantitative scoring analyzed by PCA as a supplementary quantitative variable and corresponds clearly to the direction from control to 0.5EE2. The arrow’s direction does not capture the contrast between 25BPA and 250BPA which is almost orthogonal. Number of animals per group n=8-10.

**Table 4.** Correlation between semi-quantitative scoring and the dimensions of PCA in PND21C.

| Correlation descriptors | Dim 1 (~size) | Dim 2 (~local thickness) | Dim 3 (~ductal length) |
| --- | --- | --- | --- |
| Correlation coefficient | 0.88 | -0.29 | 0.023 |
| P value | 2.2e-16 | 0.011 | 0.84 |

Note: Only dimensions 1 and 2 are correlated with the scores which means that the summary of the data obtained by PCA is close to the hierarchy chosen in the design of mammary gland scores.

The standard deviation between the semi-quantitative assessments of the two observers scoring the glands provided further insight on its relationship with the response to BPA. We interpret this standard deviation as the result of an ambiguity in evaluating the development of some glands when this development is altered. This standard deviation is negatively-correlated with the proportion of small branches (-0.28, p=0.014) and the average thickness of the gland (-0.25, p=0.035). These features were relevant in our analysis of the non-monotonic response of BPA, where the non-monotonicity corresponded to a drop, consistent with a negative correlation. This suggests that the discrepancies between the assessments of the two observers were related to the non-monotonic response to BPA and the relatively two-dimensional evaluation of the gland on a typical microscope. BPA and EE2 resulted in different responses; while EE2 accelerated gland development, BPA led to abnormal development when assessed at PND21.

#### Histopathology in PND 90 and 6-month

Eight (8) lesions were identified in whole mounts and histological sections from eight PND90 mammary glands across both continuous and stop-dose treatment groups. No lesions manifested in vehicle-treated animals and all lesions were diagnosed as benign or malignant, ranging from lobular hyperplasia, fibroadenoma, periductular fibrosis or ductal epithelial necrosis with lymphocytic infiltration to ductal carcinoma *in situ* (DCIS) (Table S5A). Thirty-three (33) total lesions were identified in whole mounts and excised from twenty-four (24) 6-month mammary glands across both continuous and stop-dose treatment groups. Three (3) malignant tumors (adenocarcinomas) were classified from continuous and stop-dose 0.5EE2 treated females, and the remaining lesions/benign tumors were found in vehicle, 2.5BPA, 25BPA, and 25000BPA treated females. The benign lesions included lobular or ductular alveolar dilatations (with and without secretions), periductular fibrosis (with and without lymphocytic infiltration), fibroadenomas, and adenomas (Table S5B).

## Discussion

In a rare distribution of tissues from a very large guideline compliant study to academic grantees, we had an opportunity to evaluate mammary gland specimens from female rats from two studies on the effects of exposure to BPA and EE2 from fetal life to weaning (PND21) and beyond (PND90 and 6 months of age).

The mammary gland is considered a sensitive target for endocrine disruption; measurable effects manifest at low levels of exposure to endocrine disruptors, and these effects appear significantly earlier than the manifestation of mammary gland cancer. Thus, there is considerable interest in including its analysis in the animal tests used for regulatory purposes (Makris 2011;Rudel et al. 2011). However, the animal of choice for these regulatory studies was the NCTR-derived Sprague Dawley rat. In contrast to mouse models (Soto et al. 2013;Paulose et al. 2015), there is a paucity of reports on the effect of fetal BPA exposure on rat mammary gland morphogenesis. This is in part due to the florid structure of the ductal tree that grows more conspicuously into the third dimension and makes quantitative assessment beyond weaning challenging (Stanko et al. 2015). This feature of the rat mammary gland hinders the use of standard morphometric tools for the analysis of the rat mammary ductal system. Instead, conventional scoring methods are used. They are called semi-quantitative because they construct a score from qualitative and countable morphological features, such as terminal end buds (Table S1). These semiquantitative methods are reproducible, fast and reliable. Additionally, because the scoring method relies on the trained human eye which interprets structures in relation to function and pathology, the scores relate to biological outcomes.

A main objective of this study was to elucidate the shape of the dose-response curve and to test three alternative hypotheses: (i) that EE2 and BPA resulted in similar qualitative effects, (ii) that BPA and EE2 affected different features or had opposite effects, and (iii) that BPA had no effect on mammary gland development. Additionally, because the size and thickness of the PND21 mammary glands were compatible with confocal scanning and complete 3D reconstruction of the ductal tree, the PND21 female rats were extensively evaluated for the effects of BPA and EE2 using a quantitative new methodology specially developed for this study.

### Evaluation of early effects in the mammary gland

Consistent with our long history of evaluating both rat and mouse mammary whole mounts using semiquantitative and quantitative methods, we scored and measured glands from PND21 and PND90 in these studies, blinded to treatments. In addition, we postulated that an unsupervised, quantitative, and semiautomated method may discover effects that are difficult to ascertain using the scoring methods. We developed and describe here a method consisting of optical confocal sections to reconstruct the gland and use of appropriate algorithms for its analysis. The choice of PND21 was motivated mostly by the size of these glands and the fact that this pre-pubertal age precedes the florid and fast development of the ductal system due to ovarian estrogens; thus, estrogenic responses, if induced by BPA and EE2, should be detected. The hypothesis behind this choice is that the effect of BPA would be qualitatively similar to that of EE2; that is BPA will behave as a classical estrogen. We used the same set of mammary glands to compare this new quantitative method with the standard semi-quantitative method (Davis and Fenton 2013). Both the semi-quantitative and the quantitative methods were able to detect significant differences between the negative control (vehicle) and the positive control (0.5EE2) (Figures 4, 5, 7). However, the results obtained by both methods did not support the default hypothesis used in the experimental design, that BPA and EE2 would produce the same effect on the developing mammary gland.

### Evidence for a breaking point between 25 and 250 BPA in the dose response

An important motivation for the development of the quantitative assay was to obtain a precise evaluation of non-monotonicity. There are inherent differences in assumptions about the shape of the dose-response curve in endocrinology and toxicology. The default assumption in toxicology is monotonicity. In contrast, non-monotonic dose-response curves are a common occurrence in endocrinology. For instance, the proliferative response for estrogens and androgens follows inverted-U patterns (Stormshak et al. 1976;Amara and Dannies 1983;Soto et al. 1995;Maffini et al. 2002;Geck et al. 2000) and the effect of estradiol on the growth of the mammary ductal system is also non-monotonic (Vandenberg et al. 2006). Moreover, distinct end points show different estrogen dose-response curves in different organs of the same animal set; while the uterotrophic assay and various other uterine morphological end points are clearly monotonic, those pertaining to ductal mammary gland morphogenesis show mostly non-monotonic dose-response curves. As expected from examples in the literature, the BPA dose-response showed evidence for non-monotonicity on data from the quantitative method in PND21C (Jenkins et al. 2011;Cabaton et al. 2011). The dose-response curves observed for several features was not that of an inverted-U shape, instead it seemed to be characterized by a sudden drop or a breaking point located between 25BPA and 250BPA (Jenkins et al. 2011).

We used the 91 distinct measurements obtained with the semi-automated method for the analysis of PND21C glands to formulate a statistical test to assess whether 25-250BPA was the locus of a breaking point for most features. In this dataset, our exploratory analysis by the permutation test led us to reject the hypothesis that BPA has no effect in favor of a breaking point between 25 and 250BPA (Figure 6). To confirm this exploratory result, we used the smaller number of quantitative end points measured at PND90 and 6 months of age. The non-monotonic pattern of the dose-response to BPA was confirmed by a single, global statistical analysis using the same permutation test. The key of this global analysis is the hypothesis that the breaking point between 25 and 250BPA is present at all time points. Again, the test leads us to reject the hypothesis that BPA has no effect in favor of a breaking point between 25 and 250 BPA. We want to emphasize that performing this single test as a confirmatory analysis, when PND21C is used for the exploratory analysis, is a very rigorous method because it avoids making multiple comparisons for the many features and several datasets available. Moreover, the permutation test rigorously accommodates the individuality of each animal (i.e., the test takes into account that many features are correlated). Our overall strategy has been to build on the non-linear feature observed in PND21C, the breaking point, in order to provide evidence that it remains valid in the other datasets taken together. In the 90-day subchronic pilot studies (Figures S1 and S2), the increased development scores in the lowest BPA dose groups is another confirmation of this rationale, even though it failed to reach significance.

Once we have concluded that 25-250BPA is a breaking point, we can perform an exploratory analysis of which features are involved and the more specific shape of the dose response. Here, the analysis is performed with datasets already taken into account above, therefore it cannot provide a confirmation of the presence of the breaking point; however, it is useful to assess the overall response curve and the features that best represent it. The model chosen was the sum of a linear response and step function, because it is significantly better than either a linear model or a step function alone (Figure 8). In PND21C, for many variables, our model appears as a linear response at low doses; a drop in the response appears between 25 and 250BPA for most relevant features. At higher BPA doses, a linear response is observed again (Figure 8). The most striking feature of the dose response curve is the non-linearity of the response that takes place between the 25 and 250BPA dose. In other datasets, specific quantities were also non-monotonic such as gland weight in PND90SD and branching density in PND90CD, 6MCD and 6MSD. Moreover, pair-wise and comparisons between 250 BPA, 0.5EE2 and control revealed that some features are consistent with hypothesis (i) and others with hypothesis (ii) (Figure 10).

These results show the importance of establishing and using statistical methods appropriate for non-monotonic responses. Linear models are a powerful tool to provide evidence of a causal relationship because they quantitatively relate the changes of a putative cause with the one of the effects. Moreover, linear responses to small causes are a common mathematical property albeit not universal. Therefore, exhibiting a linear response is a powerful method to provide empirical evidence of a causal relationship in a given context. However, this method is blind to non-monotonic responses. The latter are common in endocrinology because the putative causes are involved in multilevel, complex regulations due to the evolutionary history of hormones and their functions. In this context, a more appropriate way to show the presence of causation is to show the prevalence of a specific non-monotonic pattern, here a breaking point between 25 and 250BPA.

### Semi-quantitative scoring and quantitative analyses

In PND21C, the scoring method captured the directionality of development as the arrow linking controls with 0.5EE2-treated (Figure 11) in the first two dimensions of PCA of data obtained by the quantitative method (correlated respectively with size and thickness). Indeed, the fact that the scoring method uses EE2 as the control implies that it would preferentially capture the effects of BPA when they mimic those of natural estrogens. Many of the features measured by the quantitative method do not relate directly to the features used for the scoring method, but still provide information about them. For example, the *fractal dimension* assesses the complexity of the ductal system and the *mean variation of ductal thickness* is associated with budding. Several of these unique features revealing significant BPA effects when evaluated in 3-dimensions are illustrated in Figure 8.

In PND21C, the data and analysis presented here demonstrate that the quantitative method, by using a multitude of automatic measurements, resulted in a greater sensitivity than the semi-quantitative method to discriminate effect differences due to BPA dose. The difference between the methods reside in the fact that the quantitative method does not depend on a positive control to score development, and thus is blind to whether the effect of BPA is similar or not to that of EE2 and that the quantitative method interrogates effects existing in the third dimension (i.e., thickness, fractal dimension in 3D, angles).

In PND90 and 6months, the scoring method revealed a significant effect of BPA in PND90P mammary glands exposed to BPA from gestation to sacrifice. This effect was observed only at the BPA2.5 dose and only when the animals were sacrificed at estrus, a result consistent with the non-monotonicity observed at all time points using the quantitative methods. These results stress the importance of assessing tissues at the same stage of the estrous cycle. In fact, the effect of EE2 0.5 on an increased developmental score was not evident when glands from animals in all stages of the estrous cycle were considered (Figure S2B). Moreover, these data also suggest that unlike the results in prepuberal PND21 animals, BPA may act in conjunction with endogenous estrogens in adult animals and thus produce a more estrogen-like pattern than that observed for BPA at PND21; that is, consistent with hypothesis (i).

The data presented here demonstrated that PND90 is an appropriate time point to assess the effect of low BPA doses and to reveal non-monotonicity. It also highlighted the importance of assessing the tissue at the same stage in the estrous cycle, as even the pronounced proliferative effect of the positive control on mammary epithelium was not evident when estrous stage was not taken into account in the data analyses. Our results also suggest that EE2 was not a good control for mammary gland end points as the effects of BPA and EE2 were distinct and there was no effect of the 0.05EE2, as expected; leaving us to suppose that the NCTR rat strain, may be particularly estrogen-insensitive. In summary, most of the results in these sets of mammary glands from cycling rats were consistent with hypothesis (ii), and inconsistent with hypothesis (iii).

### Cancer: this study and the core study

Like in the mouse model, some effects of BPA are not similar to those of estrogens, for example, inhibition of ductal growth at puberty (Markey et al. 2001;Munoz de Toro et al. 2005), enhanced ductal growth during fetal life (Vandenberg et al. 2007;Speroni et al. 2017), others are clearly estrogen-like, such as the increased score at PND90P reported here and the accelerated expression of lateral branching in the mouse (Munoz de Toro et al. 2005). There are also other effects seemingly unrelated to estrogenicity; changes in the stromal fraction of the gland and inflammatory cell responses which have been noted (Tucker et al. 2018;Wadia et al. 2013) in response to developmental BPA exposures.

Given that BPA is rapidly metabolized and does not bioaccumulate, the increased propensity of developing mammary cancer in animals exposed to BPA during organogenesis has been attributed to its direct effect on fetal mammary gland development and its indirect effects through the developing hypothalamic pituitary ovarian axis (HPOA) (Soto et al. 2013). In the current study, both PND90 and 6 months stop dose animals displayed a non-monotonic response to BPA, which confirms the long lasting effects of early BPA exposure (Table 2). The direct effect of BPA on fetal mammary gland development has been verified using fetal mammary gland explants in an *ex vivo* model (Speroni et al. 2017). Fetal exposure to BPA affects all the organs of the HPOA, altering ovarian steroidogenesis (Mahalingam et al. 2017;Peretz et al. 2011), hypothalamic controls of LH levels (Rubin et al. 2006;Acevedo et al. 2018), and the gonadotroph number in the fetal pituitary (Brannick et al. 2012). These alterations, in turn, suggest altered regulation of mammotropic hormones (Soto et al. 2013). Consistent with these findings, fetal exposure to BPA in mice not only affected the fetal period of mammary gland organogenesis, but also postnatal development, long after cessation of exposure. Alterations in ductal elongation at puberty and lateral branching and budding during adulthood were attributed to altered responses to mammotropic hormones (Wadia et al. 2007;Ayyanan et al. 2011). Recent studies confirmed that developmental exposures to other BPA related substances (BPS and BPAF) in mice also induce precocious development of the mammary epithelium, and increased epithelial lesions and mammary tumors in adulthood (Tucker et al. 2018). However, these results were obtained in the mouse, which is not considered as good a model for mammary cancer as the rat. In spite of this widely held opinion, developmental exposure to BPA in mice also increased the incidence of mammary cancer in animals treated with a chemical carcinogen during adulthood, or in MMTV-erbB2 mice exposed to BPA during adulthood (Jenkins et al. 2011). It is remarkable than in this model, the effect of BPA was non-monotonic. Several studies using different rat strains reported the development of hyperplasia, carcinoma in situ and palpable adenocarcinomas of the mammary gland after prenatal or neonatal exposure to BPA. Not surprisingly, the CLARITY core study, run concurrently to this study, revealed a significant increase of adenocarcinomas as well as the combination of adenomas or adenocarcinomas in the stop dose animals treated with 2.5BPA at 2 years of age. EE2 induced a significant increase of adenocarcinomas only at the high dose and were also detected in our animals (Table S5) by 6 months of age.

## Conclusions

Here we demonstrated that semi-quantitative and quantitative methods were suitable to detect estrogenic effects in the mammary glands of NCTR SD rats, and both methods found BPA-induced mammary effects to be different from those of EE2. Additionally, the semi-quantitative method, by relying on the trained human eye, is better able to interpret structures in relation to function and pathology. The automatic quantitative method, by using a multitude of measurements in 3D, identified statistically significant differences and revealed a non-monotonic BPA dose-response curve. The non-monotonic response was confirmed by a global analysis of quantitative assessment in older animal sets. This result shows that we can and should take advantage of non-monotonic properties to perform statistical analysis rigorously, and that these features are not limited to quadratic responses.

Consistent with this finding, the CLARITY core study which used animals of the same cohort found that EE2 and BPA are not similar; EE2 increased the incidence of neoplastic lesions only at the highest dose, while BPA only increased their incidence at the lowest dose. The BPA effect was non-monotonic and differed between the stop dose and the continuous exposure regime. Thus, dose and duration of exposure contribute to the developmental and neoplastic outcomes. These data are consistent with the multiple non-GLP studies previously conducted demonstrating low dose BPA exposures induce more adverse responses than high doses and that some low-dose BPA responses are different from those of estrogens and of high dose BPA.

## Acknowledgments

We would like to acknowledge the technical help provided by Dr. Luisa Camacho and Dr. Barry Delclos regarding the generation of animals and the dissection of the mammary glands examined in this study We are also grateful to Dr. Barbara Davis for the histological assessment of the lesions and to our colleagues at Tufts University, Dr. Carlos Sonnenschein and Dr. Beverly Rubin for their critical reading of the manuscript. We thank Dr. Marie Tremblay-Franco (section of Statistics and Bioinformatics, Plateform MetaToul-AXIOM, INRA Toulouse) and Keith Shockley (NIEHS), for their critical reading and useful suggestions regarding statistical analysis. This work was supported by Grant Number U01ES020888, from the National Institute of Environmental Health Sciences (NIEHS; A.M.S.) and NIEHS Funding 1Z01ES102785 (S.E.F. & M.B.). The content is solely the responsibility of the authors and does not necessarily represent the official views of the NIEHS or the National Institutes of Health.

## References

Acevedo N, Davis B, Schaeberle CM, Sonnenschein C, Soto AM. 2013. Perinatally administered Bisphenol A as a potential mammary gland carcinogen in rats. Environ Health Perspect 121:1040–1046.

Acevedo N, Rubin BS, Schaeberle CM, Soto AM. 2018. Perinatal BPA exposure and reproductive axis function in CD-1 mice. Reprod Toxicol 79:39–46.

Amara JF, Dannies PS. 1983. 17 β-Estradiol has a biphasic effect on GH cell growth. Endocrinology 112:1141–1143.

Ayyanan A, Laribi O, Schuepbach-Mallepell S, Schrick C, Gutierrez M, Tanos T et al. 2011. Perinatal exposure to bisphenol A increases adult mammary gland progesterone response and cell number. Mol Endocrinol 25:1915–1923.

Bolte S, Cordelières FP. 2006. A guided tour into subcellular colocalization analysis in light microscopy. J Microsc 224:213–232.

Brannick KE, Craig ZR, Himes AD, Peretz JR, Wang W, Flaws JA et al. 2012. Prenatal exposure to low doses of bisphenol A increases pituitary proliferation andgonadotroph number in female mice offspring at birth. Biol Reprod 87:82.

Cabaton NJ, Canlet C, Wadia PR, Tremblay-Franco M, Gautier R, Molina J et al. 2013. Effects of low doses of bisphenol A on the metabolome of perinatally exposed CD-1 mice. Environ Health Perspect 121:586–593.

Cabaton NJ, Wadia PR, Rubin BS, Zalko D, Schaeberle CM, Askenase MH et al. 2011. Perinatal exposure to environmentally relevant levels of Bisphenol-A decreases fertility and fecundity in CD-1 mice. Environ Health Perspect 119:547–552.

Calafat AM, Kuklenyik Z, Reidy JA, Caudill SP, Ekong J, Needham JL. 2005. Urinary concentrations of Bisphenol A and 4-Nonylphenol in a human reference population. Environ Health Perspect 113:391–395.

Churchwell MI, Camacho L, Vanlandingham MM, Twaddle NC, Sepehr E, Delclos KB et al. 2014. Comparison of life-stage-dependent internal dosimetry for bisphenol A, ethinyl estradiol, a reference estrogen, and endogenous estradiol to test an estrogenic mode of action in Sprague Dawley rats.Toxicol Sci 139:4–20.

Cohn BA, La Merrill M, Krigbaum NY, Yeh G, Park JS, Zimmermann L et al. 2015. DDT exposure in utero and breast cancer. J Clin Endocrinol Metab 100:2865–2872.

Dalmasso C, Broet P, Moreau T. 2005. A simple procedure for estimating the false discovery rate. Bioinformatics 21:660–668.

Davis B, Fenton S. 2013. The mammary gland, In: Handbook of Toxicologic Pathology (Rousseaux CG, Wallig MA, Haschek WM, eds), New York:Elsevier.

Delclos KB, Camacho L, Lewis SM, Vanlandingham MM, Latendresse JR, Olson GR et al. 2014. Toxicity evaluation of bisphenol A administered by gavage to Sprague Dawley rats from gestation day 6 through postnatal day 90. Toxicol Sci 139:174–197.

Diamanti-Kandarakis E, Bourguignon JP, Giudice LC, Hauser R, Prins GS, Soto AM et al. 2009. Endocrine-disrupting chemicals: an Endocrine Society scientific statement. Endocr Rev 30:293–342.

Doerge DR, Vanlandingham M, Twaddle NC, Delclos KB. 2010. Lactational transfer of bisphenol A in Sprague-Dawley rats. Toxicol Lett 199:372–376.

Doube M, Kosowski MM, Arganda-Carreras I, Cordelières FP, Dougherty RP, Jackson JS et al. 2010. BoneJ: Free and extensible bone image analysis in ImageJ. Bone 47:1076–1079.

Durando M, Kass L, Piva J, Sonnenschein C, Soto AM, Luque EH et al. 2007. Prenatal bisphenol A exposure induces preneoplastic lesions in the mammary gland in Wistar rats. Environ Health Perspect 115:80–86.

Geck P, Maffini MV, Szelei J, Sonnenschein C, Soto AM. 2000. Androgen-induced proliferative quiescence in prostate cancer: the role of AS3 as its mediator. Proc Nat Acad Sci USA 97:10185–10190.

Gerona RR, Woodruff TJ, Dickenson CA, Pan J, Schwartz JM, Sen S et al. 2013. Bisphenol-A (BPA), BPA glucuronide, and BPA sulfate in midgestation umbilical cord serum in a northern and central California population. Environ Sci Technol 47:12477–12485.

Goh WWB, Wong L. 2018. Dealing with cofounders in omics analysis. Trends Biotechnol 36:488–498.

Hehn RS. 2016. NHANES data support link between handling of thermal paper receipts and increased urinary bisphenol A excretion. Environ Sci Technol 50:397–404.

Heindel JJ, Newbold RR, Bucher JR, Camacho L, Delclos KB, Lewis SM et al. 2015. NIEHS/FDA CLARITY-BPA research program update. Reprod Toxicol 58:33–44.

Hoover RN, Hyer M, Pfeiffer RM, Adam E, Bond B, Cheville AL et al. 2011. Adverse health outcomes in women exposed in utero to diethylstilbestrol. N Engl J Med 365:1304–1314.

Interagency Breast Cancer and the Environment Research Coordinating Committee.Department of Health and Human Services (IBCERCC) Breast Cancer and the Environment: Prioritizing Prevention 2013 http://www.niehs.nih.gov/about/assets/docs/ibcercc_full_508.pdf

Jenkins S, Wang J, Eltoum I, Desmond R, Lamartiniere CA. 2011. Chronic oral exposure to Bisphenol A results in a non-monotonic dose response in mammary carcinogenesis and metastasis in MMTV-erbB2 mice. Environ Health Perspect 119:1604–1609.

Kostopoulos J, Chen WY, Gates MA, Tworoger SS, Hankinson SE, Rosner BA. 2010. Risk factors for ductal and lobular breast cancer: results from the nurses’ health study. Breast Cancer Res 12:R106.

Lamartiniere CA, Jenkins S, Betancourt AM, Wang J, Russo J. 2011. Exposure to the endocrine disruptor Bisphenol A alters susceptibility for mammary cancer. Horm Mol Biol Clin Investig 5:45–52.

Lê S, Josse J, Husson F. 2008. FactoMineR: An R package for multivariate analysis. Journal of Statistical Software 25.

Lewis SM, Lee FW, Ali AA, Allaben WT, Weis CC, Leakey JE. 2010. Modifying a displacement pump for oral gavage dosing of solution and suspension preparations to adult and neonatal mice. Lab Anim (N Y) 39:149–154.

Longo G, Montévil M. 2014. Perspectives on Organisms: Biological Time, Symmetries and Singularities, Berlin:Springer.

Luboz V, Wu X, Krissian K, Westin C-F, Kikinis R, Cotin S et al. 2005. A segmentation and reconstruction technique for 3D vascular structures, In: Medical Image Computing and Computer-Assisted Intervention -- MICCAI 2005: 8th International Conference, Palm Springs, CA, USA, October 26-29, 2005, Proceedings, Part I Berlin,Heidelberg:Springer Berlin Heidelberg, 43–50.

Maffini MV, Geck P, Powell CE, Sonnenschein C, Soto AM. 2002. Mechanism of androgen action on cell proliferation AS3 protein as a mediator of proliferative arrest in the rat prostate. Endocrinology 143:2708–2714.

Mahalingam S, Ther L, Gao L, Wang W, Ziv-Gal A, Flaws JA. 2017. The effects of in utero bisphenol A exposure on ovarian follicle numbers and steroidogenesis in the F1 and F2 generations of mice. Reprod Toxicol 74:150–157.

Makris SL. 2011. Current assessment of the effects of environmental chemicals on the mammary gland in guideline rodent studies by the U.S. Environmental Protection Agency (U.S. EPA), Organisation for Economic Co-operation and Development (OECD), and National Toxicology Program (NTP). Environ Health Perspect 119:1047–1052.

Markey CM, Luque EH, Munoz de Toro MM, Sonnenschein C, Soto AM. 2001. *In utero* exposure to bisphenol A alters the development and tissue organization of the mouse mammary gland. Biol Reprod 65:1215–1223.

Montévil M. 2018. A primer on mathematical modeling in the study of organims and their parts, In: Systems Biology. Methods in Molecular Biology (Bizzarri M, ed), New York,Humana Press.

Munoz de Toro MM, Markey CM, Wadia PR, Luque EH, Rubin BS, Sonnenschein C et al. 2005. Perinatal exposure to Bisphenol A alters peripubertal mammary gland development in mice. Endocrinology 146:4138–4147.

Murray TJ, Maffini MV, Ucci AA, Sonnenschein C, Soto AM. 2007. Induction of mammary gland ductal hyperplasias and carcinoma in situ following fetal Bisphenol A exposure. Reproductive Toxicology 23:383–390.

Nichols TE, Holmes AP. 2001. Nonparametric permutation tests for functional neuroimaging: A primer with examples. Human Brain Mapping 15:1–25.

Palmer JR, Hatch EE, Rosenberg CL, Hartge P, Kaufman RH, Titus-Ernstoff L et al. 2002. Risk of breast cancer in women exposed to diethylstilbestrol *in utero*: preliminary results (United States). CCC 13:753–758.

Paulose T, Speroni L, Sonnenschein C, Soto AM. 2015. Estrogens in the wrong place at the wrong time: Fetal BPA exposure and mammary cancer. Reprod Toxicol 54:58–65.

Peretz J, Gupta RK, Singh J, Hernández-Ochoa I, Flaws JA. 2011. Bisphenol A impairs follicle growth, inhibits steroidogenesis, and downregulates rate-limiting enzymes in the estradiol biosynthesis pathway. Toxicol Sci 119:209–217.

Phipson B, Smyth GK. 2010. Permutation P-values should never be zero: calculating exact P-values when permutations are randomly drawn. Stat Appl Genet Mol Biol 9:1585.

Preibisch S, Saalfeld S, Tomancak P. 2009. Globally optimal stitching of tiled 3D microscopic image acquisitions. Bioinformatics 25:1463–1465.

R Development Core Team R: A Language and Environment for Statistical Computing 2008 http://www.R-project.org

Rubin BS, Lenkowski JR, Schaeberle CM, Vandenberg LN, Ronsheim PM, Soto AM. 2006. Evidence of altered brain sexual differentiation in mice exposed perinatally to low environmentally relevant levels of bisphenol A. Endocrinology 147:3681–3691.

Rudel RA, Fenton SE, Ackerman JM, Euling SY, Makris SL. 2011. Environmental exposures and mammary gland development: state of the science, public health implications, and research recommendations. Environ Health Perspect 119:1053–1061.

Schug TT, Heindel JJ, Camacho L, Delclos KB, Howard P, Johnson AF et al. 2013. A new approach to synergize academic and guideline-compliant research: the CLARITY-BPA research program. Reprod Toxicol 40:35–40.

Sheets KG, Jun B, Shou Y, Zhu M, Petasis NA, Gordon WC et al. 2013. Microglial ramification and redistribution concomitant with the attenuation of choroidal neovascularization by neuroprotectin D1. Molecular Vision 19:1747–1759.

Sonnenschein C, Soto AM. 2016. Carcinogenesis explained within the context of a theory of organisms. Prog Biophys Mol Biol 122:70–76.

Sonnenschein C, Wadia PR, Rubin BS, Soto AM. 2011. Cancer as development gone awry: the case for bisphenol-A as a carcinogen. Journal of Developmental Origins of Health and Disease 2:9–16.

Soto AM, Brisken C, Schaeberle CM, Sonnenschein C. 2013. Does cancer start in the womb? Altered mammary gland development and predisposition to breast cancer due to in utero exposure to endocrine disruptors. J Mammary Gland Biol Neoplasia 18:199–208.

Soto AM, Lin TM, Sakabe K, Olea N, Damassa DA, Sonnenschein C. 1995. Variants of the human prostate LNCaP cell line as a tool to study discrete components of the androgen-mediated proliferative response. Oncology Res 7:545–558.

Soto AM, Sonnenschein C. 2011. The tissue organization field theory of cancer: A testable replacement for the somatic mutation theory. BioEssays 33:332–340.

Speroni L, Voutilainen M, Mikkola ML, Klager SA, Schaeberle CM, Sonnenschein C et al. 2017. New insights into fetal mammary gland morphogenesis: differential effects of natural and environmental estrogens. Scientific Reports 7:40806.

Stanko JP, Easterling MR, Fenton SE. 2015. Application of Sholl analysis to quantify changes in growth and development in rat mammary gland whole mounts. Reprod Toxicol 54:129–135.

Stormshak F, Leake R, Wertz N, Gorski J. 1976. Stimulatory and inhibitory effects of estrogen on uterine DNA synthesis. Endocrinology 99:1501–1511.

Thayer KA, Taylor KW, Garantziotis S, Schurman S, Kissling GE, Hunt D et al. 2016. Bisphenol A, Bisphenol S, and 4-Hydroxyphenyl 4-Isoprooxyphenylsulfone (BPSIP) in urine and blood of cashiers. Environ Health Perspect 124:437–444.

Trichopoulos D. 1990. Is breast cancer initiated *in utero*? Epidemiology 1:95–96.

Tucker DK, Hayes Bouknight S, Brar SS, Kissling GE, Fenton SE. 2018. Evaluation of prenatal exposure to Bisphenol analogues on development and long-term health of the mammary gland in female mice. Environ Health Perspect 126:087003.

Vandenberg LN, Chauhoud I, Heindel JJ, Padmanabhan V, Paumgartten FJ, Schoenfelder G. 2010. Urinary, circulating and tissue biomonitoring studies indicate widespread exposure to Bisphenol A. Environ Health Perspect 118:1055–1070.

Vandenberg LN, Hunt PA, Myers JP, vom Saal FS. 2013. Human exposures to bisphenol-A: mismatches between data and assumptions. Rev Environ Health 28:37–58.

Vandenberg LN, Maffini MV, Wadia PR, Sonnenschein C, Rubin BS, Soto AM. 2007. Exposure to environmentally relevant doses of the xenoestrogen bisphenol-A alters development of the fetal mouse mammary gland. Endocrinology 148:116–127.

Vandenberg LN, Wadia PR, Schaeberle CM, Rubin BS, Sonnenschein C, Soto AM. 2006. The mammary gland response to estradiol: monotonic at the cellular level, non-monotonic at the tissue-level of organization? J Steroid Biochem Molec Biol 101:263–274.

Vandevoort CA, Gerona RR, vom Saal FS, Tarantal AF, Hunt PA, Hillenweck A et al. 2016. Maternal and fetal pharmacokinetics of oral radiolabeled and authentic Bisphenol A in the Rhesus Monkey. PLoS One 11:e0165410.

Villar-Pazos S, Martinez-Pinna J, Castellano-Muñoz M, Alonso-Magdalena P, Marroqui L, Quesada I et al. 2017. Molecular mechanisms involved in the non-monotonic effect of bisphenol-a on ca2+ entry in mouse pancreatic ß-cells. Sci Reports 7:11770.

Wadia PR, Cabaton NJ, Borrero MD, Rubin BS, Sonnenschein C, Shioda T et al. 2013. Low-dose BPA exposure alters the mesenchymal and epithelial transcriptomes of the mouse fetal mammary gland. PLoS One 8:e63902.

Wadia PR, Vandenberg LN, Schaeberle CM, Rubin BS, Sonnenschein C, Soto AM. 2007. Perinatal Bisphenol-A exposure increases estrogen sensitivity of the mammary gland in diverse mouse strains. Environ Health Perspect 115:592–598.

Zoeller RT, Brown TR, Doan L, Gore AC, Skakkebaek N, Soto AM et al. 2012. Endocrine-disrupting chemicals and public health protection: A statement of principles from The Endocrine Society. Endocrinology 153:4097–4110.

