## Supplemental materials for "A combined morphometric and statistical approach to assess non-monotonicity in the developing mammary gland of rats in the CLARITY-BPA study"

Current Affiliations:

[3]Institut de Recherche et d'Innovation, Centre Pompidou, Paris, France

[4]Elavo Mundi Solutions, LLC, Santa Monica, CA 90403, USA

[5]Department of Immunology, Tufts University School of Medicine, Boston MA 02111, USA

Corresponding author:
Ana M. Soto
Department of Immunology
Tufts University School of Medicine
136 Harrison Avenue
Boston, MA 02111


**Acknowledgments**: We would like to acknowledge the technical help provided by Dr. Luisa Camacho and Dr. Barry Delclos regarding the generation of animals and the dissection of the mammary glands examined in this study. We are also grateful to Dr. Barbara Davis for the histological assessment of the lesions. This work was supported by Grant Number U01ES020888 from the National Institute of Environmental Health Sciences (NIEHS; A.M.S.) and NIEHS Funding 1Z01ES102785 (S.E.F. & M.B.). The content is solely the responsibility of the authors and does not necessarily represent the official views of the NIEHS or the National Institutes of Health.

**Declaration of competing financial interests:** The authors declare they have no actual or potential competing financial interests

Table S1. Developmental scoring guideline used for morphological assessment of PND 21 mammary gland whole mounts following early life BPA or EE2 exposures.

| Score | Criterion Used in Semiquantitative Scoring |
|---|---|
| 1 | Poor development, small epithelial growth, minimal branching and budding, few/no TEBs, poor development of cranial aspect of gland 4 (asymmetric) |
| 2 | Gland almost reaches the lymph node (LN) (retarded growth), little branching or budding, few TEBs, poor development of cranial aspect of gland 4 |
| 3 | Gland touches LN, moderate branching and budding, external TEBs begin to appear around periphery, moderate development of cranial aspect of gland 4 |
| 4 | Gland touches LN, wide with equal antral and dorsal development (symmetric), internal and external TEBs, excellent branching and budding throughout gland, symmetric |
| 5 | Excessive lateral growth, gland has grown past LN, dense budding with few gaps, internal and external TEBs, external TEBs around entire periphery |
| 6 | Excessive lateral growth, growth beyond LN, 4th and 5th gland have grown together, dense budding with very few gaps, fewer TEBs because they are beginning to differentiate into lobules (looks like typical development on PND 35 or 50) |
| 7 | Excessive lateral growth, gland has reached ends of fat pads and are terminally differentiating into lobules, 4th and 5th glands have grown over each other, very dense, difficult to see ducts (looks like young adult gland) |

Notes: PND=Postnatal Day, TEBs=Terminal End Buds, LN=Lymph Node.

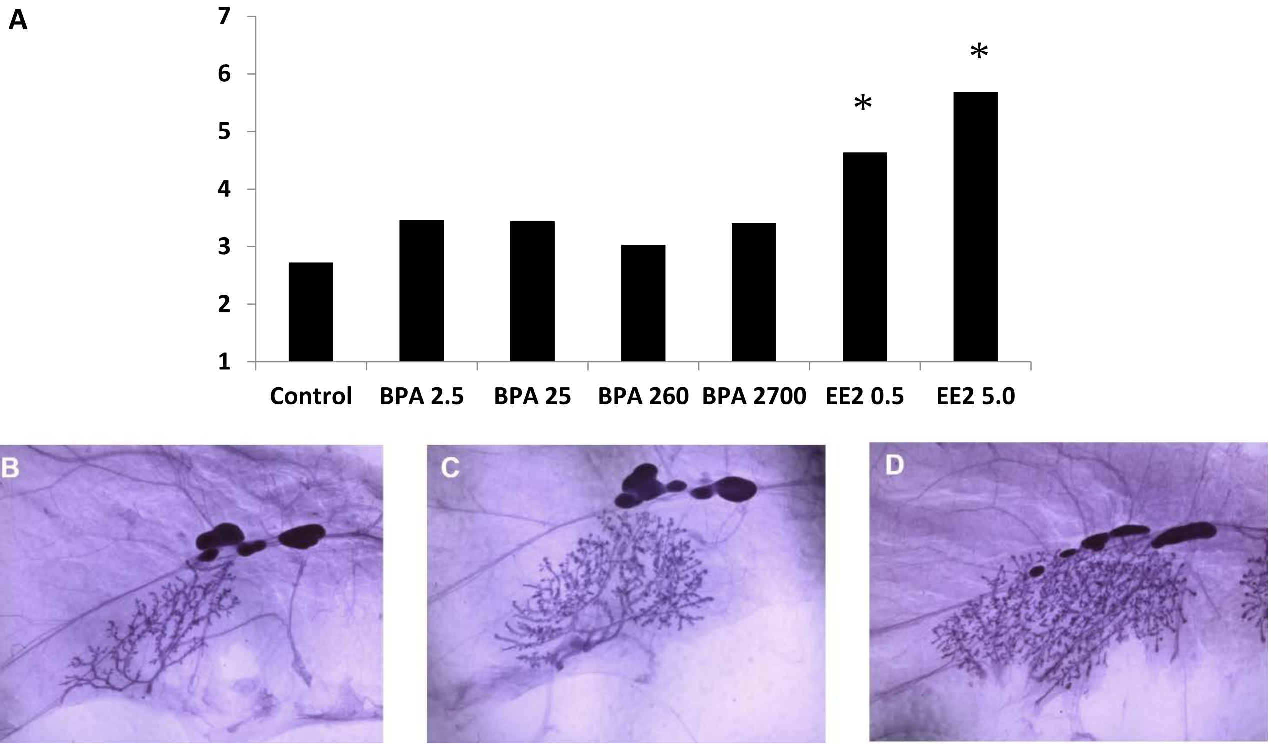


**Figure S1: Scoring evaluation of PND21P mammary glands**. [A] *Comparison of the mean score of all treatment groups.* Although there are no significant discernible effects in the subchronic PND21P mammary glands across BPA-exposed treatment groups, the reduced gland development in BPA260 compared to BPA25-exposed animals is consistent with the global analysis obtained from the chronic study animals (PND21C). Number of animals per group n=9-12. * indicates significantly accelerated gland development compared to vehicle controls (Kruskal Wallis; $p=0.004$ and $p<0.0001$). Images are representative of mammary gland development in [B] PND21P vehicle control group, [C] PND21P EE2 0.5 group, and [D] PND21P EE2 5.0 group.

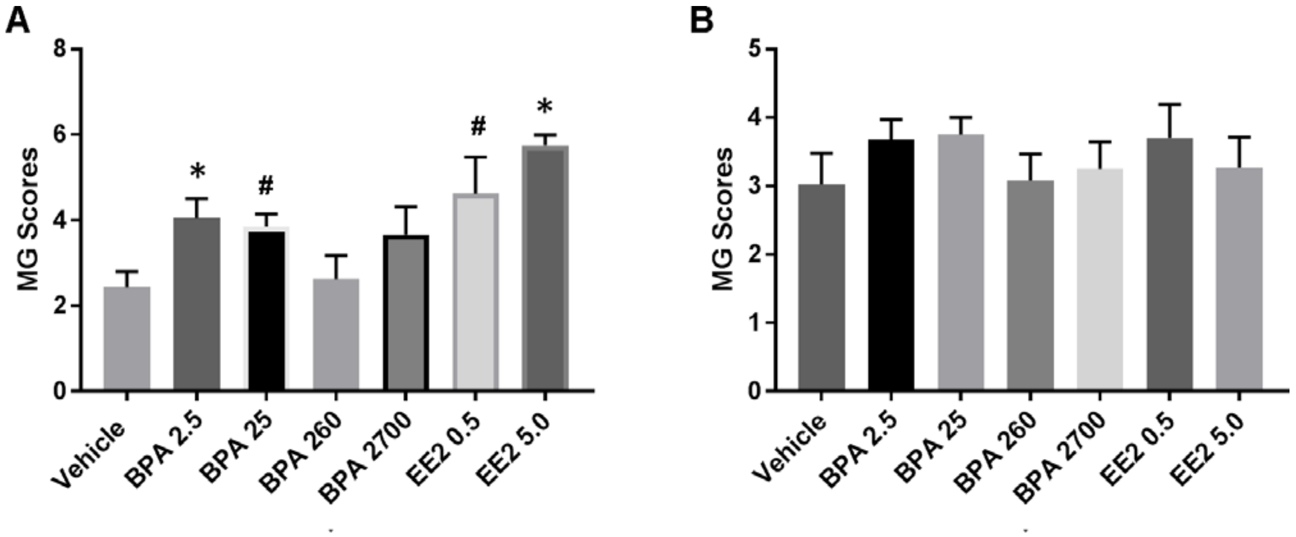


**Figure S2.** Semiquantitative scoring of postnatal day 90 pilot (PND90P) glands. A) PND90P animals from Fenton group in which the majority of animals were in estrus at necropsy (only females in estrus included; n=7, 10, 10, 4, 6, 4, 4; from left to right). A drop between BPA25 and BPA260, consistent with the global analysis of non-monotonicity exposed in the main text, was detected in animals in estrus. * Indicates significantly accelerated gland development compared to vehicle controls (Kruskal Wallis; BPA 2.5 p=0.05, EE5 p=0.01). # Indicates increased gland proliferation that did not reach significance (Kruskal Wallis; BPA 25 p=0.09, EE0.5 p=0.1). B) PND90P animals that were cycling from both Fenton and Soto groups, with all estrous cycle stages at necropsy included except anestrus (n=12, 18, 14, 10, 12, 12, 15, from left to right). All animals in A were included in B analysis. Note lack of effect of BPA and EE2 when cycle stage is disregarded.

**Table S2.** Features measured by the semi-automatic method applied to PND 21 mammary glands and complementary quantities used jointly in PCA and other analyses.

| Type of analysis performed | Feature Label | Explanation of Feature Label |
|---|---|---|
| Weights | Necropsy Weight (g) | Body weight at necropsy (grams) |
| | Mammary Gland Weight (mg) | Weight of mammary gland (milligrams) |
| Manual assessment | TEB | Number of terminal end buds |
| Analyses of the 2D projection of the mammary tree | Area (µm2) | Surface of 2D projection (square micrometers) |
| | Major (µm) | Size of the major axis of the gland (micrometers) |
| | Minor (µm) | Size of the minor axis of the gland (micrometers) |
| | Feret (µm) | Feret diameter (micrometers) |
| | AR | Aspect ratio of the gland |
| | Round | Roundness (inverse aspect ratio) |
| | Fractal Dimension | Self-explanatory (higher for denser glands, lower for sparse glands) |
| | Extension LV (µm) | Farthest distance from the lymph vessels; negative when it is not reached (micrometers) |
| | Vesselp | Proportion of the gland beyond a specific lymph vessel |
| | Nodep | Proportion beyond the lymph node |
| Global analyses in 3D | Width (µm) | Width of the gland along its main directions (micrometers) |
| | Height (µm) | Height of the gland along its main directions (micrometers) |
| | Depth (µm) | Depth of the gland along its main directions (micrometers) |
| | Vol (µm3) | Raw volume of epithelium (cubic micrometers) |
| | SA (µm2) | Surface of the epithelium (i.e., surface the boundary epithelium/stroma) (square micrometers) |
| | Solidity 3D (µm3) | Volume / convex volume (cubic micrometers) |
| | Encl Vol (µm3) | Volume with some corrections (cubic micrometers) |
| | I1 | Momentum of inertia along axis 1 |
| | I2 | Momentum of inertia along axis 2 |
| | I3 | Momentum of inertia along axis 3 |
| | Euler | Assessment of Euler characteristic, which provides information on the lack of convexity of the object |
| | Holes | Number of topological holes. |
| | Thickness (µm) | Average local thickness of the gland (estimates the diameter, but biased by the compression exerted on the gland) (micrometers) |
| | SD Thickness (µm) | Average local thickness of the gland (estimates the diameter, but biased by the compression exerted on the gland) (micrometers) |
| | Max Thickness (µm) | Average local thickness of the gland (estimates the diameter, but biased by the compression exerted on the gland) (micrometers) |
| | Dimension 3D | Fractal dimension in 3D - high if the gland fills space in 3 dimension (thick, no lacunarity, high budding, ...) |
| Direct skeleton analysis (raw) | X Branches | Number of branches |
| | X Junctions | Number of junctions |
| | X Junction Voxels | Number of junction voxels |
| | Average Branch Length (µm) | Branch length (micrometers) |
| | X Triple Points | Number of bifurcation |
| | X Quadruple Points | Number of triple branching |
| | Maximum Branch Length (µm) | Maximum branch length (micrometers) |

| | | |
|---|---|---|
| Direct skeleton analysis after pruning | X Branches1 | Number of branches (only for non-terminal branches) |
| | X Junctions1 | Number of junctions (only for non-terminal branches) |
| | X Junction Voxels1 | Number of junction voxels (only for non-terminal branches) |
| | X Slab Voxels1 | Number of voxels (only for non-terminal branches) |
| | Average Branch Length1 (µm) | Branch Length (micrometers) (only for non-terminal branches) |
| | X Triple Points1 | Number of bifurcation (only for non-terminal branches) |
| | X Quadruple Points1 | Number of triple branching (only for non-terminal branches) |
| | Maximum Branch Length1 (µm) | Maximum branch length (micrometers) (only for non-terminal branches) |
| Specialized analysis. When quantities are defined per branch the average over all branches is reported. All branches larger than 20µm are taken into account. | Size (µm) | Length of branch (micrometers) |
| | Number of Neighbors | Number of disregarded connections |
| | Depth from Root | Number of bifurcation from the nipple to the branch |
| | Depth Subtree (µm) | Average depth of the subtree of each branch (micrometers) |
| | Number of Children | Average number of sub branches |
| | Euclidean Distance (µm) | Distance between beginning and end of each branch (micrometers) |
| | Tortuosity | Ratio: length of branches /Euclidean distance |
| | Angle Between Beginning and End | Angle between beginning and end of a branch |
| | Angle with Parent Local | Angle between the end of the parent branch and the beginning the branch |
| | Angle with Parent Global | Angle between the direction of the parent branch and the branch |
| | Angle Wr Main Dir | Angle between the direction of the branch and the average direction of all branches |
| | Length to Nipple (µm) | Distance in the tree between a branch and the nipple (micrometers) |
| | Mean Width (µm) | Mean distance map of the branch without the z axis (i.e., 2D width of the branch) (micrometers) |
| | Max Width (µm) | Max distance map of the branch without the z axis (i.e., 2D width of the branch) (micrometers) |
| | SD Width (µm) | Standard deviation of the distance map of the branch without the z axis (i.e., 2D width of the branch) (micrometers) |
| | Mean Width2 (µm) | Mean local thickness of the branch (micrometers) |
| | Max Width2 (µm) | Max local thickness of the branch (micrometers) |
| | SD Width2 (µm) | Standard deviation of the local thickness of the branch (micrometers) |
| | Length Farthest Leaf (µm) | Distance in the tree between a branch and farthest leaf (micrometers) |
| | Topodepth | Total depth (number of bifurcation from nipple to the farthest branch) |
| | Nblarge | Putative bud clusters (structures with a wide end) |
| | Secondary Bud | Putative number of budding from ducts |
| | Nbbranchestree | Number of branches |
| | Type1 (%) | Percent secondary bifurcation |
| | Type2 (%) | Percent subbranches of secondary bifurcations |
| Specialized analysis. When quantities are defined per branch the average over all branches is reported. Only branches larger than 75µm are taken into account. | Size1 (µm) | Length of branches (micrometers) |
| | Number of Neighbours1 | Number of disregarded connections |
| | Depth from Root1 | Number of bifurcation from the nipple to the branch |
| | Depth Subtree1 (µm) | Average depth of the subtree of each branch (micrometers) |
| | Number of Children1 | Average number of sub branches |
| | Euclidean Distance1 (µm) | Distance between beginning and end of each branch (micrometers) |
| | Tortuosity1 | Ratio: length of branches /Euclidean distance |
| | Angle Between Beginning and End1 | Angle between beginning and end of a branch |

| | | |
|---|---|---|
| | Angle with Parent Local1 | Angle between the end of the parent branch and the beginning the branch |
| | Angle with Parent Global1 | Angle between the direction of the parent branch and the branch |
| | Angle Wr Main Dir1 | Angle between the direction of the branch and the average direction of all branches |
| | Length to Nipple1 (µm) | Distance in the tree between a branch and the nipple (micrometers) |
| | Mean Width1 (µm) | Mean distance map of the branch without the z axis (i.e., 2D width of the branch) (micrometers) |
| | Max Width1 (µm) | Max distance map of the branch without the z axis (i.e., 2D width of the branch) (micrometers) |
| | SD Width1 (µm) | Standard deviation of the distance map of the branch without the z axis (i.e., 2D width of the branch) (micrometers) |
| | Mean Width2.1 (µm) | Mean local thickness of the branch (micrometers) |
| | Max Width2.1 (µm) | Max local thickness of the branch (micrometers) |
| | SD Width2.1 (µm) | Standard deviation of the local thickness of the branch (micrometers) |
| | Length Farthest Leaf1 (µm) | Distance in the tree between a branch and farthest leaf (micrometers) |
| | Topodepth1 | Total depth (number of bifurcation from nipple to the farthest branch) |
| | Nblarge1 | Putative bud clusters (structures with a wide end) |
| | Secondary Bud1 | Putative number of budding from ducts |
| | Nbbranchestree1 | Number of branches |
| | Type1.1 (%) | Percent secondary bifurcation |
| | Type2.1 (%) | Percent subbranches of secondary bifurcations |

**Supplementary analyses by PCA**

PCA is a method for dimensional reduction, i.e. for summarizing data sets where many quantities are assessed simultaneously. The starting point of PCA is to build new quantities called dimensions (Dim, named Dim 1, Dim 2, etc.) as linear combinations of the original quantities, for example if *A*, *B*, *C* are quantities measured, Dim 1= $aA+bB+cC$ where *a*, *b* and *c* are determined by a computation. The new quantities are built to be independent of each other and to explain as much of the variance as possible. They are sorted by decreasing contribution to variance. The meaning of these dimensions with respect to the original quantities is proper to a given dataset because the coefficients *a*, *b*, *c*,… are different for different datasets. The strength of PCA is that it summarizes data in an automated fashion. Its limitation is that properties not included in the first or first few dimensions may still be relevant. As an example, the subtended area of a gland is correlated to many variables since it is a way to assess the "size" of the gland, but the abundance of epithelial structures per unit volume conveys a different biological meaning. The latter can be more relevant to the understanding of the effect of the treatment even though it can be independent of some "size" variations that dominate spontaneous variability. As a result, the first dimension of PCA may not necessarily be of biological interest when discussing the response to a treatment. Since the dimensions of PCA depend on the entire data set, the results of PCA will be different depending on whether we include the positive controls (0.5EE2 and 0.05EE2) in the analysis.

**PND21C PCA with EE2**

In addition to the correlation between PCA dimensions and mammary gland measurements examined in the main text we found that Dim 4 did not correlate with any of the treatments. Dim 5 was correlated with the aspect ratio (length/width, see Figure 6F for an illustration) of the gland (0.72, p=9.3e-14). For this variable, control was higher (p=0.043) than the other exposure conditions. Dim 7 was correlated with the maximum duct length (-0.64, p=3.3e-10). For Dim 7, 25BPA was higher than other conditions (p=0.018), while control and 0.5EE2 were lower (p=0.038 and 0.0060 resp.).

**PND21C PCA without EE2**

Dimensions 1 to 3 of PCA without EE2 are similar to those of the PCA analysis with EE2 (Figure S2); in the absence of EE2, dimension 3 separates 250BPA from other conditions (p=0.022).

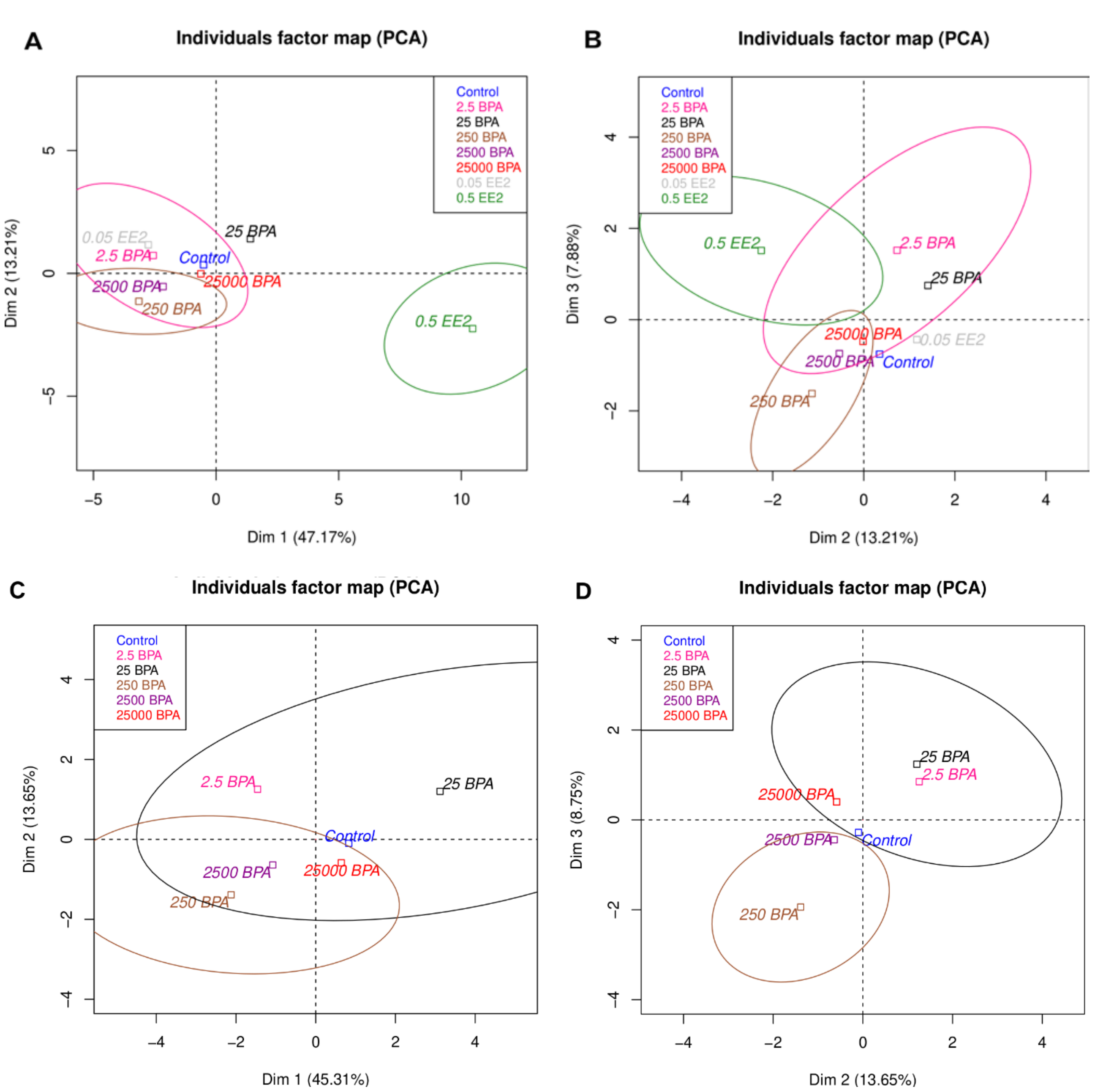


Figure S3. *Dimension 1 to 3 from PCA of PND21C animals with (top) and without (bottom) EE2 treatments.* We represent the average of each exposure group. Taking the EE2 conditions in the data sets changes the definition of the dimensions resulting from PCA. These changes are nevertheless limited (for example, for Dim 3, 2.5BPA is above 25BPA with EE2 while it is the opposite without EE2). In both cases, the dimensions obtained by PCA display a non-monotonic response. Control is between 25BPA and 250BPA. Number of animal per group n=8-10.

**PCA 90 days and 6 months with EE2**

For PND90CD, the Dim 1 of PCA is correlated with the average density of the gland (0.88, p=3.5e-26). For this feature, 2.5BPA are significantly higher than other conditions while 250BPA is lower (p=0.043, 0.019 respectively.).

For PND90SD, Dim 2 is related to the number of TEB (0.57, p=8.3e-8) and is higher in 0.5EE2 than other conditions (p=0.0032). Dim.4 is related to epithelial area (0.75, p=4.5e-15) and is lower in 0.5EE2 than in other conditions (p=0.0013.).

For 6MCD rats, Dim 1 is related to alveolar budding (0.78, p=3..8e-18) and for Dim 1, 0.5EE2 is lower than the other conditions (p=6e-8). Dim 4 is correlated with body weight (-0.66, p=1.5e-11), and 2500BPA is lower than other conditions (p=0.038).

For 6MSD animals, Dim 1 is related to lobular alveolar budding (0.65, p= 5.8e-11). For this quantity 0.5EE2 is higher and 250BPA is lower than the other conditions (p=0.00058 and 0.0018 resp.). Dim.4 is correlated with lateral budding (0.56, p=5e-8) and 250BPA is lower than other conditions (p=0.0081).

**Clustering on PCA**

Using only averages to assess the relationship between the data and the treatments is reductive in a biological situation where variability is high. Instead, clustering classifies the different individuals independently of the treatment and the outcome can show that a given kind of morphology (a cluster) is associated to a specific treatment. We used agglomerative hierarchical clustering on principal components with the Ward method implemented in factomineR. For the PND21C data, we used only the 15 (12 without EE2) first dimensions of PCA which means that we considered that other dimensions were not biologically meaningful. For other datasets, we used only the first 5 (4) dimensions since far fewer variables were measured. We partitioned the data in 10 (8) clusters in order to have more clusters than 8 (6) conditions. Then we compared statistically these clusters with animal treatments, which is a comparison between the factor “cluster” and the factor “treatment” that we perform using the v-test implemented in factomineR.

**PND21C with EE2**

We performed clustering on principal components and looked for correlations between clusters and animal treatments. We found clusters associated with 250BPA (p=0.0067) and two associated with 0.5EE2 (p= 5.5e-6, 0.014). This means that some glands in condition 250BPA are remarkable and different from the other conditions. To illustrate these clusters we display their paragons (or representative gland at center of that cluster) and most typical elements (representative gland at the farthest points in the cluster) in Figure S3.

**PND21C without EE2**

Clustering analysis defined a cluster correlated to 250BPA (p=0.0034) and another to 25BPA (p=0.028). This confirms that exposure 250BPA is distinct from the 25BPA one. The difference is due to the absence of EE2 data in this PCA and clustering which affects the clustering results.

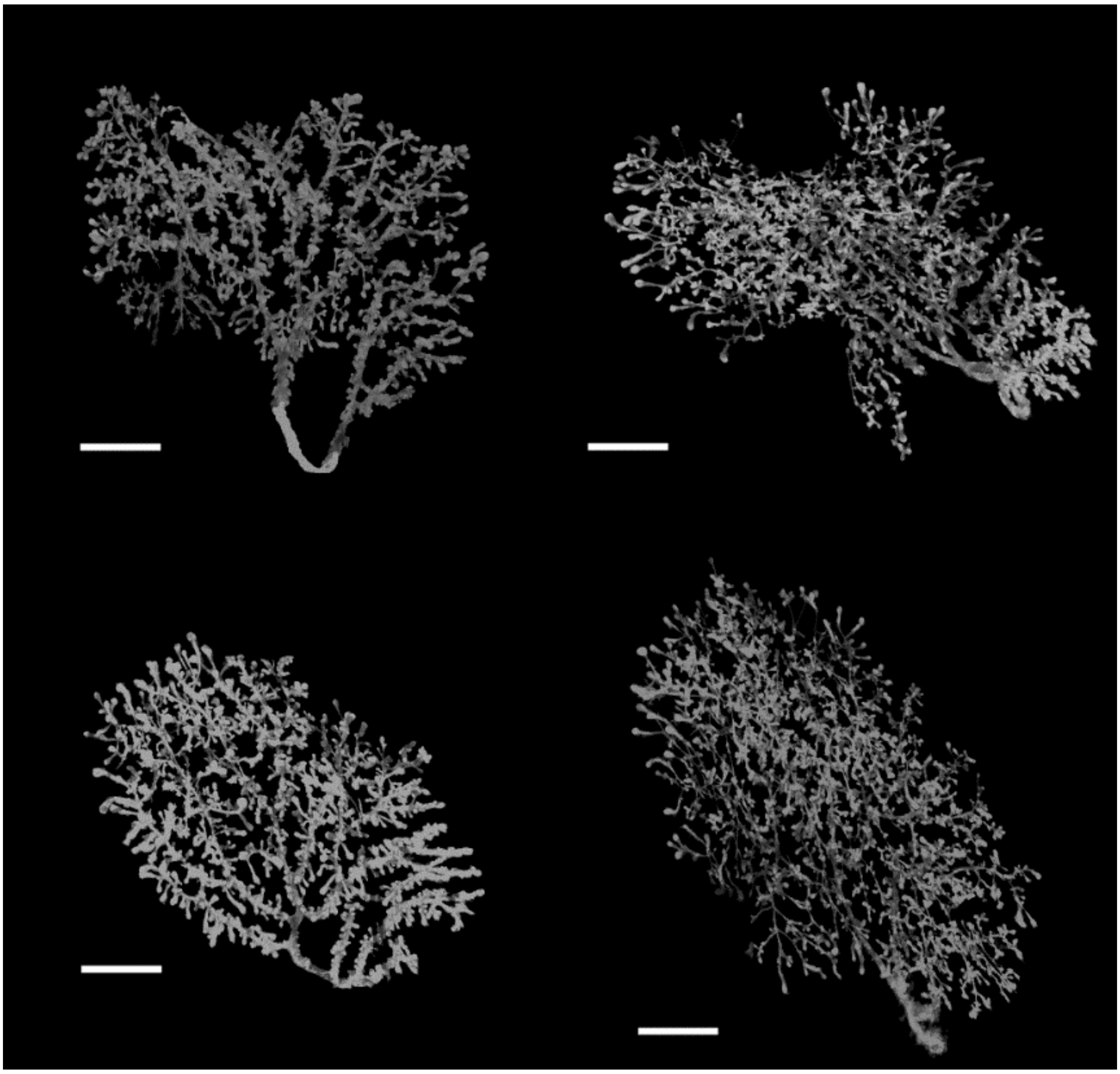

**Figure S4.** Illustration of PCA-identified clusters significantly related to treatment. Top, samples typical of significant clusters, left 250BPA, right 25BPA. Bottom, paragons of significant clusters, left 250BPA, right 25BPA. The 250BPA glands are less developed, with less space-filling branches than the 25BPA. Scale bar=2mm.

**90 days and 6 months:** Following the same method as for PND21C, we performed clustering on principal components for animals in the older groups. Then, we looked at significant correlations between clusters and treatments and report p-values in Table S3.

Overall, three conditions stand out since they are detected in the largest number of datasets: EE2, 25BPA and 250BPA Nevertheless, we are not satisfied with the stability of the method, for example between dataset with or without EE2.

**Table S3**. Results of clustering on principal components in PND 90 and 6 months animals.

| Dataset | 0.5EE2 | 0.05EE2 | Control | 2.5BPA | 25BPA | 250BPA | 2500BPA | 25000BPA |
|---|---|---|---|---|---|---|---|---|
| PND21C | 5.5e-6, 0.014 | | | | | 0.0068 | | |
| PND90CD | | | | 0.027 | | | | |
| PND90SD | 0.016 | | | | 0.030 | | | |
| 6MCD | 0.0015 | | | | 0.031 | 0.0098 | | 0.043 |
| 6MSD | 0.0019, 0.015 | | 0.0056 | | | 0.00064 | | 0.020 |
| PND21C | NA | NA | | | 0.028 | 0.0034 | | |
| PND90CD | NA | NA | | | 0.016 | | | |
| PND90SD | NA | NA | | | 0.0075 | | | |
| 6MCD | NA | NA | | | | 0.0089 | | |
| 6MSD | NA | NA | | | | | | |

Note: The presence of more than one p-values means that a condition is correlated with several clusters. Number of animal per group and per dataset n=8-10.

### Global analysis

Table S4 provides systematic results of the permutation test in our global analysis of 25BPA - 250BPA as a breaking point. In almost all cases, we see that $X_{observed}$ is significantly higher than the mean of the simulations. The only cases where the results are less significant is when the criteria are too restrictive, for $p_{thr} \leq 0.3$ or for high values of $r_{thr}$ (note that for $r_{thr}$ higher than 2, the distribution of $X_{sim}$ convergence is less clear with 10000 iterations).

**Table S4.** Comparison of $X_{observed}$ and the results of the permutation test for different values of the criteria A and B with datasets PND90CD, PND90SD, 6MCD and 6MSD.

| Criterion | $X_{observed}$ | 95 % of $X_{sim}$< | 99 % of $X_{sim}$< | 99.5% of $X_{sim}$< | $P_{estimated}$ |
|---|---|---|---|---|---|
| A(1)=no threshold | 1.43 | 1.08 | 1.24 | 1.29 | 0.00085*** |
| A(1.05) | 1.43 | 1.08 | 1.24 | 1.30 | 0.00091*** |
| A(1.1) | 1.49 | 1.09 | 1.26 | *1.32* | 0.00064*** |
| A(1.2) | 1.67 | 1.13 | 1.31 | 1.38 | 0.00016*** |
| A(1.3) | 1.66 | 1.15 | 1.34 | 1.41 | 0.00029*** |
| A(1.4) | 1.73 | 1.16 | 1.36 | 1.44 | 0.00026*** |
| A(1.5) | 1.93 | 1.17 | 1.32 | 1.45 | 2.2e-05*** |
| A(1.75) | 1.83 | 1.21 | 1.44 | 1.53 | 0.00029*** |
| A(2) | 1.55 | *1.23* | 1.47 | 1.56 | 0.0055** |
| A(2.5) | 1.29 | 1.27 | 1.52 | 1.61 | 0.044* |
| B(1)=no threshold | 1.24 | 1.11 | 1.27 | 1.33 | 0.014* |
| B(0.75) | 1.25 | 1.10 | 1.27 | 1.33 | 0.012* |
| B(0.6) | 1.29 | 1.11 | 1.28 | 1.34 | 0.0086** |
| B(0.5) | 1.37 | 1.12 | 1.29 | 1.35 | 0.0038*** |
| B(0.4) | 1.36 | 1.14 | 1.32 | 1.39 | 0.0066** |
| B(0.3) | 1.16 | *1.20* | 1.40 | 1.48 | 0.061 |
| B(0.2) | 1.31 | *1.26* | 1.50 | 1.59 | 0.037* |
| B(0.1) | 1.41 | 1.47 | 1.81 | 1.95 | 0.060 |

**Note:** The results show that a remarkable effect occurs between 25BPA and 250BPA. Number of animals per group n=8-10. Number of groups: 6.

**Simulations:** We assessed type 1 and type 2 error rates by simulations. To this end we used distributions mimicking our data. First we used Gaussian distributions with standard deviation 1 and mean 0, a/2,a,0,a/2,a where a is a parameter which determines the magnitude of the breaking point (the standard deviation being 1, a=a/SD). In our experimental data we have 4 sets

(PND90CD, PND90SD, 6MCD,6MSD) with 6 “doses” and roughly 10 “animals” per group. In each of our simulations, we used this format to generate data 10000 times (1000 times in the case of type 2 errors. To take into account a key feature of our data, we also tested simulations where the different Gaussian variables are correlated according to the correlations of the N first variables of the 6MCD data.
For computational reasons, we estimate the statistic of X only in the first of the 10000 generated datasets, which we expect, leads to a moderate increase of errors rates.

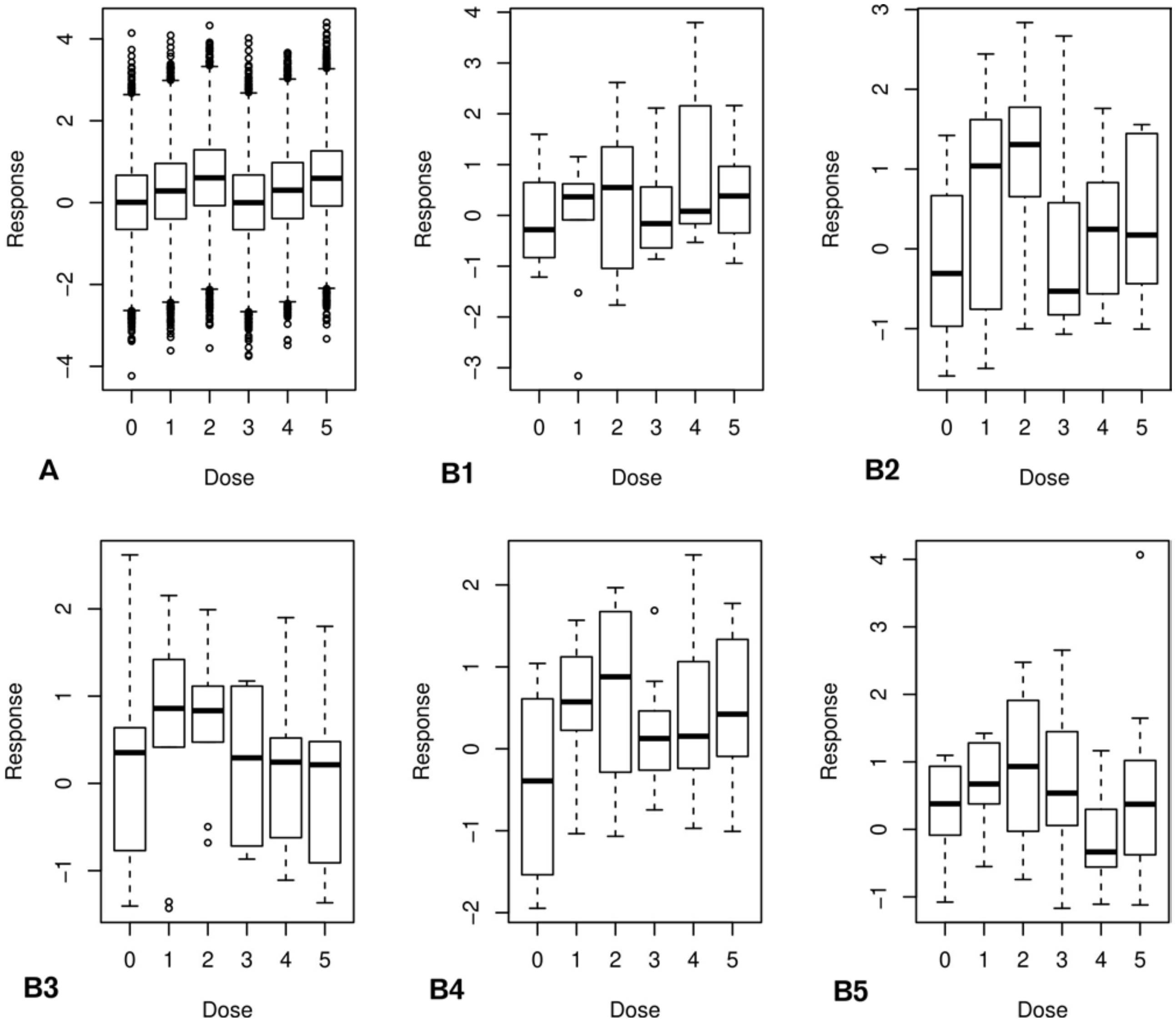


**Figure S5.** *Simulated dose response with a=0.6 (without correlations).* A: We represent a simulation with 10000 “animals” per group to show the shape of our simulated distribution. B: several iterations of our simulated distribution with the usual 10 animal per group.

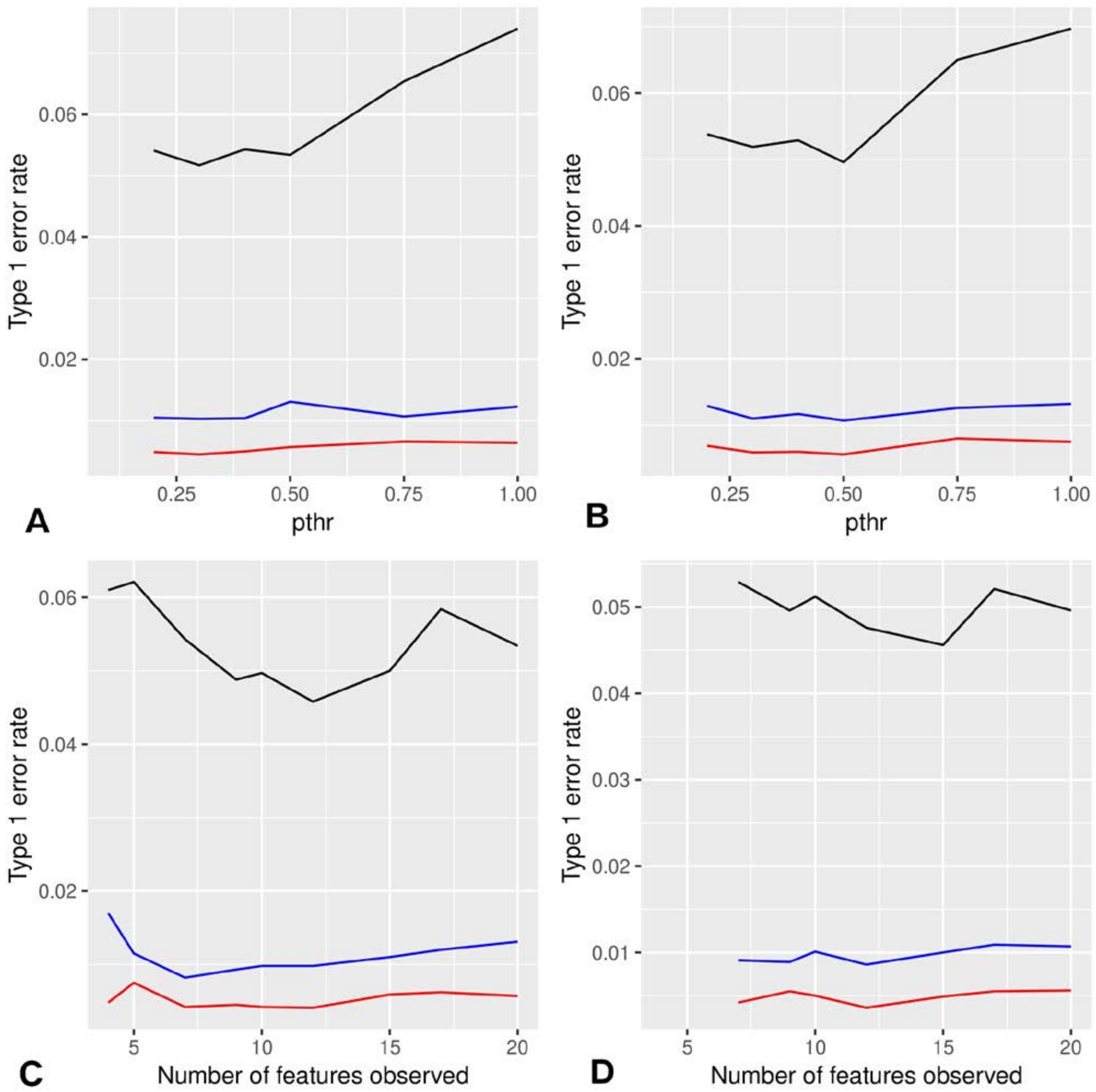


**Figure S6.** *Estimated type 1 error rates (0.05 in black, 0.01 in blue, 0.005 in red).* A, C; the different variables are not correlated. B,D: the different variables are correlated according to our data. A, B: Type 1 error rate as a function of the threshold for criterion B($p_{thr}$), with 20 variables. The error rate remains close to the target with a loss of precision for $p_{thr}$ =0.75 or 1 which is consistent with the idea that these values are not restrictive enough. The drift remains moderate and does not change in B with correlated variables. C, D: Type 1 error rate as a function of the number of features observed for $p_{thr}$ =0.5. The error rates remain consistent with moderate variations.

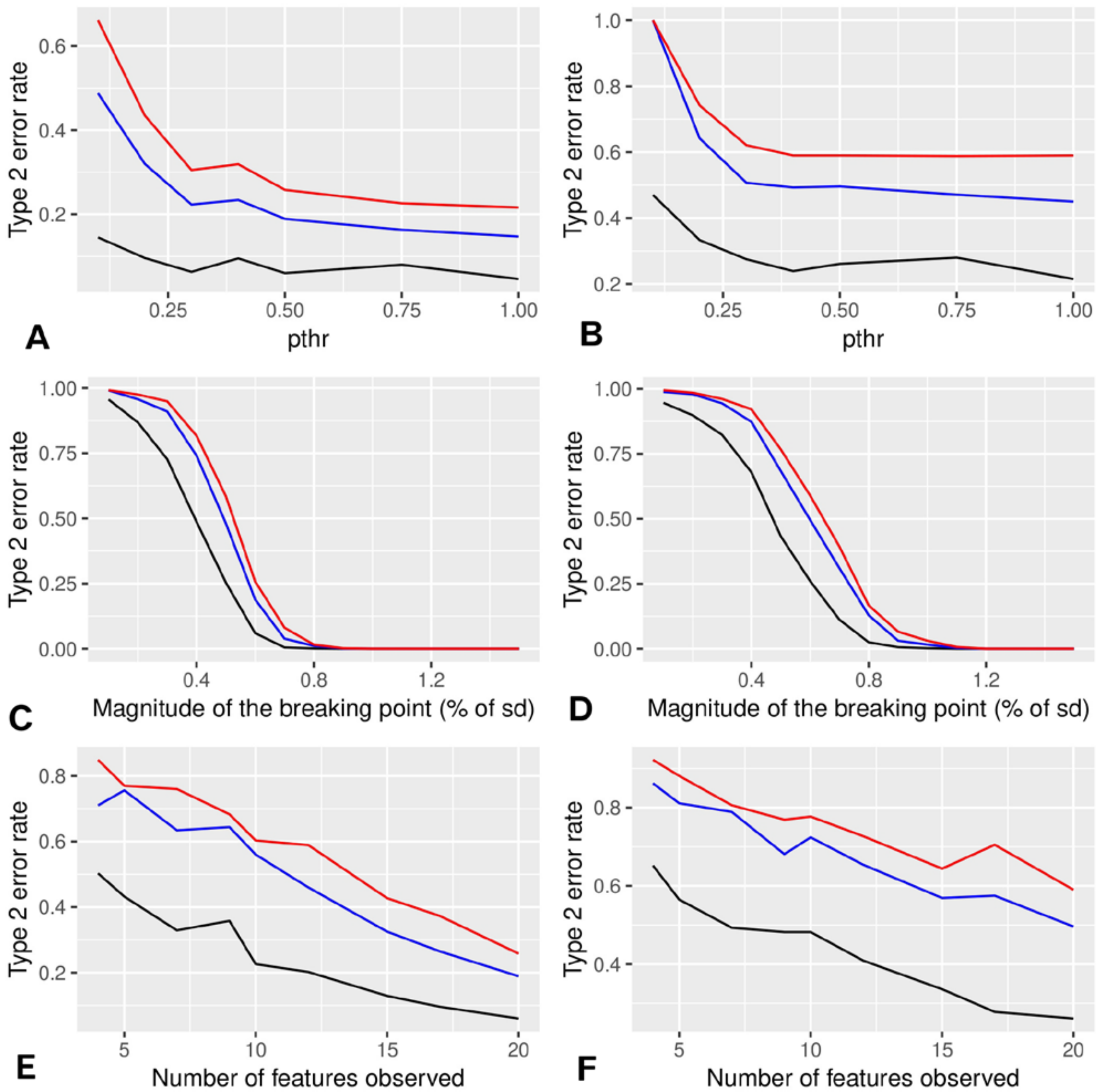


**Figure S7.** *Estimated type 2 error rates (0.05 in black, 0.01 in blue, 0.005 in red).* A, C, E: the different variables are not correlated. B,D,F: the different variables are correlated according to our data. A, B: type 2 error rate as a function of the threshold for criterion B($p_{thr}$), with 20 variables and a=0.6 which is an intermediate value. Higher values of $p_{thr}$ lead to decreased type 2 error rates. Overall, the error rate with correlated variables is higher which is logical since the number of degree of freedom of the data is lower. C,D: type 2 error rate as a function of a with N=20. The error rate becomes very low starting with a=0.6 ( C ) and a=0.8 ( D ). E, F: type 2 error rate as a function of the number N of variables describing each individual with a=0.6, and $p_{thr}$=0.5. The higher N is, the lower the error rate is which is logical since the number of degree of freedom of the data is increased.

#### Regression analysis

Figure S8 displays several graphical tests to assess the quality of the regressions performed in the main text as provided by the lm method in cran R. The first graph, Residual versus Fitted assesses the presence of a pattern not taken into account by the model and homoscedasticity (i.e., that variance is constant). The second graph assesses the normality of residuals. The third graph is used to assess homoscedasticity. The fourth graph aims at assessing the presence of outliers. Last, the fifth graph displays a box plot of the data and the fitted model.

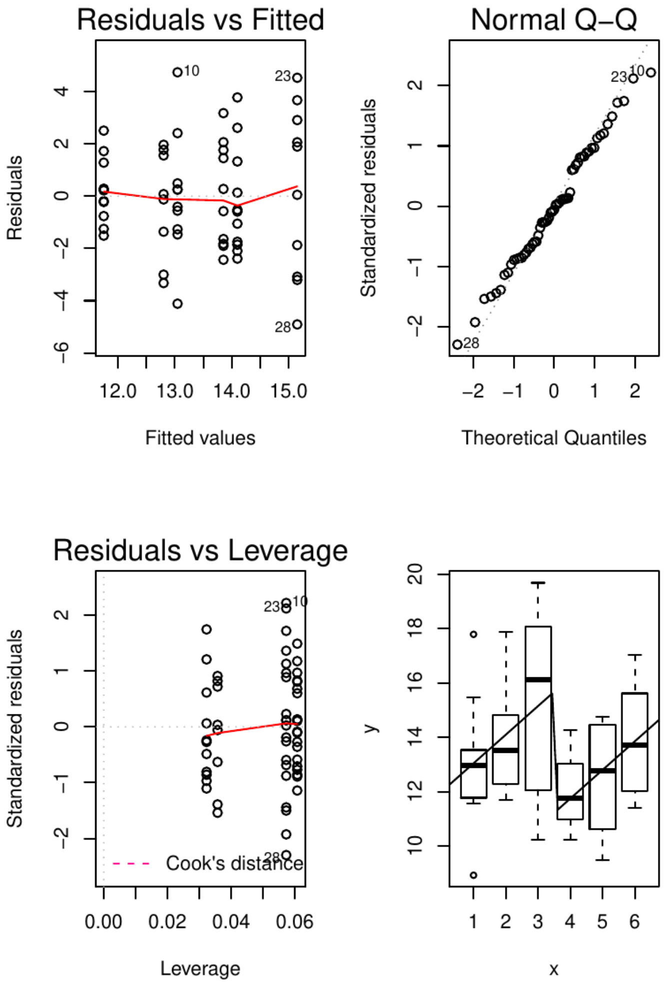


**Figure S8A** sd width 3D in PND21C

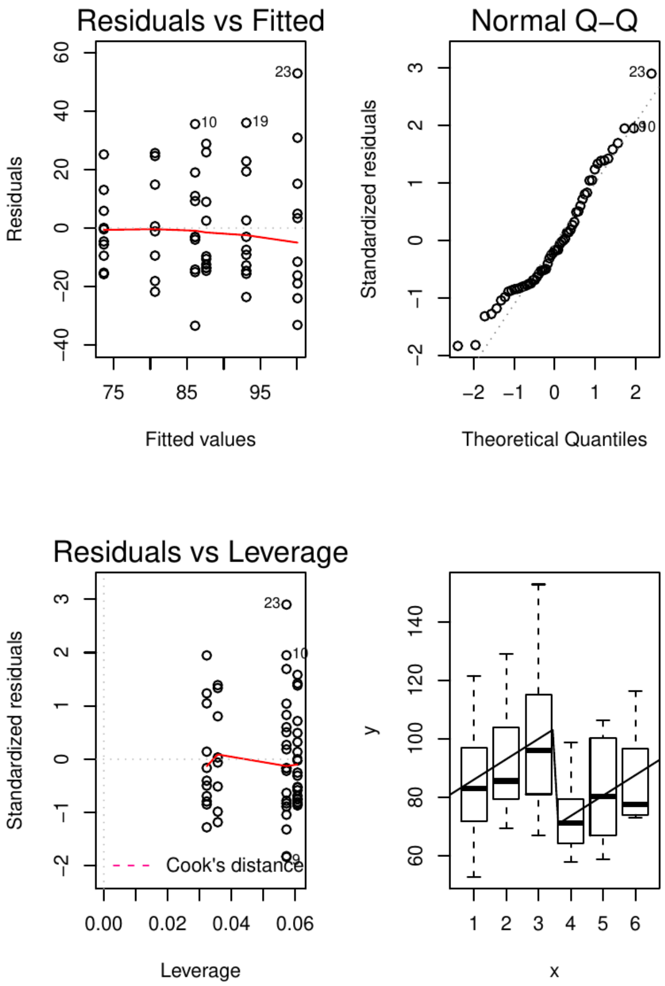


**FigureS8B** Thickness in PND21C

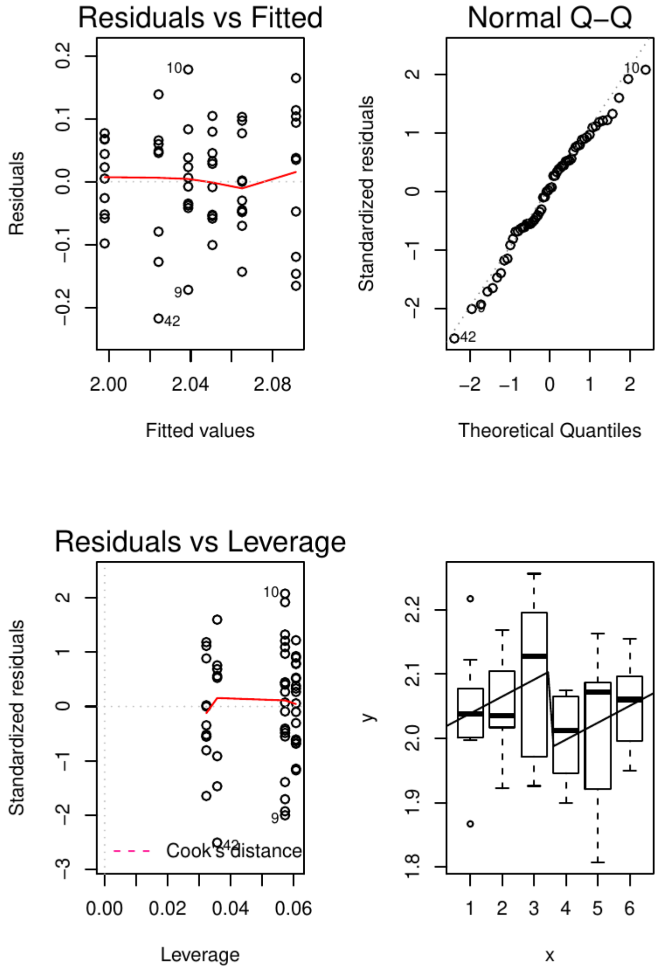


**Figure S8C** Fractal dimension in 3D in PND21C

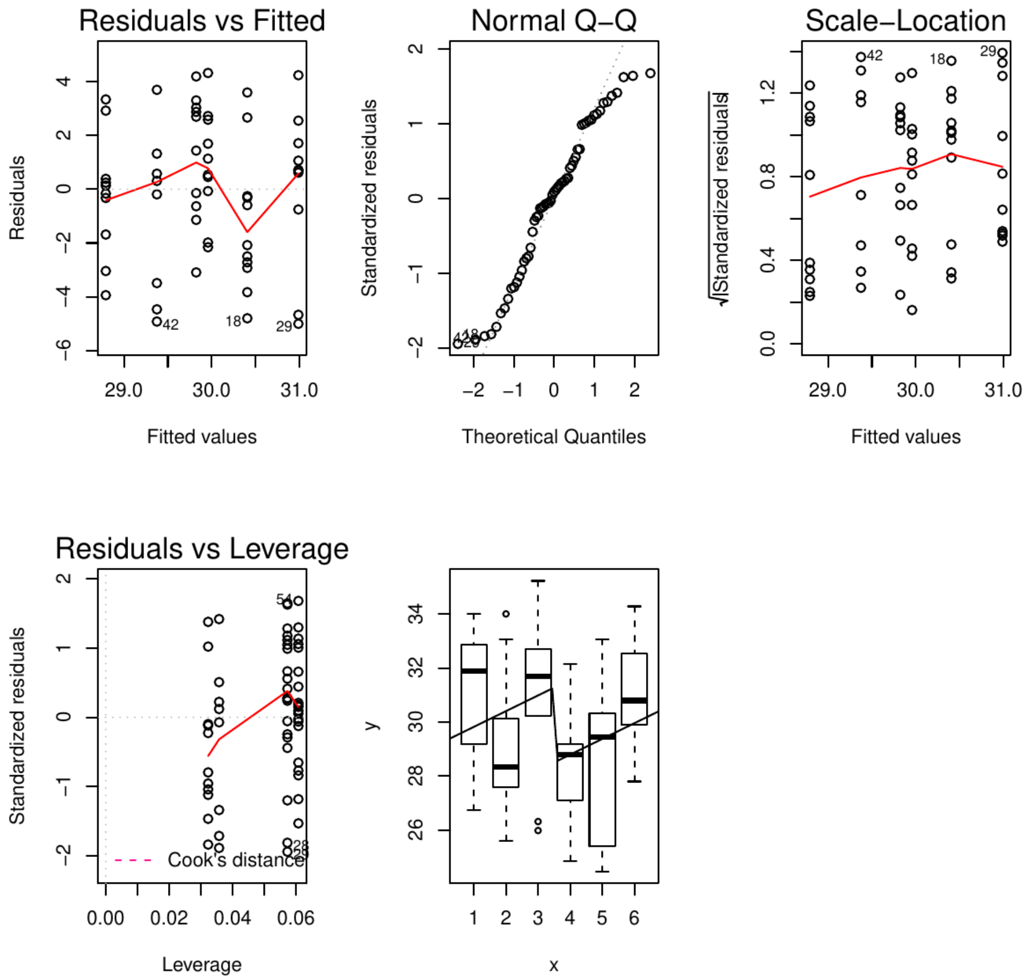


**Figure S8D** Angle between beginning and end in PND21C. Here, the pattern does not fit the model completly.

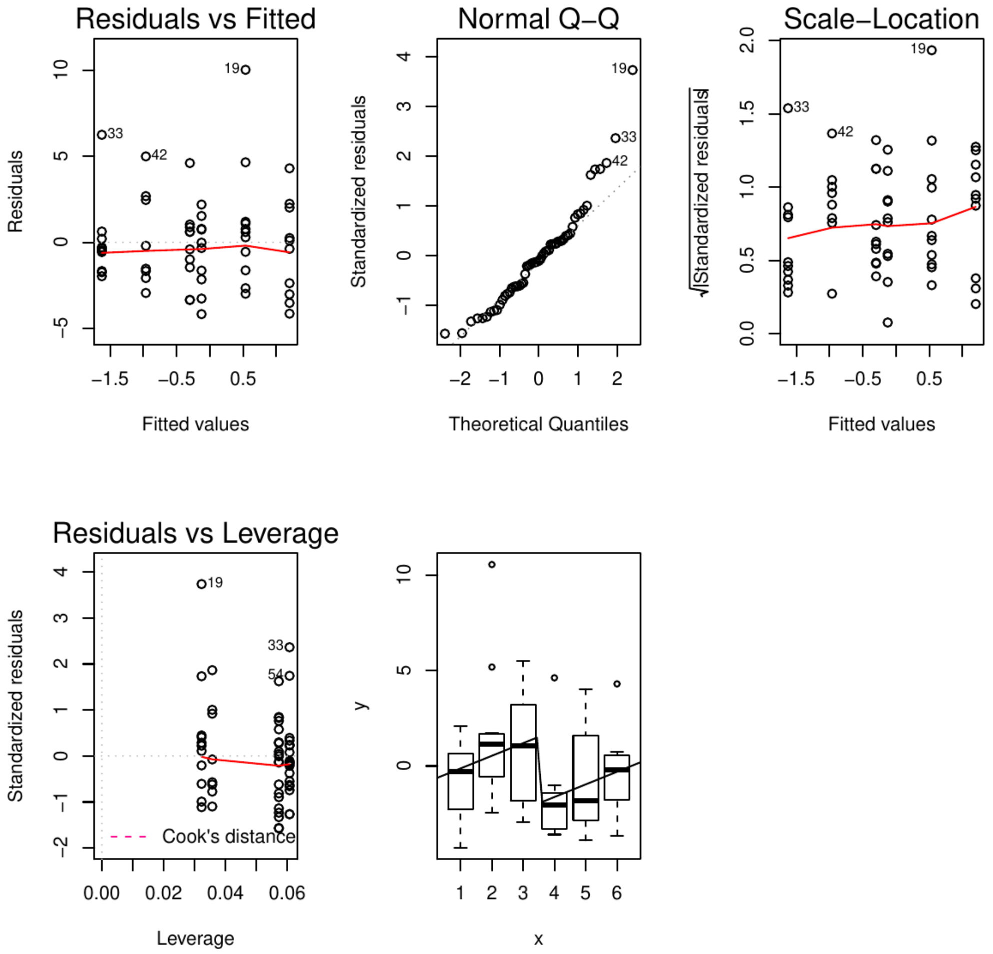


**Figure S8E** Dim.3 resulting from PCA in PND21C

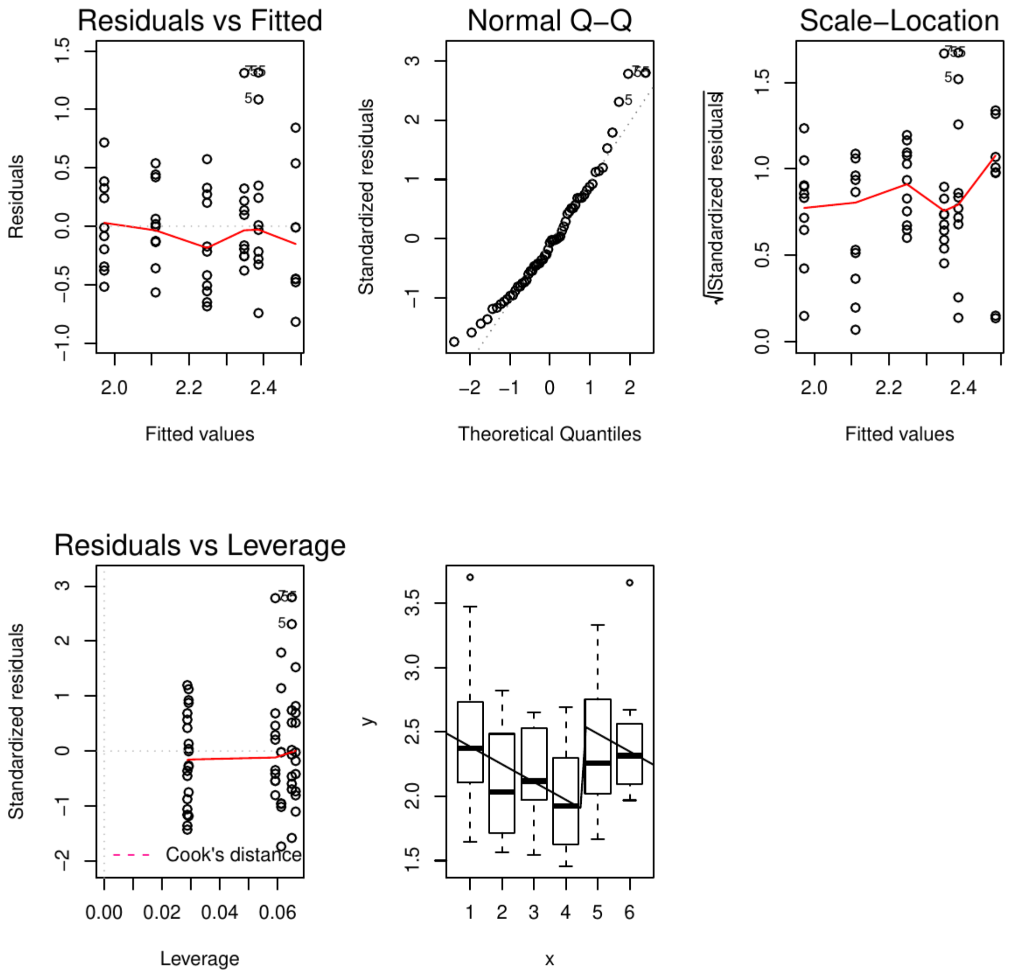


**Figure S8F** Aspect ratio in PND21C

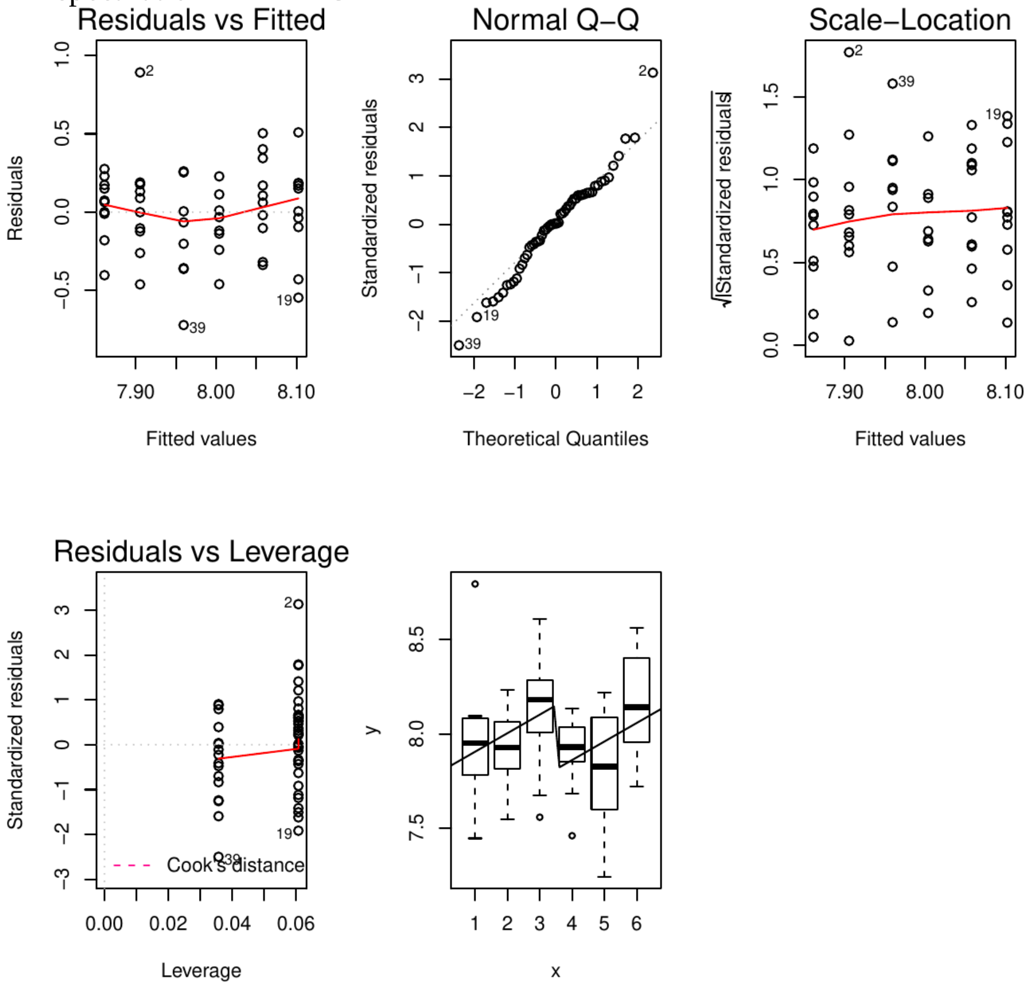


**Figure S8G** Mammary gland weight in PND90SD

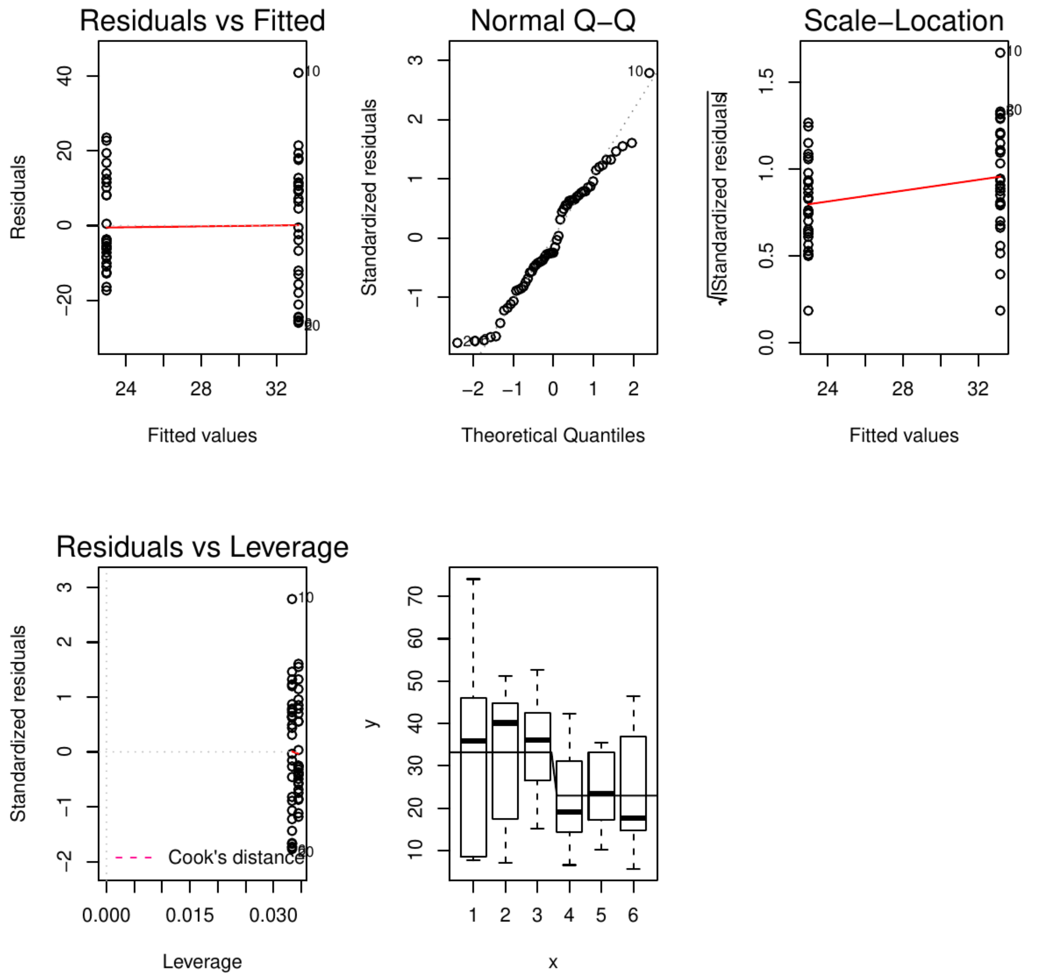


**Figure S8H** Density in area 3 in PND90CD

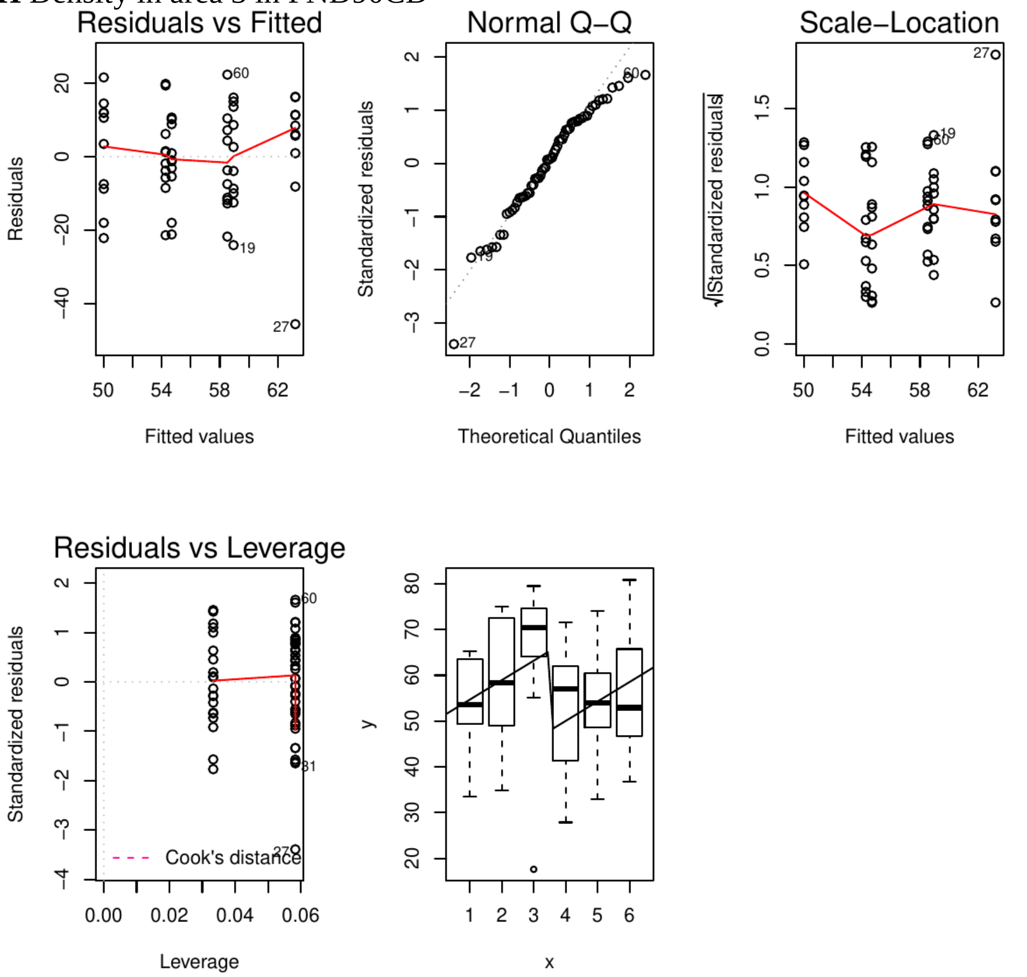


**Figure S8I** Density in area 3 in 6MCD

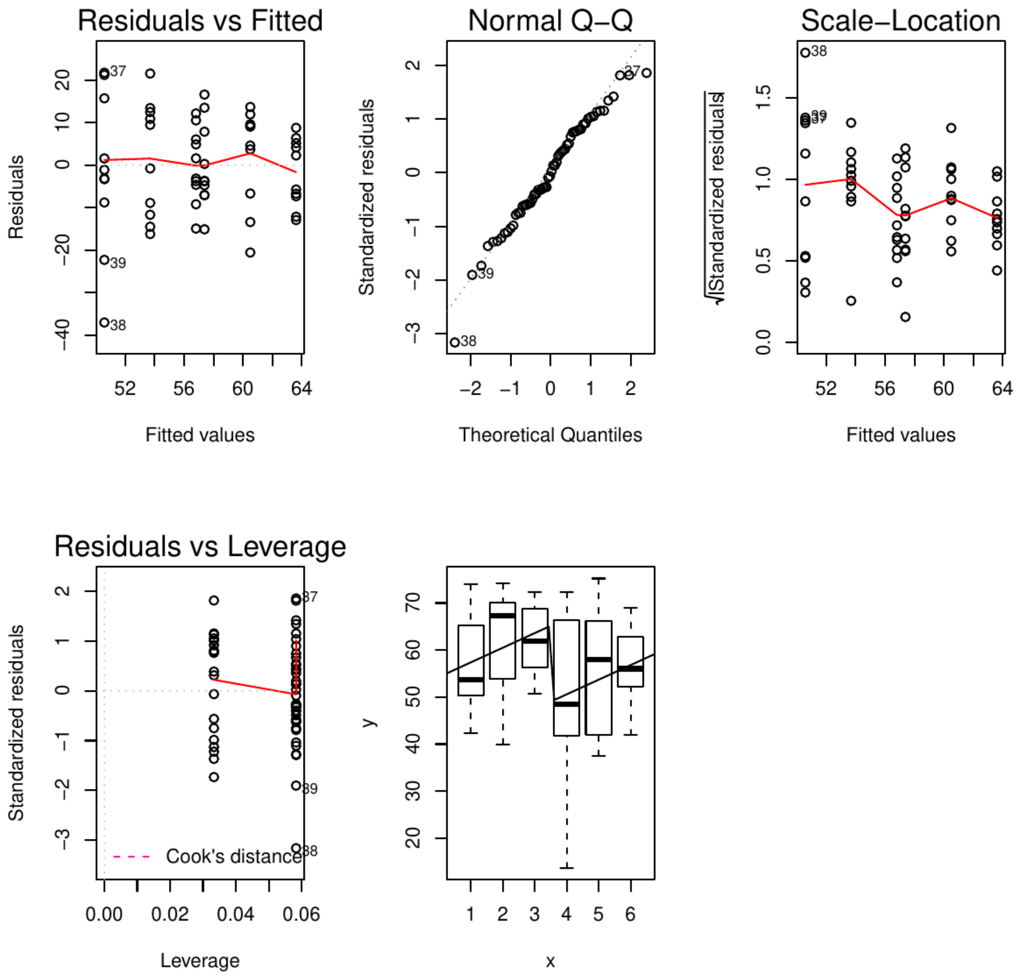


**Figure S8J** Density in area 3 in 6MSD

**Table S5.** Incidence of benign and malignant lesions/tumors identified from A) PND 90 and B) 6 month mammary glands following either continuous or stop-dose exposures across all treatment groups.

A

**PND 90 Continuous Dose (PND90CD)**

| Treatment | Animals (n) | Lobular Hyperplasia | Fibroadenoma | Periductular Fibrosis (± lymphocytic infiltration) | Ductal epithelial necrosis with inflammatory infiltrate | DCIS |
|---|---|---|---|---|---|---|
| Control | 10 | 0 | 0 | 0 | 0 | 0 |
| 2.5BPA | 9 | 0 | 0 | 0 | 0 | 0 |
| 25BPA | 10 | 0 | 0 | 1 | 0 | 0 |
| 250BPA | 9 | 0 | 0 | 0 | 0 | 0 |
| 2500BPA | 9 | 0 | 0 | 0 | 0 | 0 |
| 25000BPA | 10 | 0 | 0 | 1 | 1 | 0 |
| 0.05EE2 | 10 | 0 | 0 | 0 | 0 | 0 |
| 0.5EE2 | 10 | 0 | 0 | 0 | 0 | 1 |

**PND 90 Stop Dose (PND90SD)**

| Treatment | Animals (n) | Lobular Hyperplasia | Fibroadenoma | Periductular Fibrosis (± lymphocytic infiltration) | Ductal epithelial necrosis with inflammatory infiltrate | DCIS |
|---|---|---|---|---|---|---|
| Control | 10 | 0 | 0 | 0 | 0 | 0 |
| 2.5BPA | 8 | 0 | 0 | 0 | 0 | 0 |
| 25BPA | 10 | 0 | 0 | 1 | 0 | 0 |
| 250BPA | 10 | 0 | 0 | 0 | 0 | 2 |
| 2500BPA | 8 | 0 | 0 | 0 | 0 | 0 |
| 25000BPA | 10 | 0 | 0 | 0 | 0 | 0 |
| 0.05EE2 | 9 | 1 | 1 | 0 | 0 | 0 |
| 0.5EE2 | 10 | 0 | 0 | 0 | 0 | 0 |

B

**6 Month Continuous Dose (6MCD)**

| Treatment | Animals (n) | Lobulo/Ductular-alveolar dilatation (± secretions) | Periductular Fibrosis (± lymphocytic infiltration) | Fibroadenoma | Adenoma | Adenocarcinoma (±cyst) |
|---|---|---|---|---|---|---|
| Control | 10 | 0 | 1 | 0 | 0 | 0 |
| 2.5BPA | 10 | 0 | 0 | 1 | 0 | 0 |
| 25BPA | 10 | 0 | 0 | 1 | 0 | 0 |
| 250BPA | 10 | 0 | 0 | 0 | 0 | 0 |
| 2500BPA | 10 | 0 | 0 | 0 | 0 | 0 |
| 25000BPA | 10 | 0 | 0 | 0 | 0 | 0 |
| 0.05EE2 | 10 | 0 | 0 | 0 | 0 | 0 |
| 0.5EE2 | 10 | 4 | 0 | 2 | 3 | 1 |

**6 Month Stop Dose (6MSD)**

| Treatment | Animals (n) | Lobulo/Ductular-alveolar dilatation (± secretions) | Periductular Fibrosis (± lymphocytic infiltration) | Fibroadenoma | Adenoma | Adenocarcinoma (±cyst) |
|---|---|---|---|---|---|---|
| Control | 10 | 0 | 0 | 0 | 0 | 0 |
| 2.5BPA | 10 | 1 | 0 | 0 | 0 | 0 |
| 25BPA | 10 | 0 | 0 | 0 | 0 | 0 |
| 250BPA | 10 | 0 | 0 | 0 | 0 | 0 |
| 2500BPA | 10 | 0 | 0 | 0 | 0 | 0 |
| 25000BPA | 10 | 0 | 1 | 0 | 0 | 0 |
| 0.05EE2 | 10 | 0 | 0 | 0 | 0 | 0 |
| 0.5EE2 | 10 | 4 | 1 | 1 | 1 | 2 |
